# Sequence features and dynamics of subcellular translation revealed by APEX-Ribo-Seq

**DOI:** 10.1101/2025.05.26.656194

**Authors:** Kotaro Tomuro, Shiho Makino, Takanori Akaiwa, Akane Harada, Kenta Kobayashi, Nobuyuki Shiina, Takashi Fukaya, Kyogo Kawaguchi, Yuichi Shichino, Shintaro Iwasaki

## Abstract

Local translation at specific subcellular regions is proposed to define appropriate protein destinations and functions. However, our understanding of local translation still remains far from complete due to a lack of versatile analytical tools. Here, we developed a new method, termed “APEX-Ribo-Seq”, that integrates ribosome profiling (Ribo-Seq) with APEX2-based proximity labeling. We mapped local translation across 14 distinct organelles and identified over 3,000 genes undergoing localized translation, encoding components of complexes that function at these specific sites. RNA language-model predictors captured compartment-associated information beyond protein sequence, and model-derived attribution nominated sequence-sensitive transcript regions for mechanistic follow-up. Gene-set-level RBP enrichment and knockdown experiments separately identified RBPs associated with compartment-specific translation. The utility of APEX-Ribo-Seq was demonstrated in primary neurons, revealing compartment-specific translatomes in dendrites, axons, and presynapses. Extending this approach *in vivo*, we applied APEX-Ribo-Seq to germ granules in *Drosophila* embryos, profiling the temporal transition from translationally silent mRNA storage to localized translation during early embryogenesis. Our results provide a comprehensive view of the spatiotemporal translatome and its regulation.

## Introduction

Spatiotemporal regulation of protein synthesis is a phenomenon that is broadly conserved from bacteria to humans, allowing fine-tuning of gene expression and cellular functions ^1,2^. Local translation offers significant advantages for maintaining proteome integrity within cells that are densely packed with structurally distinct organelles. For example, spatially restricted protein synthesis supports protein maturation via cotranslational translocation into the endoplasmic reticulum (ER) ^3,4^ and *in situ* assembly of protein complexes, as observed for the nuclear pore complex ^5^. Spatial restriction of protein synthesis also ensures high concentrations of specific proteins in discrete subcellular compartments, such as actin at the leading edge in migrating cells ^6^ and axial patterning proteins during early embryonic development ^7–10^. Moreover, local protein synthesis is highly advantageous in polarized cells such as neurons; these cells transport specific mRNAs to dendrites and axons and translate them locally, regulating activity-dependent synaptic plasticity, axon guidance, and memory formation ^11,12^.

Recent studies have suggested that local translation is more widespread than previously appreciated. In addition to the ER ^3,4^ and mitochondria ^13^, a variety of membranous organelles have been reported to exhibit local translation, such as endosomes/lysosomes ^14–18^, the Golgi apparatus ^19^, nuclear pore complexes ^5,20^, and peroxisomes ^21^. Cytoskeleton-associated structures such as centrosomes ^19,22–26^, spindles ^24,27,28^, cilia ^29–31^, midbodies ^32,33^, and cell protrusions ^34–37^ also serve as sites of translation. Furthermore, increasing evidence suggests active translation within ribonucleoprotein (RNP) granules, which are biomolecular condensates composed of RNAs and proteins ^10,38–44^. These findings imply that translation may occur in subcellular regions beyond those currently characterized; however, the global landscape of local translation remains unclear.

Local translation has traditionally been inferred from the subcellular localization of mRNAs; however, the presence of mRNAs is not always coupled with active translation ^34,35,45^, highlighting the necessity for direct observations of translation. Imaging-based approaches, such as nascent chain tracking in living cells ^46–50^, offer high spatial resolution but suffer from limited scalability and throughput. More recently, RIBOmap has enabled the visualization of thousands of ribosome-bound mRNAs via DNA amplicons and *in situ* sequencing ^51^; however, this approach requires predefined target genes for probe design.

In contrast, deep sequencing-based approaches allow higher throughput and less biased analysis. Ribosome profiling (Ribo-Seq), a comprehensive method for measuring cellular translation through RNase footprinting of ribosome-protected mRNAs ^52^, can be used to investigate local translation if combined with microdissection ^53–55^ or biochemical fractionation ^56,57^. However, purifying specific compartments without cross-contamination remains challenging. As an alternative, proximity-labeling of localized ribosomes using the biotin ligase BirA tethered to subcellular membranes and its biotin acceptor peptide (AviTag) fused to the ribosome protein has enabled more spatially restricted Ribo-Seq for the endoplasmic reticulum membrane (ERM) ^58–60^, outer mitochondrial membrane (OMM) ^61^, and peroxisomes ^21^. However, this strategy requires laborious optimization, including biotin depletion from culture media and ribosomal protein tagging. While the use of light-controllable split-BirA ^62^ or the promiscuous biotin ligase TurboID ^63^ has addressed some of the limitations, these methods still detect only ribosome-bound mRNAs and thus cannot accurately differentiate translation activity from RNA recruitment. Therefore, a more versatile and unbiased methodology for the genome-wide measurement of local translation has long been awaited.

Here, we present APEX-Ribo-Seq, a newly developed method that integrates APEX2-based proximity labeling, which targets both proteins and RNAs and is widely used in spatial proteomics and transcriptomics ^64–71^, with ultralow-input Ribo-Seq ^72^. This setup enables robust, genome-wide measurement of local translation at high spatial resolution. Importantly, simultaneous APEX-RNA-Seq ^64,65^ allowed us to dissect the effect of RNA localization and on-site translational control. To demonstrate the versatility of APEX-Ribo-Seq, we applied our technique across 14 distinct subcellular compartments—the ERM, OMM, peroxisome, *cis*-Golgi, *trans*-Golgi, lysosome, endosome, early endosome, centrosome, plasma membrane, cell junction, intermediate filaments, outer nuclear membrane, and clathrin-coated vesicles—and established a comprehensive subcellular translation atlas in human cells. Integration with machine learning and RNA language models captured RNA-sequence information associated with compartment-specific translation beyond that available from protein sequence alone and nominated prediction-associated transcript regions for mechanistic follow-up. Gene-set-level RBP enrichment and knockdown experiments provided separate functional evidence for selected RBPs. Beyond regular human tissue culture, we extended our analysis to mouse primary neurons for compartment-specific translatomes in dendrites, axons, and presynapses. Finally, we expanded APEX-Ribo-Seq to an *in vivo* method for intact fly embryos, targeting germ granules, membraneless RNA-protein condensates. This allowed us to monitor the temporal dynamics of the local translatome during early embryogenesis, shifting from translationally silent mRNA storage. Together, these findings highlight APEX-Ribo-Seq as a powerful and broadly applicable platform for investigating spatiotemporal translation regulation in diverse biological contexts.

## Results and Discussion

### APEX2-mediated proximity labeling of ribosome mRNA complexes

To develop a method to survey local translation, we sought to combine APEX2-based proximity labeling with ribosome profiling. In the presence of H_2_O_2_, the APEX2 protein—an engineered peroxidase ^67^—generates short-lived radicals from phenol and covalently conjugates phenol to nearby RNAs and proteins ^64,65,67,73–75^. Thus, the fusion of biotin to phenol provides a tag for subsequent purification using streptavidin beads. Given that APEX2 tagging of organelle markers allows a survey of the local proteome ^66,69,73,76,77^ and transcriptome ^64,65,74^, we reasoned that this technique could be used to label local ribosome mRNA complexes for subsequent Ribo-Seq experiments (Figure 1A).

**Figure 1.**
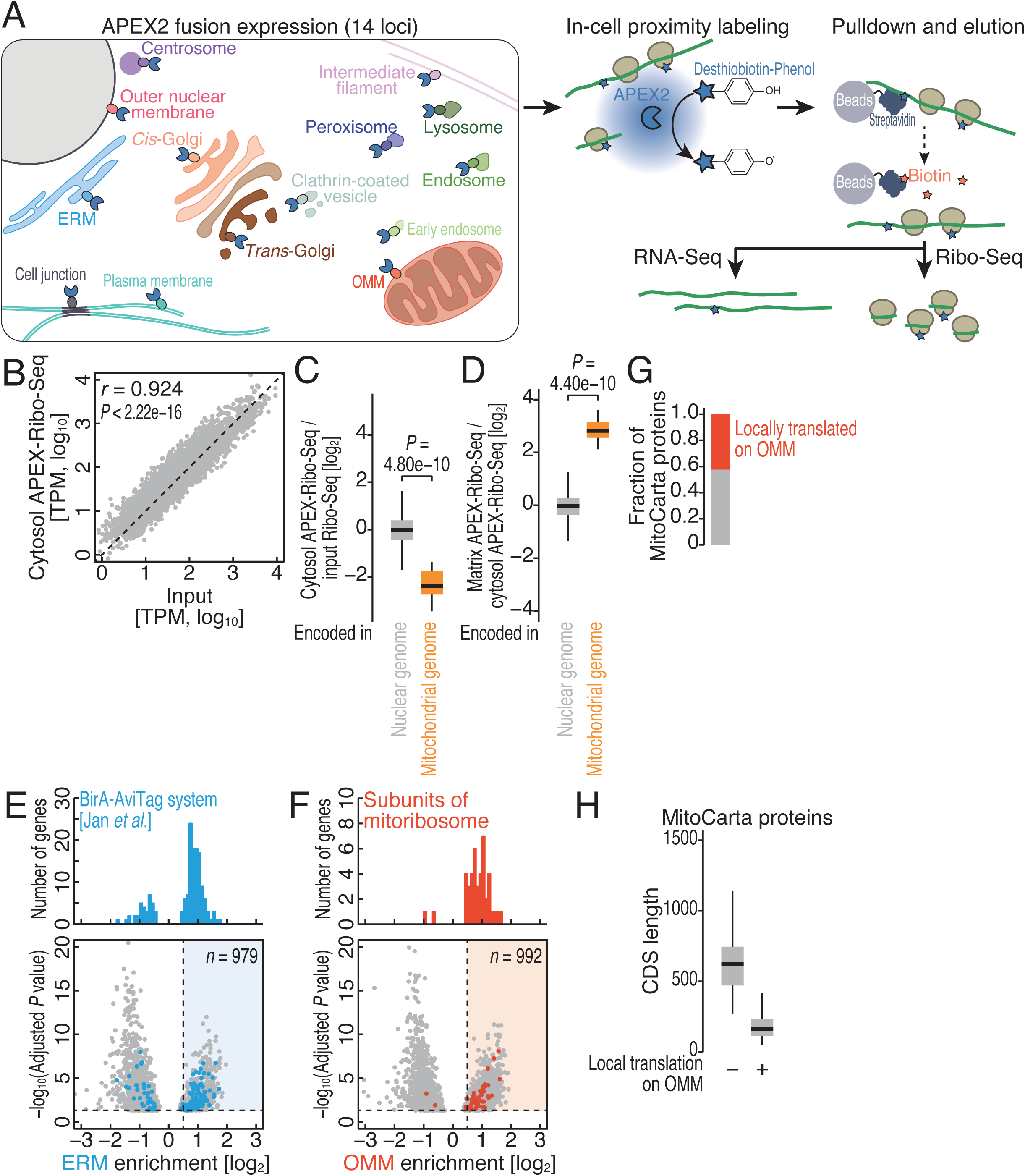
Benchmarking of APEX-Ribo-Seq. (A) Schematic of APEX-Ribo-Seq and APEX-RNA-Seq. (B) Scatter plot of the ribosome footprints from each mRNA in standard Ribo-Seq (input) and cytosol APEX-Ribo-Seq. *P* values were calculated with Pearson’s product-moment correlation test (two-sided). (C) Box plots of cytosol APEX-Ribo-Seq enrichment over input Ribo-Seq for mRNAs encoded in the nuclear genome and mitochondrial genome. *P* values were calculated with the Mann Whitney *U* test (two-tailed). (D) Box plots of matrix APEX-Ribo-Seq enrichment over input Ribo-Seq for mRNAs encoded in the nuclear genome and mitochondrial genome. *P* values were calculated with the Mann Whitney *U* test (two-tailed). (E-F) Volcano plots showing organelle enrichment determined by APEX-Ribo-Seq (Figure S2E). mRNAs with an adjusted *P* value of 0.05 or lower are shown. The colored areas represent mRNAs that met the fold change threshold defined by ROC analysis (Figure S2F). On top, histograms for the indicated groups of mRNAs depicted within the volcano plots are shown. (G) Fraction of mRNAs locally translated on the OMM among all mitochondria-targeting proteins (defined by MitoCarta 3.0) ^97^. (H) Box plots of the CDS length for the indicated groups of mRNAs. TPM, transcripts per million; *r*, Pearson’s correlation coefficient. Box plots show the median (centerline), upper/lower quartiles (box limits), and 1.5× interquartile range (whiskers). See also Figures S1 and S2 and Tables S1 and S4.

Here, we optimized the method, termed APEX-Ribo-Seq. Although the biotin streptavidin interaction is strong, contaminating proteins and RNAs that remain after purification via streptavidin beads may hamper specificity. To overcome this issue, we substituted the conventional biotin-phenol with desthiobiotin-phenol, which can be utilized with APEX2 ^78–80^. Desthiobiotin has weaker, reversible binding to streptavidin (*K*_a_ = 10^12^ M^−1^), allowing competitive elution from the beads with the higher-affinity biotin (*K*_a_ = 10^15^ M^−1^) ^81^, reducing nonspecific contamination during purification (Figure 1A).

To evaluate the labeling and purification performance of this approach, we established HEK293 T-REx cells stably expressing a nuclear export signal (NES) tagged with APEX2 (APEX2-NES-EGFP) (Figure S1A); this protein should be uniformly diffused in the cytosol. As expected, colocalization of the protein (detected via enhanced green fluorescent protein [EGFP]) and desthiobiotinylated molecules was observed throughout the cytosol (Figure S1B). Desthiobiotinylated RNAs and proteins blotted on membranes were monitored in a desthiobiotin phenol- and H_2_O_2_-dependent manner, with detection via fluorophore-conjugated streptavidin (Figure S1C). The co-sedimentation of desthiobiotinylated RNAs and proteins with polysomes in a sucrose density gradient suggested the successful labeling of ribosomal complexes by the APEX2 system (Figure S1D-E). These ribosome–mRNA complexes were tethered to streptavidin beads, washed, and competitively eluted with free biotin (Figure 1A). However, this purification approach exhibited the drawback of low material recovery. To overcome this issue, we harnessed a Ribo-Seq derivative tailored for low input, termed T7 High-resolution Original RNA (Thor)-Ribo-Seq ^72^.

One of the general concerns associated with the APEX2-based method is the use of H_2_O_2_, which induces oxidative stress. Therefore, here, we used a short treatment (1 min) with subsequent quick quenching; critically, stress-response induction at the translational level was not detected under these experimental conditions (Figure S1F). Consistent with this, stress-induced phosphorylation of eIF2α, which leads to global translation shutoff ^82^, was minimal under immunostaining during 1-min short H_2_O_2_ treatment (Figure S1G-H), while longer incubation accumulated phosphorylated eIF2α.

We subsequently conducted cytosolic APEX-Ribo-Seq with APEX2-NES-EGFP for further optimization. Earlier APEX2-based techniques employed a high salt wash (1 M NaCl) to reduce contamination ^64,65^. However, under our conditions, a high salt concentration (Figure S1I) did not improve the data quality; rather, we found that the yield of ribosome footprints mapped to the genome (Figure S1J) was reduced relative to that obtained using the standard medium salt concentration for regular Ribo-Seq (150 mM NaCl) ^83,84^. Additionally, the mild salt condition resulted in a typical footprint length distribution (Figure S1K), marked 3-nt periodicity (Figure S1L-M), high reproducibility (Figure S1N), and high correspondence to the input standard Ribo-Seq (Figure 1B), indicating uniform recovery of cytosolic ribosomes. In contrast, a high salt wash caused high variability in quality between replicates, as represented by footprint length distribution (Figure S1K). As we focused on 29-nt footprints in the metagene plot (Figure S1L), the read yields were lower than those in the moderate salt condition. Thus, we used the medium salt concentration wash for the pulldown.

Given that mitochondrial ribosomes (mitoribosomes) in the matrix constitute a separate translation system, distinct from cytosolic ribosomes, the NES-fused APEX2 enzyme in the cytosol should not have access to mitoribosomes. Consistent with this expectation, mitoribosome footprints corresponding to all 13 coding sequences (CDSs) in the mitochondrial genome ^85,86^ were depleted in cytosol APEX-Ribo-Seq (Figure 1C). We subsequently conducted reciprocal experiments to capture mitochondrial translation in the matrix by using APEX2 fused to the mitochondrial targeting signal (MTS) (Figure S1A). As expected, the localization of MTS-APEX2 and the desthiobiotinylation reaction were restricted to the mitochondria (Figure S1O). Moreover, matrix APEX-Ribo-Seq exploiting MTS-APEX2 was enriched in mitoribosome footprints (Figure 1D). The 3-nt periodicity confirmed the recovery of mitoribosome footprints ^87–89^ (Figure S1P), indicating explicit targeting of mitochondrial translation by the APEX-Ribo-Seq matrix.

Thus, we employed these refined procedures for the downstream analyses.

### Benchmarking APEX-Ribo-Seq

To evaluate the performance of our approach for on-organelle translation, we focused on the ERM, which is a well-characterized site of local translation ^3,58,64,66,90,91^. We used APEX2 as a C-terminal tag for the CYP2C1 TM domain, which faces the cytosolic side of the ER ^64–66,68^ (Figure S1A); as expected, the tagged protein expression and desthiobiotinylation were localized to the ER (Figure S2A). ERM APEX-Ribo-Seq maintained the hallmarks of ribosome footprints, including read length distribution and strong 3-nt periodicity (Figure S2B-D).

Relative to the membrane-enclosed ER lumen or mitochondrial matrix, the cytosolic surfaces of organelles present intrinsic challenges for proximity labeling; the highly open and dynamic configuration may hinder efficient tagging by limited desthiobiotin diffusion within a short timeframe (1 min in our conditions), resulting in a relatively high background ^64,66^. Thus, we calculated “fold enrichment” for these sites carefully, considering 1) normalization by input lysate which may include the effects of H_2_O_2_ treatment while they were minimal in our hands (Figure S1F-H) and 2) the same APEX experiment setup for cytosolic free pool of ribosomes with APEX2-NES (Figure S2E); the latter point was not considered in BirA–AviTag-mediated system ^58,61^ due to its high labeling selectivity for the immediate vicinity of the organelle surface. Although simple statistical analysis could screen the subgroups of mRNAs, to add further stringency, we considered BirA–AviTag-mediated proximity-specific ribosome profiling data ^58^ as a reference and defined another fold enrichment threshold through receiver operating characteristic (ROC) analysis ^92^ (Figure S2F). Through this conservative definition, ultimately, we designated 979 genes that were locally translated on the ERM (Figure 1E and Table S1). This set of genes showed high correspondence with the secretome (see Methods for details), as well as with the set of genes identified by ERM-targeted Frac-Seq ^91^, APEX-RIP ^66^, and APEX-RNA-Seq ^64^ (Figure S2G). Search Tool for the Retrieval of Interacting Genes/Proteins (STRING) ^93^ also revealed the enrichment of ER-related genes (Figure S2H). Among the mRNAs identified as translated on the ERM (Table S1), *XRP1* and *CAST* were confirmed via single-molecule inexpensive fluorescent *in situ* hybridization (smiFISH) to localize to the ERM ^94^ (Figure S2I-J).

We found that APEX-Ribo-Seq captures a broader range of local translation than the BirA–AviTag approach. To characterize APEX-Ribo-Seq, we compared our data with ERM translation mapped by the BirA–AviTag system ^58^. Since this study, unlike ours, did not include a free cytosolic pulldown control, we calculated enrichment over input for both systems (Figure S2K). APEX-Ribo-Seq detected most (73.9%) mRNAs identified by BirA–AviTag (Figure S2K, “Detected in both”) and additionally captured many unique transcripts (Figure S2K, “Only detected in APEX”). This likely reflects the limited labeling radius of BirA–AviTag, which targets only 60S ribosomal proteins near the translocon but not 40S ^58^. In contrast, APEX labeling may extend further, encompassing ribosomes translating both membrane-translocated and peripherally associated proteins (Figure S2L, see below for details).

We subsequently focused on the translation around the OMM, another organelle associated with cotranslational membrane insertion. For this purpose, we fused APEX2 to the mitochondrial antiviral signaling (MAVS) C-terminal transmembrane (TM) domain, which anchors to the OMM ^95,96^ (Figure S1A); the N-terminal tag (APEX2-MAVS TM) faces the cytosolic side ^64^. The protein localization and desthiobiotinylation of the OMM were confirmed by immunostaining (Figure S2M). OMM APEX-Ribo-Seq recovered 3-nt periodic ribosome footprints (Figure S2N-O). Through enrichment analysis (Figure S2E) and threshold determination by ROC in ERM (Figure S2F), we identified 992 genes as OMM locally translated mRNAs (Figure 1F and Table S1); this set of genes aligned with the gene list produced by earlier APEX-RNA-Seq experiments ^64^ (Figure S2P). STRING analysis also indicated an overrepresentation of mitochondria-localized proteins (Figure S2Q). We observed that approximately 40% of the mitochondria-targeted proteins annotated in MitoCarta 3.0 ^97^ were found in the OMM dataset (Figure 1G), which is consistent with the results of studies using Transit-ID ^98^ and APEX-RNA-Seq ^64^. We also noted that the local translation detected on the OMM was biased toward mRNAs with short CDSs (Figure 1H). The examples of mRNAs locally translated on the OMM were those encoding mitoribosome subunits (Figure 1F), as was also reported in the Transit-ID study ^98^. To further validate OMM APEX-Ribo-Seq, we compared our data with OMM translation mapped by the LOV domain-controlled ligase for translation localization (LOCL-TL) ^99^. Importantly, OMM APEX-Ribo-Seq also detected most (85.3%) mRNAs identified by LOCL-TL (Figure S2R, “Detected in both”) and additionally captured many unique transcripts (Figure S2R, “Only detected in APEX”).

A comparison of the translation survey in the matrix and on the OMM further ensured the specificity of APEX-Ribo-Seq. OMM APEX-Ribo-Seq did not capture 13 CDSs encoded in the mitochondrial genome; conversely, matrix APEX-Ribo-Seq did not capture mRNAs locally translated on the OMM (Figure S2S). These data confirmed the specific detection of protein synthesis inside and outside mitochondria via the APEX-Ribo-Seq technique.

Overall, these benchmark analyses demonstrate the versatility and robustness of APEX-Ribo-Seq for probing subcellular translation across cellular compartments.

### Local translation atlas across 14 distinct subcellular compartments

The performance of APEX-Ribo-Seq led us to expand our survey to a wider variety of subcellular loci. We fused the APEX2 tag to well-defined bait proteins ^100^ (Figure S1A) to conduct proximity labeling at 12 additional loci—the peroxisome, *trans*-Golgi, *cis*- Golgi, lysosome, endosome, early endosome, clathrin-coated vesicle, plasma membrane, cell junction, outer nuclear membrane, intermediate filament, and centrosome (Figures S1A and S3A). We carefully selected the bait protein species and their expression levels. In particular, we generated cells with genomic knock-in of the centrosome marker CEP192 (Figure S1A), since the additional copies of other markers in stable integrants caused abnormal centrosome duplication or aggregation (Figure S3B). Again, we detected a typical footprint length with a 3-nt phase in APEX-Ribo-Seq, targeting specific loci with high reproducibility (Figure S3C-D). Overall, considering these sites along with the NES, matrix, ERM, and OMM, APEX-Ribo-Seq showed high reproducibility (Figure S3E). As for the ERM and OMM, we defined the locally translated mRNAs for each organelle; the number of such mRNAs ranged from 174 to 1159 at different sites (Figure S3F). Among the 14 loci (12 loci above, ERM, and OMM), mRNAs from 3,227 genes were locally translated at least one site (Figure S3G). We also assessed the empirical false-positive rate (see the figure legends for details) across the compartments and found that our threshold reasonably suppressed false-positive hits (the maximum is 4.2%) (Figure S3H). Although the efficiency of APEX2-mediated labeling may depend on the molecular size of the mRNA–ribosome complex—determined by transcript length and ribosome loading, with CDS length serving as a proxy, normalization to NES APEX-Ribo-Seq and input profiles (*i.e.*, organelle enrichment; Figure S2E) mitigated this potential bias (Figure S3I).

Our data closely corresponded with the known localizations of mRNAs identified via smFISH; for example, the outer nuclear membrane localization of *CLIP1*, *DDX11*, and *ADAM10* mRNAs and the *cis*-Golgi localization of *AKAP9* mRNA ^19^ were validated in our local translation profiles (Figure S3F). Furthermore, the mRNAs found in both lysosome and ERM APEX-Ribo-Seq may represent secretome translation at the ER-lysosome contact site (Figure S3J) ^101^.

To provide further confidence in our datasets, we performed smiFISH ^94^ for 12 mRNAs found via APEX-Ribo-Seq at the peroxisome, *trans*-Golgi, *cis*-Golgi, plasma membrane, and intermediate filament and found the colocalization of the mRNAs at the corresponding loci (Figure S3J-T), while nuclear paraspeckle *NEAT1* RNA ^102^, the negative control, did not show such a trend. In addition, local translation on the centrosome was validated by treatment with centrinone, a Polo-like kinase 4 (Plk4) inhibitor ^103^ that eliminates the centrosome, as Plk4 is required for the initiation of centriole assembly. The translation efficiency of mRNAs locally translated on the centrosome, as measured by standard Ribo-Seq and RNA-Seq, decreased upon centrinone treatment (Figure S3U).

To obtain an overview of the local translation on organelles, we conducted dimension reduction by t-distributed stochastic neighbor embedding (t-SNE) clustering ^104^ (Figure 2A). While the patterns of local translation at each subcellular compartment were separated overall, some of them were closer to each other than others (Figure 2A). These clusters included ERM and OMM, which have close physical contacts ^105^ (see below for details); the plasma membrane, endosome, and lysosome, which are connected to each other through membrane trafficking ^106,107^; early endosomes and clathrin-coated vesicles, which likewise are connected via trafficking ^108^; the *trans*-Golgi network (TGN) and clathrin-coated vesicles that bud from the TGN ^109^; and the *cis*-Golgi and centrosome, which are in close spatial proximity in cells ^110^ (Figure 2A). Not only the overall pattern similarity among each data set, but also overlaps of individual genes for each locus, were observed (Figure 2B). Despite the similarities among some organelles, 64% of the listed transcripts were locally translated at only one or two loci (Figure 2C), suggesting the specificity of local translation. We found that 1105 mRNAs (34%) were uniquely translated at a single locus (Figure 2D).

**Figure 2.**
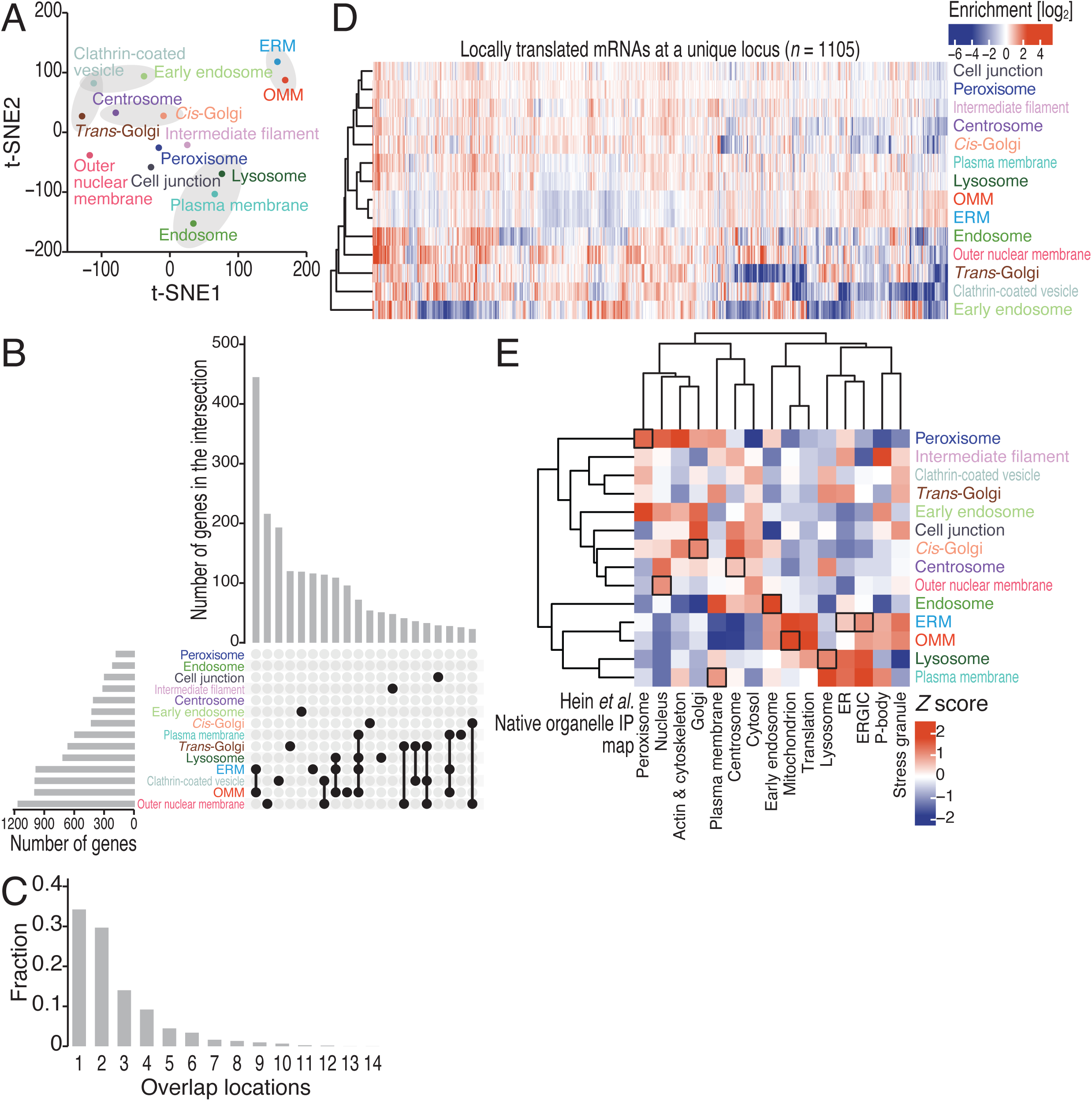
Local translation atlas for 14 subcellular compartments. (A) t-SNE plot ^193^ for mRNAs locally translated at least one site in 14 organelles (Figure S3G), according to organelle enrichment determined by APEX-Ribo-Seq (Figure S2E). The gray groups are described in the main text. (B) UpSet plot showing intersections of mRNAs among organelles analyzed by APEX-Ribo-Seq. (C) Fraction of overlap among locations where the mRNA is classified as undergoing local translation by APEX-Ribo-Seq. (D) Heatmap of organelle enrichment determined by APEX-Ribo-Seq (Figure S2E) for mRNAs exclusively found at a unique locus. The color scale indicates the degree of organelle enrichment. (E) Heatmap of the correspondence of locally translated mRNAs and the localization of proteins encoded by them. The gene sets uniquely translated at each locus (Figure 2D) were used. Protein localization is defined by native organelle immunopurification ^114^. The *Z* score represents the standardized enrichment or depletion of the localization term within the local translation category. The color scale indicates the *Z* score. The black box tiles are described in the main text. See also Figures S3-5 and Tables S1, S4, and S5.

### Dual local translation associated with organelle contacts

Given the highly similar translation profiles detected by the ERM and OMM APEX-Ribo-Seq (Figure 2A-B), we tested whether this stems from the inherent nature of the tight mitochondria-ER contact sites. We conducted mitochondria immunoprecipitation (MitoIP) using an anti-TOMM22 antibody, followed by Ribo-Seq in HEK293 T-REx cells (MitoIP-Thor-Ribo-Seq, here denoted simply as MitoIP-Ribo-Seq) ^89^ (Figure S4A and Table S1). Originally, this method was developed for the enrichment of mitoribosome footprints; however, here we focused on footprints originating from cytoribosomes attaching to purified mitochondria. In addition to mRNAs enriched in OMM APEX-Ribo-Seq, those in ERM APEX-Ribo-Seq showed a high enrichment score in the MitoIP fraction over the input standard Ribo-Seq (Figure S4B). The same trend could be found for the mRNAs defined by the BirA–AviTag system for ERM (Figure S4C). Moreover, we performed smiFISH for the 14 mRNAs whose translation was detected at both ERM and OMM by APEX-Ribo-Seq (Figure S4D-E). Our quantification revealed that those mRNAs have similar proximity to both the ERM and OMM, whereas nuclear *NEAT1* RNA did not (Figure S4E). Thus, ERM-translated mRNAs are closely located at the OMM.

To examine the relationship between the similarity of ERM-OMM APEX-Ribo-Seq and mitochondria-ER contact site formation, we also performed additional ERM and OMM APEX-Ribo-Seq in another cell line, U2OS (Figure S4F-H and Table S1); mitochondria in U2OS cells showed a more elongated morphology ^111^, which is associated with decreased levels of ER-mitochondria contact sites ^112^. We overviewed the similarity of local translation at the ERM and OMM in U2OS and HEK293 T-REx cells through t-SNE and observed a significant separation of ERM-OMM translation profiles in U2OS cells, consistent with the reduced ER-mitochondria contact sites in this cell line (Figure S4I). We further detected a significant distinction in the corresponding protein synthesis at the two sites (Figure S4J); mitochondria-localized proteins were more enriched in OMM APEX-Ribo-Seq, and ER-localized proteins were in ERM APEX-Ribo-Seq.

Notably, even in APEX-Ribo-Seq in HEK293 T-REx cells, which may have tight mitochondria-ER sites, we observed higher enrichment of cognate categories of genes; Gene Set Enrichment Analysis (GSEA) ^113^ revealed that ER-related categories showed higher scores in organelle enrichment in ERM APEX-Ribo-Seq than in OMM APEX-Ribo-Seq and vice versa (Figure S4K). The threshold-based qualitative definition of locally translated mRNAs at each locus may not be exclusive (Figures 1E-F and S2E and Table S1); our organelle enrichment values of APEX-Ribo-Seq quantitatively distinguished OMM and ERM.

Taken together, the high similarity of ERM and OMM local translation may represent the physiological nature of the mitochondria-ER contact sites. Thus, especially in HEK293 T-REx cells, these local translation profiles on the ERM and OMM should be considered as a pool of ER-mitochondria contacts and those organelle peripheries. To avoid technical issues, the downstream analysis, which required gene set definition, mainly focused on the unique genes found in the specific organelle (Figure 2D, *cf.* Figure S3G).

### Local translation for the local proteome

Our local translation atlas showed a close connection between the site of translation and the localization of the synthesized proteins. We employed the protein subcellular localization datasets generated by native organelle immunopurification ^114^ (Figure 2E), BioID-based proximity labeling ^100^ (Figure S5A), and immunostaining (The Human Protein Atlas) ^115^ (Figure S5B). The correspondence between local translation and protein localization was found in the ERM (Figures 2E, S5A, and S5B), OMM (Figures 2E, S5A, and S5B), endosomes (Figures 2E and S5A), lysosomes (Figures 2E and S5A), plasma membrane (Figures 2E, S5A, and S5B), cell junctions (Figures S5A and S5B), *trans*-Golgi (Figure S5A), *cis*-Golgi (Figures 2E and S5B), peroxisomes (Figure 2E and S5B), intermediate filaments (Figure S5B), and centrosomes (Figure 2E and S5B). Our data also showed the association between nuclear-localized proteins and their synthesis at the outer nuclear membrane (Figures 2E, S5A, and S5B). We note that due to the wider sub-cellular regions covered by protein localization studies, scores can be obtained for loci whose local translation was not directly assessed by our APEX-Ribo-Seq.

Detailed inspection of locally translated mRNAs revealed that key factors for organelle function were supplied locally. Functional enrichment analysis based on complex formation via STRING revealed that this trend was global (Figure S5C). Some membrane-embedded assemblies included the super complex of the Sec61 complex, the translocon-associated protein (TRAP) complex, and the oligosaccharyltransferase (OST) complex ^116,117^ for the ERM and the translocase of the outer membrane (TOM) complex ^118,119^ for the OMM (Figure S5D), reflecting the cotranslational membrane insertion of nascent peptides in these organelles.

In addition, APEX-Ribo-Seq revealed the on-site assembly of complexes peripherally associated with organelles. While the ERM and OMM are linked to the translation of mRNAs encoding membrane-inserted or translocated proteins, few examples of this phenomenon have previously been reported for other organelles. Indeed, the detection of mRNAs encoding membrane proteins was limited to peroxisomes, endosomes, and cell junctions, and to the centrosome, which is membraneless (Figure 3A). These findings suggested that remaining proteins locally synthesized consist of auxiliary organelle-associated factors. Indeed, we found the following examples: the outer nuclear membrane, nuclear pore outer ring Nup107-160 subcomplex ^120,121^; the clathrin-coated vesicle, COPI coat complex ^122,123^, and HOPS tethering complex ^124,125^; the *trans*-Golgi, TRAPPIII complex ^126,127^ (Figure 3B).

**Figure 3.**
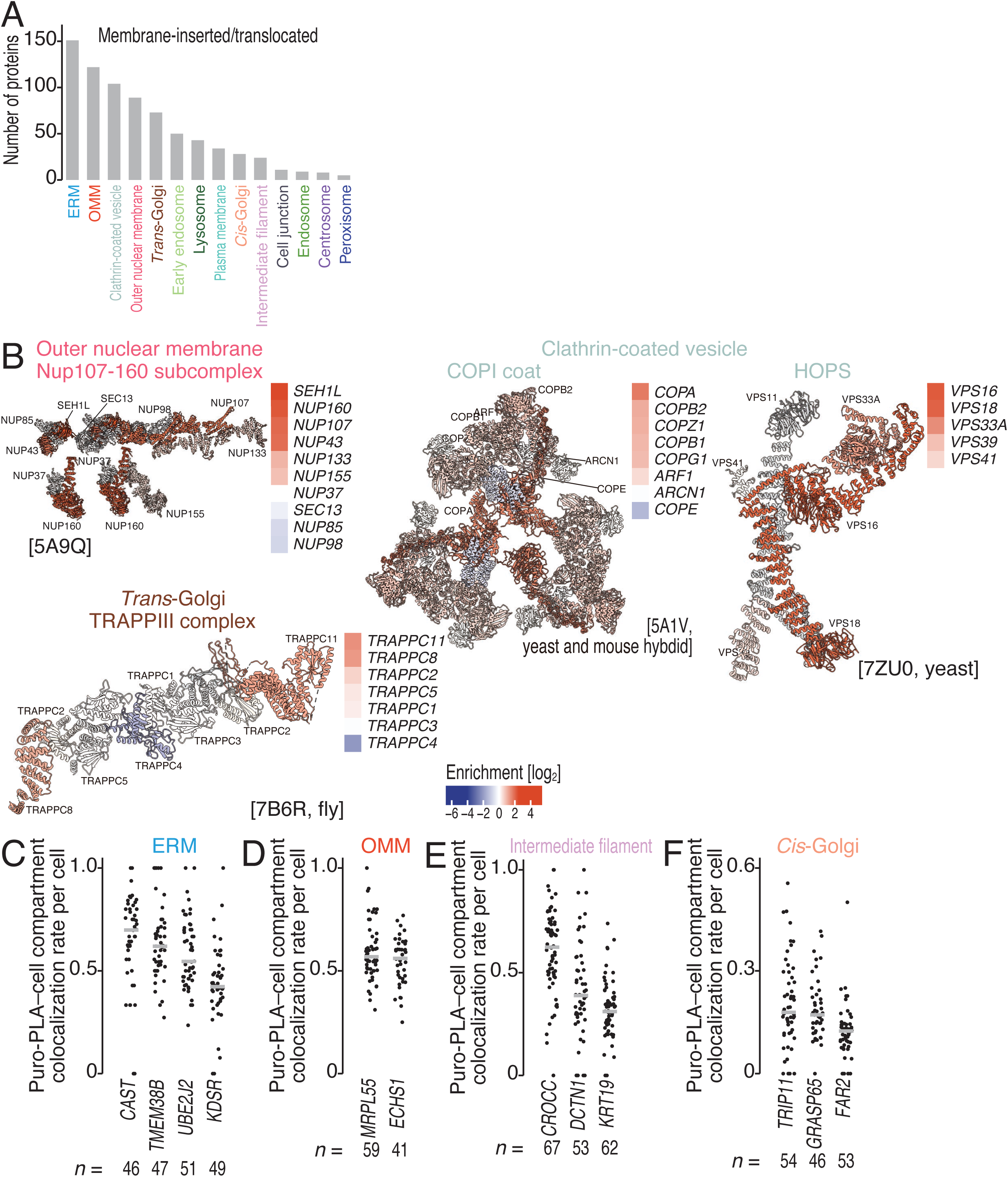
Local translation supplies subunits of complexes peripherally associated with organelles. (A) The number of membrane-inserted or membrane-translocated proteins (see Methods for details) for each local translation category. Gene sets uniquely translated at each locus (Figure 2D) were used. (B) Structural mapping of locally translated proteins at the indicated locus. The structures of the Nup107-160 subcomplex (Protein Data Bank [PDB]: 5A9Q) ^121^, COPI coat (PDB: 5A1V) ^123^, HOPS (PDB: 7ZU0) ^124^, and TRAPPIII complex (PDB: 7B6R)^127^ are shown by UCSF Chimera X (ver. 1.1) ^194^. The color scale in the structure and heatmap of organelle enrichment determined by APEX-Ribo-Seq (Figure S2E) indicates the degree of organelle enrichment. (C-F) Quantification of the colocalization rates of the marker proteins and the Puro-PLA signal for the indicated proteins/RNAs is shown. The median (bars) and individual data (points, n for the number of cells analyzed) are shown. See also Figure S5 and Tables S1 and S7.

Given that protein complexes can be formed through the interaction of two nascent chains emerging from two ribosomes (so-called co-co–assembly) ^128,129^, we reasoned that on-organelle translation may serve as a unified platform that simultaneously facilitates both cotranslational complex assembly and localization. We examined the candidate co-co–assembly pairs in a structure-based cotranslational assembly prediction dataset ^129^ and cross-referenced them with our local translation atlas (Figure S5E). This analysis identified 204 protein pairs that co-co–assembled locally, among which 103 pairs were cotranslated at only a single locus (Figure S5E). Consistent with our observation of local supply on clathrin-coated vesicles (Figure 3B), the HOPS complex subunits VSP16 and VPS33A were co-co–assembled at this site (Figure S5E). At the OMM, we also found potential co-co–assembly of the alpha and beta subunits of several nascent polypeptide-associated complex (NAC) chaperones (NACA–BTF3, NACA–BTF3L4, NACA2–BTF3, and NACA2–BTF3L4), which are required for the targeting of polytopic proteins to the OMM ^130^ (Figure S5E).

Because APEX-Ribo-Seq relies on relative enrichment analysis (Figure S2E), this method did not directly quantify the absolute fraction of mRNAs undergoing local translation. To assess the actual fraction of translation that occurred locally, we applied puromycylation with the proximity ligation assay (Puro-PLA) ^131,132^. We note that these experiments are restricted to a subset of targets whose antibodies are commercially available and compatible for immunostaining. Despite such a limitation, our Puro-PLA experiments monitored local translation of 12 mRNAs for ERM, OMM, the intermediate filament, and the *cis*-Golgi (Figures 3C-F and S5F-I). The fraction of mRNAs locally translated varied across organelles; a high local translation fraction (∼70%) was observed on ERM, while a moderate rate (∼18%) was found at the *cis*-Golgi.

### Localization of mRNAs and on-site translational regulation

Then, we investigated the general rules that allow efficient local translation. The mechanisms could include the following: 1) nascent chain-mediated recruitment (such as signal peptide recognition by the SRP for on-ERM translation), 2) mRNA recruitment, 3) the on-site control of translation efficiency, or 4) a combination of these mechanisms (Figure S6A). Mechanisms 1) and 2) must align with mRNA localization on the organelles. To quantify the role of RNA localization in local translation, we conducted APEX-RNA-Seq in parallel using the same samples used for APEX-Ribo-Seq (Figures 1A and S6B) and calculated the enrichment using the same method that was applied for APEX-Ribo-Seq (Figure S2E). We observed a diverse range of correlations between the APEX-Ribo-Seq and APEX-RNA-Seq results across subcellular compartments (Figure 4A-B). We detected high correspondence at the centrosome, ERM, and OMM. The centrosome is known to recruit mRNA ribosome complexes through emerging nascent chains ^19,22,25^, whereas the ERM and OMM feature the cotranslational membrane insertion of nascent chains ^58,61,98,116,133,134^. In contrast, translation efficiency control was observed to make a greater contribution at intermediate filaments, endosomes, and the plasma membrane (Figure 4A-B). Statistically, no clear difference between locus-matched correlation and cross-correlation was observed in those loci (Figure 4A, color bar vs. gray bar). This also highlighted that the connection between mRNA localization and local translation is weaker in these compartments, supporting the idea that translational regulation plays a particularly prominent role there.

**Figure 4.**
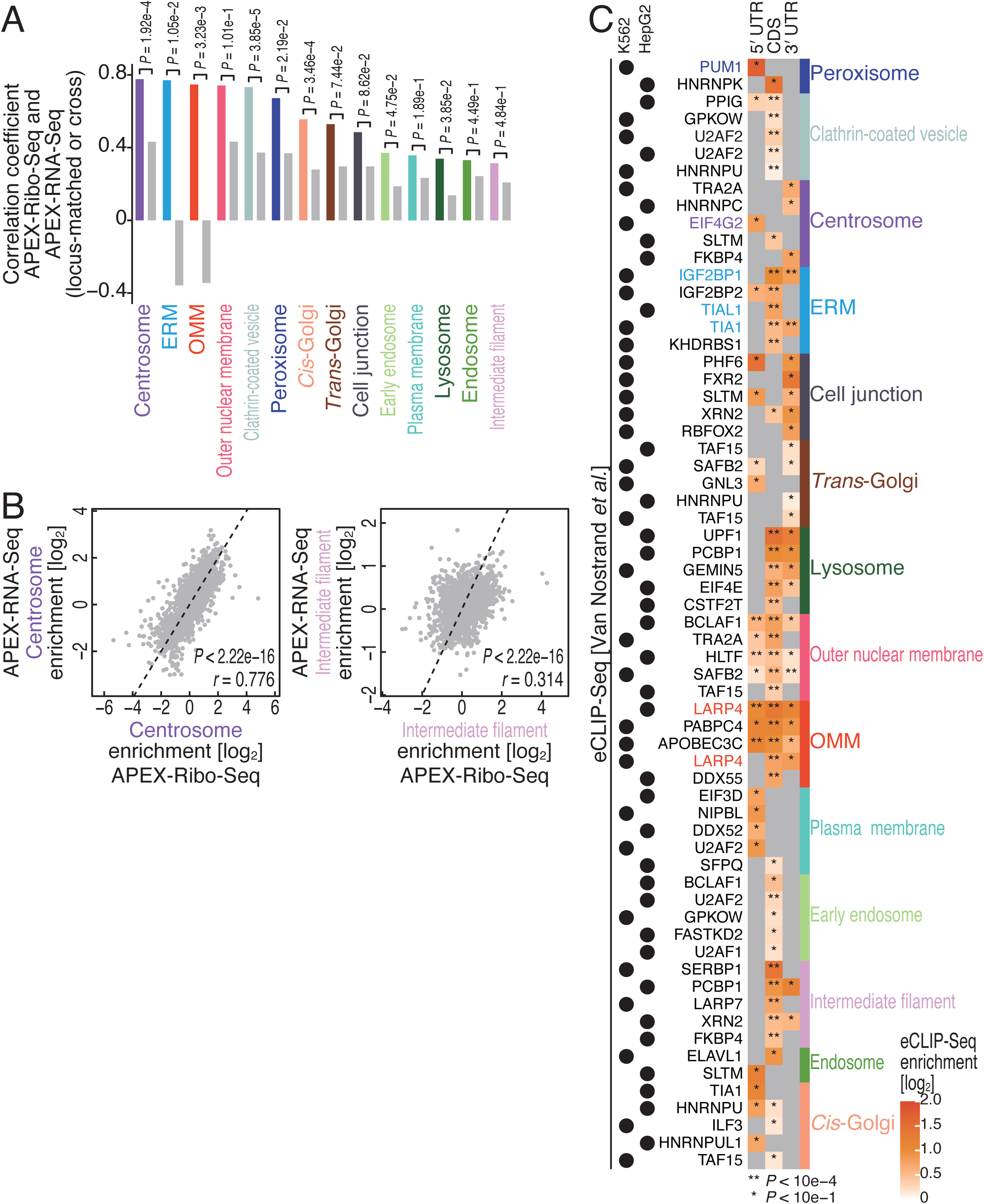
Local translational control beyond RNA localization. (A) Bar plots of the Pearson’s correlation coefficients between organelle enrichment (locus matched) determined by APEX-Ribo-Seq and that by APEX-RNA-Seq (Figure S2E); those *P* values (Pearson’s product-moment correlation test [two-sided]) are less than 2.22e−16 in all cases. Gray bars represent the cross-correlation of the organelle enrichment determined by APEX-Ribo-Seq vs. that determined by APEX-RNA-Seq in other loci. *P* values comparing the locus-matched correlation against the corresponding cross-correlation distribution were computed by a cross pair label permutation test (one-sided) and adjusted by the Benjamini–Hochberg method. (B) Scatter plots of organelle enrichment determined by APEX-Ribo-Seq and APEX-RNA-Seq (Figure S2E) at the indicated loci. *P* values were calculated with Pearson’s product-moment correlation test (two-sided). (C) Heatmap of the enrichment of eCLIP-Seq reads within the 5′ UTR, CDS, and 3′ UTR for the indicated group of mRNAs. The top five RBPs (log_2_ eCLIP-Seq enrichment > 0 and *P* value < 0.1), ranked by *P* value calculated with the Mann Whitney *U* test (two-tailed) and adjusted by the Benjamini–Hochberg method, were visualized. All the results are summarized in Table S6. The color scale indicates the degree of the eCLIP-Seq read enrichment. Gene sets uniquely translated at each locus (Figure 2D) were used. *r*, Pearson’s correlation coefficient. See also Figure S6 and Tables S4 and S6.

We note that ribosome occupancy (APEX-Ribo-Seq enrichment over APEX-RNA-Seq enrichment, Figure S6C) is low in the mitochondrial matrix (or mitoribosome-mediated translation), which is consistent with the previously observed extremely slow translation kinetics ^89,135^.

### RNA-binding proteins and local translation

Given the substantial contribution of on-site translation control, we reasoned that RNA-binding proteins (RBPs) might affect subcellular mRNA translation (Figure S6A). Indeed, recent studies highlighted the importance of RBPs in local translation even for the loci where nascent chains have been thought to play a key role ^136^. To explore RBP binding profiles across the transcriptome, we harnessed the collection of enhanced crosslinking and immunoprecipitation followed by sequencing (eCLIP-Seq) datasets covering 250 conditions for 168 RBPs in two cell lines (K562 and HepG2) ^137^. eCLIP-Seq reads on 5′ untranslated regions (UTRs), CDSs, and 3′ UTRs were evaluated separately. We found that some RBPs exhibited high loading (*i.e.*, more eCLIP-Seq reads compared to the size-matched input samples) on unique mRNAs locally translated on specific organelles (Figure 4C). Our findings aligned with those of prior studies, such as TIA1/TIA1L binding to the 3′ UTRs of mRNAs translated on the ERM ^138^ and LARP4 localization to the OMM (see below for further analysis) ^139–141^. Additionally, PUM1 binding to mRNAs locally translated on the peroxisome aligned well with reports that Puf5, a PUM1 homolog, recruits mRNAs that encode peroxisome proteins to the peroxisome in yeasts ^142^.

### Prediction of local translation via language models

A straightforward mechanism of localized translation is that mRNAs carry short RNA motifs that can be recognized by RBPs already present at specific sites. We analyzed mRNA sequences for such motifs using HOMER (see Methods for details; Figure 5A and Table S2) and found many motifs that were indeed enriched in the sequences with high organelle enrichment. However, the short motifs occurring in the mRNAs were diverse across the subcellular compartments; for example, many motif hits were found for mRNAs translated on the intermediate filaments, outer nuclear membrane, centrosome, *cis*-Golgi, clathrin-coated vesicle, and *trans*-Golgi, whereas the mRNAs translated at the other organelles possessed few or no enriched motifs (Figure 5A).

**Figure 5.**
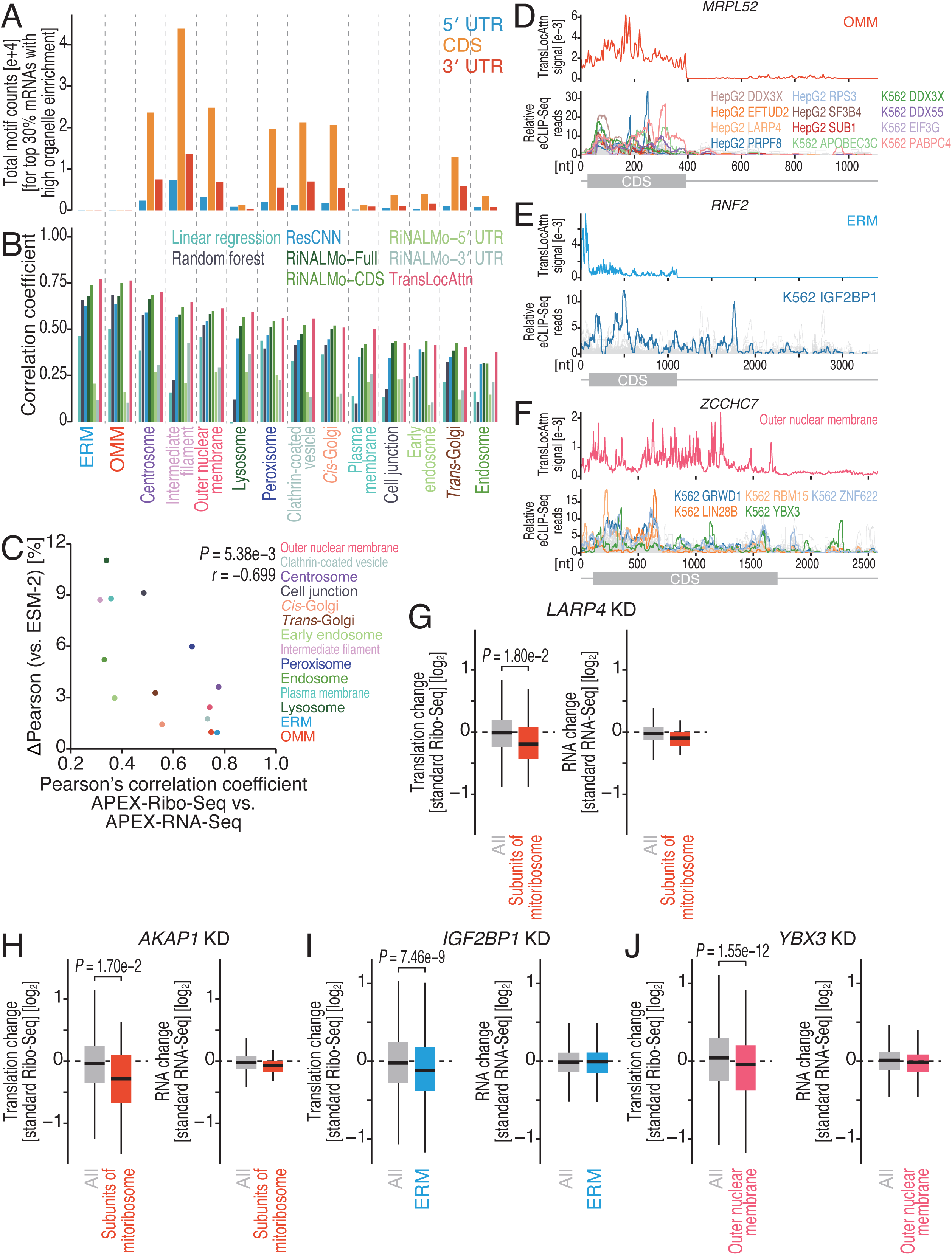
Prediction of local translation via a language model. (A) Occurrence of short motifs (5-7 mer) that are enriched in the mRNAs with high organelle enrichment according to APEX-Ribo-Seq. (B) Pearson’s correlation between machine-learning-predicted and actual organelle enrichment determined by APEX-Ribo-Seq on the test set. The linear regression and random forest models included only the lengths of the 5′ UTR, CDS, and 5′ UTR as inputs. ResCNN, ResNet using one-hot encoded sequences; RiNALMo-Full/CDS/5′ UTR/3′ UTR, feed-forward network (FNN) applied to mean-pooled RiNALMo embeddings from the specified sequence region (full mRNA sequence, CDS, 5′ UTR, or 3′ UTR); TransLocAttn, FNN applied to attention-weighted mean-pooled RiNALMo embeddings. See Methods for model details. (C) Scatter plot of Pearson’s correlation coefficients between organelle enrichment determined by APEX-Ribo-Seq and APEX-RNA-Seq versus the gain in prediction by RNA sequence over the ESM-2 amino-acid sequence baseline across the 14 subcellular compartments. ΔPearson indicates the increase in held-out Pearson’s correlation coefficient obtained by adding the RiNALMo full-transcript residual prediction to ESM-2. *P* value was calculated with Pearson’s product-moment correlation test (two-sided). (D-F) Examples of attention weights from TransLocAttn (top) and eCLIP-Seq read distributions (bottom) along the indicated mRNAs. eCLIP-Seq datasets with high correlation with the attention weights are shown in color; all other eCLIP-Seq datasets (*i.e.*, those that did not meet the threshold) are shown in gray. (G-J) Box plots of changes in Ribo-Seq and RNA-Seq after siRNA-mediated knockdown of *LARP4* (Figure 5G), *AKAP1* (Figure 5H), *IGF2BP1* (Figure 5I), and *YBX3* (Figure 5J) in the indicated mRNA groups. *P* values were calculated with the Mann Whitney *U* test (two-tailed). Effect sizes of RBP knockdown were estimated by the Hodges–Lehmann (HL) statistic (*LARP4*, HL = -0.145, 95% CI [-0.271, -0.016]; *AKAP1*, HL = -0.225, 95% CI [-0.410, -0.036]; *IGF2BP1*, HL = -0.116, 95% CI [-0.150, -0.082]; *YBX3*, HL = -0.125, 95% CI [-0.154, -0.096]). Box plots show the median (centerline), upper/lower quartiles (box limits), and 1.5× interquartile range (whiskers). See also Figure S6 and Tables S2 and S6.

To assess whether RNA sequence features beyond short motifs determine organelle-specific protein synthesis, we developed models to predict local translation levels from sequence information. First, we tested the predictive power of RNA length alone, since we observed differential lengths in CDS across locally translated mRNAs (Figure S6D). Although we observed differential CDS size (especially short CDSs for mRNAs enriched in OMM and ERM APEX-Ribo-Seq) (Figure S6D), this feature could simply represent the characteristics of proteins functioning in each organelle, rather than the direct determinant of local translation specificity. Nonetheless, this feature was strongly reflected in modeling; simple linear regression and random forest models using only sequence length effectively predicted the log_2_ organelle enrichment in APEX-Ribo-Seq (Figures 5B and S6E).

For better prediction, we next employed machine learning approaches using the full nucleotide sequence to explore more complex sequence determinants. A key advantage of nucleotide language models ^143^ is their ability to embed sequences of different lengths into a fixed-size numerical vector that can capture the most salient information on the basis of the context learned during large scale pre-training. We first predicted translation location from embeddings generated by the RNA language models AIDO.RNA ^144^ and RiNALMo ^145^, as well as the DNA model Evo 2 ^146^. In all three cases (Figure S6F), even simple mean pooling of embeddings derived from the full sequences, coupled with a standard feedforward network, yielded superior predictive performance to a baseline convolutional neural network (Figure 5B). Organelle enrichments not associated with short motifs (at the ERM and OMM), RNA length (at intermediate filaments), or both (at lysosomes, cell junctions, the plasma membrane, endosomes, and early endosomes) were predicted reasonably well (Figure 5B), indicating that the language-model embeddings capture information beyond simple motif count and sequence length. Notably, the signal was concentrated in the coding sequence: for all three models, mean-pooled embeddings derived solely from the CDS consistently outperformed embeddings of the full sequence (Figures 5B and S6F), so a partial sequence is not trivially subsumed by the whole-sequence embedding.

This raised a question about what the CDS embeddings represent, because the coding sequence determines the amino-acid sequence of the encoded protein, and protein identity is itself informative about where a protein functions. To separate these possibilities, we replaced the RNA input with the protein it encodes and predicted compartment enrichment from ESM-2 protein embeddings ^147^ (Figure S6F). Protein sequence alone was highly predictive (mean target-wise Pearson’s correlation, 0.527), below RiNALMo CDS embeddings (0.552, which exceeded ESM-2 in all five seeds) but above RiNALMo full-transcript embeddings (0.513; Figure S6F).

If protein sequence is this predictive on its own, does RNA sequence add anything? We predicted compartment enrichment from ESM-2 first and then tested whether frozen RiNALMo representations could predict what the protein baseline had missed, using the same held-out genes throughout. RNA representations did add information: adding a UTR-based prediction of the protein-model residual raised the mean held-out correlation slightly over ESM-2 alone and improved 12 of 14 compartments, whereas the same UTR features shuffled between genes gave no improvement (Figure S6G-H). CDS and full-transcript representations gave larger gains and improved all 14 compartments. Notably, the size of this gain was not uniform across cellular compartments, and its distribution tracked the mode of local translation.

Ranking compartments by gain in the full-transcript sequence over amino-acid sequence inversely associated with correspondence between APEX-Ribo-Seq and APEX-RNA-Seq (Figure 5C); compartments where nascent chain-mediated recruitment is expected, such as the ERM and OMM, showed the highest prediction by amino-acid sequences. Conversely, local translation in other compartments may also be influenced by features encoded in the RNA sequence.

To ask whether this information is positional, we modified the sequence itself, comparing a high-scoring window with a matched low-scoring window in the same gene under constraints that leave the composition or the protein intact: exact dinucleotide-preserving shuffles in UTRs, and synonymous recoding in CDSs, which leaves the encoded amino-acid sequence and the ESM-2 input unchanged. Window scores were chosen by how well they predicted the effect of masking each window on validation genes (Figure S6I; see Methods). High-scoring windows changed the prediction about 60% more than matched low-scoring windows in UTRs and about a third more in CDSs (median per-gene ratios 1.60 and 1.34; 161 and 168 held-out genes). The model therefore uses nucleotide-level information that is not reducible to the encoded amino-acid sequence.

To pinpoint position-dependent RNA sequence information, we developed TransLocAttn, a regression model that applies compartment-specific attention pooling to concatenated RiNALMo embeddings from the 5′ UTR, CDS, and 3′ UTR sequences (Figure S6J). The model achieved strong predictive accuracy for local translation across the 14 compartments on a held-out test set. A capacity-matched mean-pooling model was slightly less accurate (Figure 5B). TransLocAttn learned nonuniform, transcript- and compartment-dependent pooling weights (Figures 5D-F and S6L-N, top) which could be interpreted as prediction-associated candidate-region scores. A predictive advantage of attention pooling does not, however, make the weights faithful explanations of the model’s decisions ^148^. We therefore treated these profiles as candidate-region scores rather than causal effects, and tested RNA information and position-level attribution separately below.

The presence of attention profile signals in CDSs of the RNA (Figures 5D-F and S6L-N, top) could be explained either by nascent chain-mediated recruitment or by RBP binding. To test the latter possibility, we investigated whether the regions highlighted by the attention mechanism corresponded to known protein-RNA interactions. We systematically compared the attention profiles against RBP binding profiles experimentally determined by eCLIP-Seq ^137^. We aligned the smoothed attention tracks and eCLIP read distributions (see Methods for details). The compartment-specific attention track was weakly but positively aligned with eCLIP occupancy in both region sets, exceeding a null that circularly shifts the track within the same transcript regions (Figure S6K). Examples include IGF2BP1 binding to *RNF2* mRNA, detected in ERM APEX-Ribo-Seq (Figure 5E, bottom), and RBM15 binding to *JARID2* mRNA, locally translated at the centrosome (Figure S6L, bottom). Interestingly, multiple RBPs seem to target the same regions with high attention signal (Figures 5D, 5F, and S6L-N, bottom).

To test whether selected RBPs affect translation of compartment-associated gene sets, we focused on LARP4. Gene-set-level eCLIP enrichment nominated LARP4 in OMM-associated mRNAs (Figure 4C), whereas MRPL52 provided a descriptive local example in which the TransLocAttn profile overlapped the LARP4 eCLIP signal (Figure 5D). We also tested the involvement of AKAP1 in OMM translation, since this protein also presented eCLIP-Seq reads on mRNAs translated on the OMM (Figure S6O). Moreover, earlier studies have also suggested that LARP4 and AKAP1 may mediate OMM translation ^139,141^. Indeed, standard Ribo-Seq revealed reduced translation of mRNAs encoding mitoribosome subunits upon the knockdown of *LARP4* and *AKAP1* (Figure 5G-H).

As further validation, we tested IGF2BP1 for a role in ERM translation, as this RBP showed high loading across mRNAs locally translated on the ERM (Figure 4C) and a correspondence between the attention profile and the eCLIP-Seq read distribution on *RNF2* mRNA (Figure 5E). We detected decreased protein synthesis from ERM-translated mRNAs upon *IGF2BP1* knockdown (Figure 5I).

For the third locus for validation, we focused on mRNAs locally translated on the outer nuclear membrane, for which the eCLIP-seq pattern of YBX3 and FXR2 showed descriptive local correspondence with pooling-weight profiles on *ZCCHC7* (Figure 5F) and *CUL1*/*KIF1B* mRNAs (Figure S6M-N), respectively. Those RBP KDs again reduced the translation from mRNA subsets translated at the outer nuclear membrane (Figures 5J and S6P).

Together, the gene-set enrichments, selected local correspondences, and knockdown phenotypes indicate that RBP-guided mechanisms can explain the specificity of local translation.

### APEX-Ribo-Seq in primary neuron culture

We extended our investigation to mouse primary neurons and local translation at dendrites, axons, and presynapses ^11^ (Figure S7A). APEX2 tagging of Map2 ^149^, Nefl ^150^, and Snca ^151^ enabled labeling at dendrites, axons, and presynapses (Figure S7B). We observed that the transduction of AAV used here led to limited effects on local translation, at least on ERM, monitored by *in situ* detection of puromycylated peptides ^152^ (Figure S7C-D). Genome-wide Ribo-Seq also detected a restricted number of genes with significant changes by AAV infection [35 / 14,023 (0.250%) for Map2, 3 / 14,023 (0.021%) for Nefl, and 1 / 14,023 (0.007%) for Snca], beyond the expressed marker proteins (Figure S7E). As found in HEK293 T-REx (Figure S1G-H), short H_2_O_2_ treatment did not accumulate phosphorylated eIF2α, especially on OMM (Figure S7F-G), where local phosphorylation of eIF2α has been reported ^153^. Again, both puromycylation (Figure S7H-I) and Ribo-Seq (14 among 14597 detected genes, 0.096%) (Figure S7J) showed minimal alteration of translation by H_2_O_2_ treatment in our conditions.

Using this setup, we conducted APEX-Ribo-Seq in the corresponding neuronal compartment (Figure S7K-M and Table S3) and calculated the read enrichment relative to the input (Figure S7N). We identified 1023 genes locally that were translated in neuronal subcellular compartments (Figure 6A-B). Gene Ontology (GO) analysis confirmed the enrichment of mRNAs encoding proteins associated with corresponding CCs (Figure S7O). Comparison of our APEX-Ribo-Seq dataset with previously reported datasets, including RNA-Seq ^154^, Ribo-Seq ^53^, and translating ribosome affinity purification (TRAP)-Seq ^155^ from microdissected neuropil data, revealed moderate concordance (Figure S7P), which is at least partially due to the mixture of diverse neuronal compartments in neuropils. Recent data from TurboID-based proximity ligation (PL)-Ribo-Seq of dendrites ^63^ were most congruent with the APEX-Ribo-Seq profiles, especially for dendrite translation surveys (Figure S7P-Q). The majority of surveyed mRNAs were translated at only one of the studied neuronal compartments (Figure 6C-D), indicating that these compartments possess distinct translation programs.

**Figure 6.**
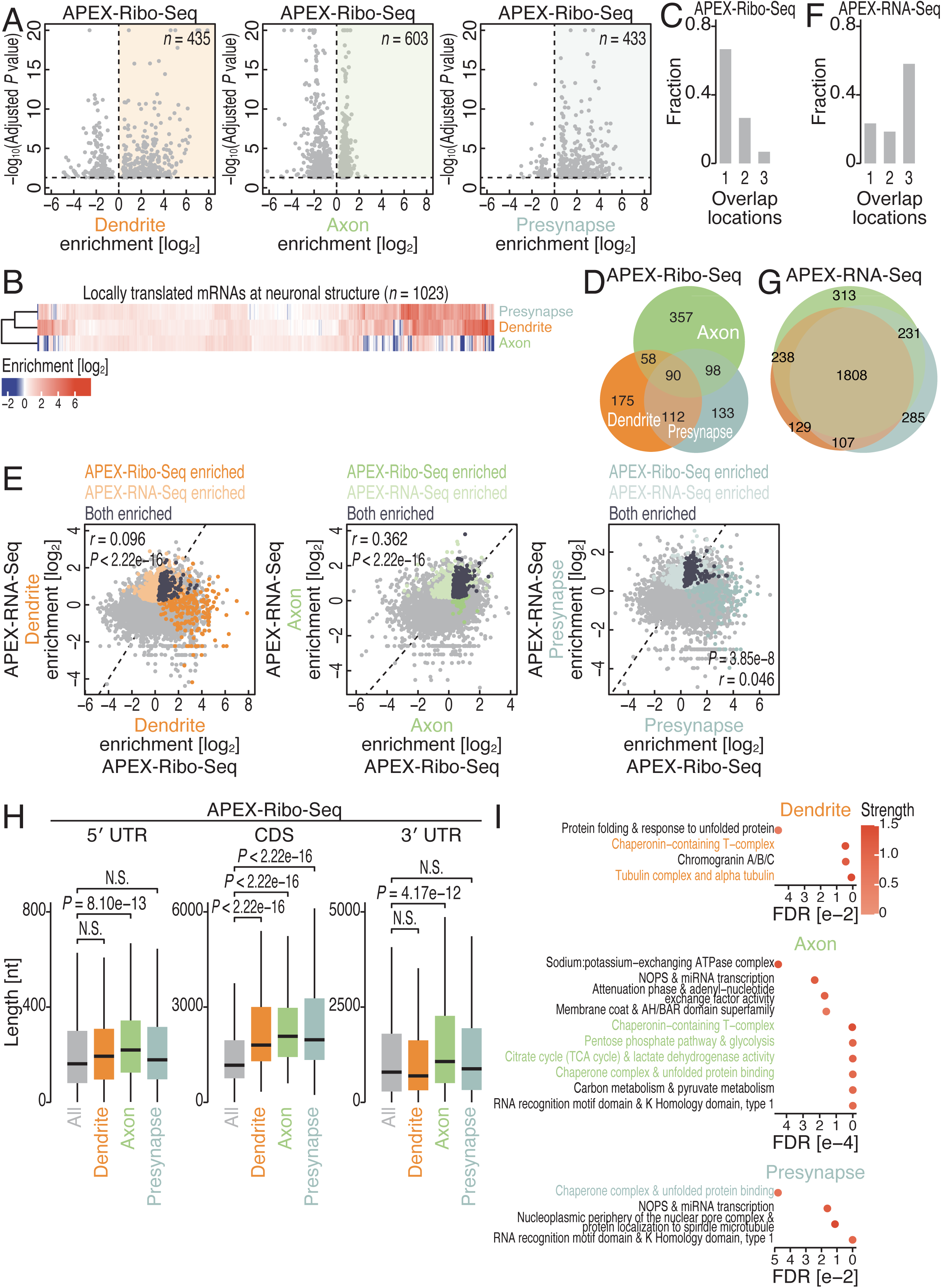
Local translation profiling of neuronal compartments. (A) Volcano plots showing neuronal compartment enrichment (Figure S7N) for dendrites, axons, and presynapses determined by APEX-Ribo-Seq. mRNAs with an adjusted *P* value of 0.05 or lower are shown. The colored areas represent enriched mRNAs. (B) Heatmap of neuronal compartment enrichment determined by APEX-Ribo-Seq (Figure S7N) for the indicated neuronal compartments. The color scale indicates the degree of neuronal compartment enrichment. (C) Fraction of overlap among locations where the mRNA is classified as undergoing local translation by APEX-Ribo-Seq. (D) Venn diagram of locally translated mRNAs in the indicated neuronal compartments. (E) Scatter plots of neuronal compartment enrichment as determined by APEX-Ribo-Seq and APEX-RNA-Seq (Figure S7N). Localized or locally translated mRNAs defined by APEX-RNA-Seq and APEX-Ribo-Seq are highlighted. *P* values were calculated with Pearson’s product-moment correlation test (two-sided). (F) Fraction of overlap among locations where the mRNA is classified as undergoing local translation by APEX-RNA-Seq. (G) A Venn diagram of localized mRNAs in the indicated neuronal compartments (H) Box plots of the lengths of the 5′ UTR, CDS, and 3′ UTR of the indicated mRNA groups defined by APEX-Ribo-Seq. *P* values were calculated with the Mann Whitney *U* test (two-tailed) and adjusted by the Benjamini–Hochberg method. (I) Complexes encoded in locally translated mRNAs in the indicated neuronal compartments were assessed by STRING analysis ^93^. The results of the top 10 terms, ranked by false discovery rate (FDR) value, after removing clusters present in more than three compartments, are shown. Terms shown in color are described in the main text. The color scale indicates the strength of the associations in the network (ranging from 0 to 1), which is a confidence indicator ^93^. *r*, Pearson’s correlation coefficient. Box plots show the median (centerline), upper/lower quartiles (box limits), and 1.5× interquartile range (whiskers). See also Figure S7 and Tables S3 and S4.

The assessment of RNA localization with simultaneous APEX-RNA-Seq at each locus (Figure S7R) revealed a low correlation with local translation (Figure 6E). As in APEX-Ribo-Seq (Figure S7N), we identified more than 2000 mRNAs localized to each compartment (Figure S7S). In contrast to the local translation profiles (Figure 6C-D), the RNA localization profiles could not be used to distinguish the neuronal compartments (Figure 6F-G); a wider array of mRNAs was localized to two or three neuronal compartments. Indeed, our findings were consistent with those of previous studies, suggesting highly mRNA-selective translational activation ^156^ and simultaneous maintenance of other mRNAs at low translation efficiency before stimuli ^45,54,157^. This may be related to the formation of neuronal RNP granules ^158^, which could maintain mRNAs inaccessible to translational machinery.

The 5′ UTRs, CDSs, and 3′ UTRs of neuropil-localized/translated mRNAs are longer than those of somatic mRNAs ^63,155^. The characteristics of the mRNAs present at the three neuronal compartments were consistent with these features (Figure S7T). However, locally translated mRNAs showed a differential contribution of those RNA elements, suggesting that 5′ UTR and 3′ UTR length may not be detrimental for on-site translation at dendrites and presynapses (Figure 6H).

We also explored the enrichment of complex subunits supplied in neuronal compartments by STRING. The local synthesis of chaperonin-containing t-complexes (CCT complexes, also known as T-complex protein Ring Complexes, or TRiCs) and/or chaperone complexes (including Hsp90, Hsp70, and Hsp40) in the neuronal compartments (Figure 6I) suggested the importance of local protein folding. We also identified the tubulin complex, which functions as a key determinant of neuronal polarity beyond a mere structural element ^159^, as a complex supplied specifically in dendrites, and complexes for glycolysis and the citrate cycle, which may facilitate the maintenance of local ATP ^160^, supplied specifically in axons (Figure 6I).

### In vivo APEX-Ribo-Seq in fly embryos

Finally, we applied our technique in an *in vivo* context, focusing on *Drosophila* embryogenesis, especially at the germ granule, an RNA-protein condensate (Figure 7A) ^41,161^. The physiological significance of local translation on this nonmembranous organelle is that it enables the spatial and temporal production of germline determinants required for the specification and development of primordial germ cells (PGCs) to be established, while preventing ectopic protein production in the soma (Figure 7A). Recent studies ^10,41^ used nascent chain tracking (NCT) ^50,162^ and showed local translation of *nanos* (*nos*) mRNA on the surface of the germ granule, which provides localized Nanos protein, helping maintain germ cell identity, repressing somatic differentiation programs, and regulating the PGC transcriptome. However, beyond *nos* mRNA, the comprehensive view of the locally translated mRNAs on the germ granules and their dynamics before and after PGC establishment remains unexplored.

**Figure 7.**
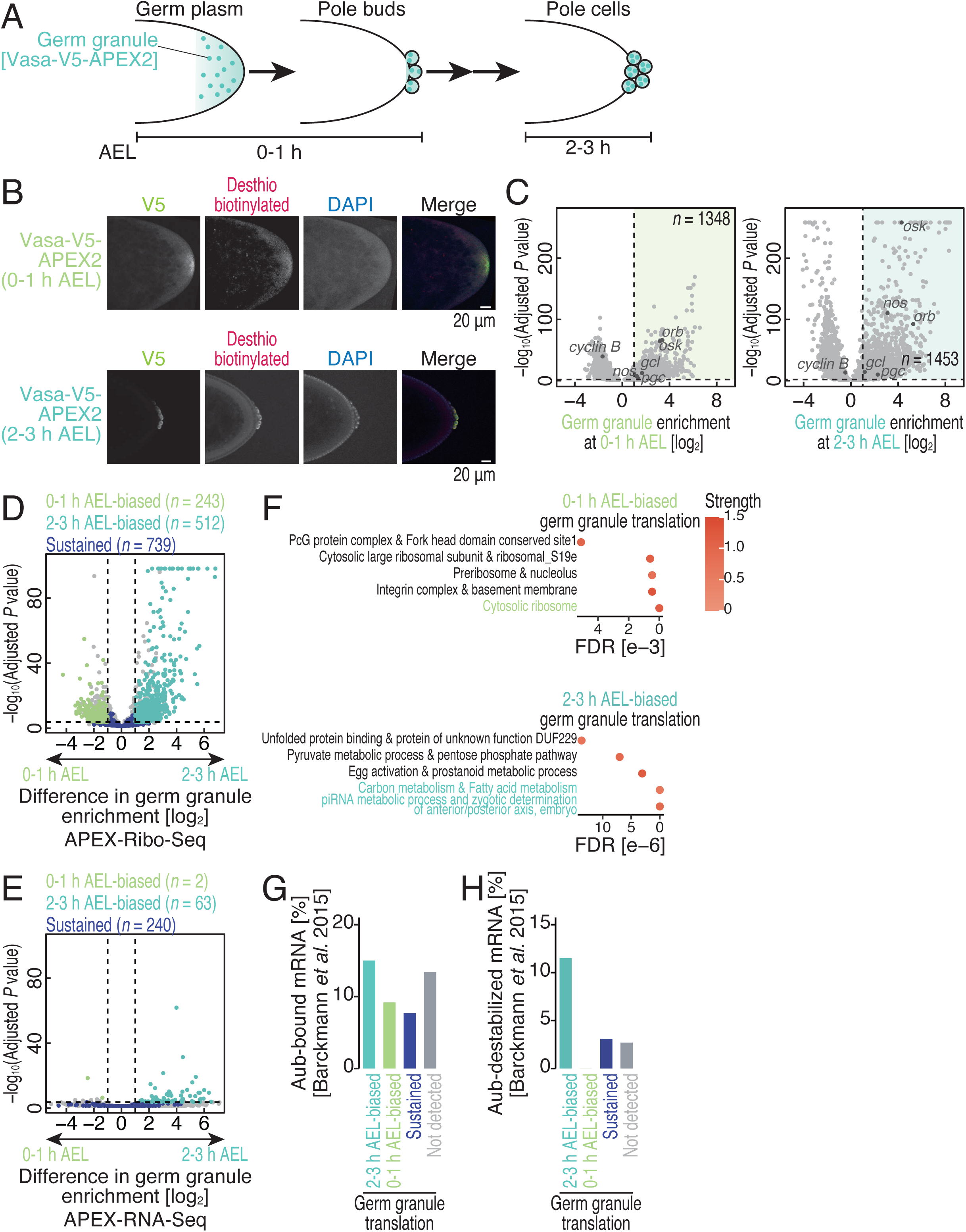
Local translation dynamics on fly germ granules. (A) Schematic of APEX-Ribo-Seq and APEX-RNA-Seq on germ granules along fly embryogenesis. (B) Immunofluorescence analysis of the indicated proteins (detected by V5) and desthiobiotinylated proteins/RNAs (detected with Alexa Fluor 555-conjugated streptavidin) in fly embryos. DAPI (4’,6-diamidino-2-phenylindole) was used to stain the DNA. The scale bar represents 20 μm. (C) Volcano plots showing germ granule enrichment determined by APEX-Ribo-Seq (Figure S8F). mRNAs with an adjusted *P* value of 0.05 or lower are shown. The colored areas represent enriched mRNAs. The genes mentioned in the main text are highlighted. (D) Volcano plots showing differences in germ granule enrichment in the embryogenesis stage (Figure S8G) by APEX-Ribo-Seq. Genes with 0-1 h AEL-biased, 2-3 h AEL-biased, and sustained translation are highlighted. (E) Volcano plots showing differences in germ granule enrichment in the embryogenesis stage (Figure S8G) by APEX-RNA-Seq. Genes with 0-1 h AEL-biased, 2-3 h AEL-biased, and sustained RNA localization are highlighted. (F) Complexes encoded in locally translated mRNAs in a stage-biased manner (Figure 7D) were assessed by STRING analysis ^93^. The significant terms, ranked by false discovery rate (FDR) value, are visualized. The terms shown in color are described in the main text. The color scale indicates the strength of the associations in the network (ranging from 0 to 1), which is a confidence indicator ^93^. (G and H) The fraction of Aub-bound (Figure 7G) and Aub-destabilized mRNAs (Figure 7H) for each category of stage-biased translation on germ granules (Figure 7D). See also Figure S8 and Tables S3, S4, and S7.

To perform APEX-Ribo-Seq at fly germ granules, we knocked APEX2 into the endogenous locus of *vasa*, which encodes the germ granule core protein Vasa, using the CRISPR-Cas9 system (Figure S8A) ^163^. To facilitate the efficient entry of desthiobiotin-phenol and H_2_O_2_ into small-molecule–refractory embryos, we optimized the protocol using digitonin-mediated permeabilization originally developed for the fly ovary ^164,165^. Using this setup, we conducted proximity labeling at 0-1 h (at the single-cell stage) and 2-3 h (after PGC cellularization) after egg laying (AEL) (Figure 7A) and detected desthiobiotinylation restricted to the germ plasm, a specific cytoplasm at the posterior pole of embryos that contains germ granules (Figure 7A-B).

Then, we profiled local translation in the vicinity of germ granules with APEX-Ribo-Seq at each developmental stage (Figure S8B-D and Table S3). Accurate embryonic staging was confirmed by the increased translation of zygotic genes ^166^ at 2– 3 h AEL relative to 0–1 h AEL in the input Ribo-Seq data (Figure S8E). Calculating the read enrichment in germ granule APEX-Ribo-Seq relative to the input Ribo-Seq (Figure S8F), we identified 1348 and 1453 genes that were translated at 0-1 h and 2-3 h AEL, respectively (Figure 7C). Importantly, our data recapitulated previously characterized germ granule mRNAs ^7^: *nos*, *gcl*, and *pgc* were all enriched in the pulldown fraction at both stages (Figure 7C). Although *cyclin B* mRNA is well known to be recruited to germ granules, its translation is repressed ^167,168^. Consistent with this, *cyclin B* was significantly depleted from our germ granule APEX-Ribo-Seq at both stages (Figure 7C).

Next, we investigated the dynamics of local translation at the germ granule between the two stages, defining the difference in APEX-Ribo-Seq enrichment over time (Figure S8G). We detected diverse alterations of germ plasm local translation over time: “2-3 h AEL-biased” genes (*n* = 512) and “0-1 h AEL-biased” genes (*n* = 243) (Figure 7D). Strikingly, these dynamics were not mainly driven by changes in the mRNA species localized at the germ plasm; APEX–RNA-seq analysis of the same samples (Figure S8H) revealed relatively stable profiles over time (Figure 7E). Thus, while the mRNA content was largely maintained across the two stages, the temporal shift of on-site translational control was dynamically induced in germ granules.

We found a clear functional division of local translation shift between the two developmental windows (Figure 7F). STRING analysis showed that genes biased toward 0-1 h AEL were significantly enriched for components of the translational machinery, including the cytosolic ribosome (Figure 7F). In contrast, genes biased toward 2-3 h AEL were associated with pathways related to Piwi-interacting RNA (piRNA) biogenesis and carbon metabolism, particularly glycolysis (Figure 7F). Notably, glycolytic enzymes have been shown to localize to germ granules at the protein level specifically at 2–3 h AEL ^169^. This spatial and temporal coincidence suggests that not only these enzymes are components of germ granules, but their synthesis would also be increased in the vicinity of germ granules.

To investigate the mechanism for the temporal switch in granule translation, we focused on the translational control mediated by Aubergine (Aub), a PIWI family protein that binds to piRNAs. Aub plays an important function in silencing transposable elements in the germline through the piRNA pathway ^170–173^. However, several studies have also shown its role in mRNA regulation ^174–181^, in particular in the early embryo. Aub is involved in the decay of a group of maternal mRNAs in the somatic part of the embryo and in their localization and stabilization in germ granules ^172,174,176,181,182^. In addition, Aub activates the translation of *nos* mRNA ^180^. To address a more general role of Aub in translational activation at germ granules, we harnessed individual-nucleotide resolution CLIP-Seq (iCLIP-Seq) data of Aub obtained from 0-2 h AEL embryos ^174^. We observed that the presence of an Aub-binding cluster alone did not distinguish the translation status of mRNAs on germ granules (Figure 7G), in line with widespread Aub binding to maternal mRNAs ^174,181^. In contrast, the functional Aub targets that were destabilized in the soma and stabilized in germ granules ^174,176^ were strongly enriched in a group of germ granule mRNAs translation-biased at 2-3 h AEL than those at 0-1 h AEL (Figure 7H). Our data support the spatially distinct functions of Aub during early embryogenesis and point to a general function of Aub in translational activation at germ granules.

## Conclusion

In this study, we introduced APEX-Ribo-Seq as a robust and versatile platform for profiling local translation at subcellular compartment resolution, across diverse contexts in human tissue culture, mouse primary neurons, and fly embryos. After optimizing APEX2-mediated proximity labeling for ribosome and RNA purification, we surveyed translation across 14 distinct subcellular regions, 3 neuronal compartments, and the fly *in vivo* germ granule, revealing strong and dynamic connections between organelle/locus/compartment functions, their locally supplied proteins, and the contribution to cell fate specification. This comprehensive survey demonstrates the substantial advantages of APEX-Ribo-Seq over preexisting methods. First, this technique does not require the purification of organelles/loci/compartments *per se,* avoiding cell fixation or crosslinking, and uses a simple and common protocol for labeled complex purification that can be applied to any subcellular region. Second, our labeling and purification approach is compatible with RNA-Seq, allowing dissection of the contribution of translation control beyond RNA localization. This point is typically difficult to address through BirA–AviTag-mediated methods ^21,58,61,62^. Third, our technique has fine temporal resolution, with a labeling time of only 1 min (shorter than the 30 min of biotin exposure in TurboID-based PL-Ribo-Seq ^63^), enabling the capture of translation profiles before the diffusion of the mRNA ribosome complex to other subcellular regions.

The perspective provided by our transcriptome-wide, unbiased APEX-Ribo-Seq method complements recent single-molecule imaging approaches, such as ribosome-bound mRNA detection using proximity ligation and *in situ* sequencing (RIBOmap) ^51^ and high-throughput mRNA detection (multiplexed error-robust FISH [MERFISH] ^183^ and bacterial artificial chromosome [BAC]-based screening ^19^). The subcellular heterogeneity of ribosome distribution revealed by ribosome expansion microscopy (RiboExM) ^62^ and density-based localization monitoring for RNAs and proteins ^184^ could be integrated with our dataset for a deeper understanding of the molecular mechanism involved in local translation.

## Limitations of the study

Although APEX2 has provided successful results for local proteome ^66,67,73,76,185–189^ and transcriptome ^64,65,74^ identification, the reaction specificity in cells and the purity of the biotin-labeled complex *in vitro* require further optimization. Here, we carefully conducted enrichment analysis, considering both total input and cytosolic controls (Figure S2E). To achieve this normalization, 4 conditions (and their replicates) of Ribo-Seq/RNA-Seq are needed to calculate localization enrichment for one target. Due to the reaction diameter through APEX2 ^64,65,67,73–75^, local translation on tightly connected organelles (such as ER and mitochondria) could not be exclusively addressed. It also carries a risk of contaminating signals from unwanted mRNA–ribosome complexes at nearby loci. The enrichment analysis effectively suppresses this background.

Although here we took a conservative definition with a statistical cutoff and enrichment threshold based on ER with the earlier BirA–AviTag-mediated system ^58^ (Figure S2F), more references, ideally through orthogonal methods, will improve the accuracy and confidence of local translation likelihood for mRNAs. Recent surveys of OMM local translation in human cells, which were reported during the process of this work ^99,190^, complemented our findings of OMM translation, although these two studies may monitor mRNAs more closely located to OMM than those detected in the APEX2-based method. In addition, a local translation survey in more diverse loci that matches protein localization studies will resolve the inconsistency between the place of synthesis and the usage in some sub-cellular regions (Figures 2E, S5A, and S5B).

Because it is based on short reads, Ribo-Seq lacks the resolution to distinguish transcript isoforms, which may have distinct 5′ UTR and 3′ UTR sequences and possess different patterns of mRNA localization and translation control ^191,192^.

While our language model-based approach demonstrates that high-dimensional embeddings capture information beyond traditional features such as motif enrichment or region length, a current limitation is the incomplete understanding of the specific sequence determinants learned by the model. Precise decoding of the sequence features identified by TransLocAttn that correlate with RBP interactions will be an interesting next step. One strategy will be to enhance model interpretability based on methods adopted from natural language models, as has been recently applied in DNA models ^146^. Importantly, the modeling results should be supported experimentally to understand the causality, as exemplified by our comparison of attention and RBP binding profiles (Figure 5).

## Resource availability

### Lead contact

Requests for further information and resources should be directed to and will be fulfilled by the lead contact, Shintaro Iwasaki.

### Materials availability

The reagents, commercial assays, model organisms/strains, software, and equipment can be found in the key resources table. Further information and requests for resources and reagents should be directed to and will be available upon completion of appropriate material transfer agreements by the lead contact, Shintaro Iwasaki.

### Data and code availability

- The Ribo-Seq, APEX-Ribo-Seq, RNA-Seq, and APEX-RNA-Seq data obtained in this study (BioProject accession number: PRJDB16777) were deposited in the DNA Data Bank of Japan. This study also used previously deposited Ribo-Seq and RNA-Seq data GSE304445 [https://www.ncbi.nlm.nih.gov/geo/query/acc.cgi?acc=GSE304445] ^72^).
- Custom scripts for the mapping and quantification of ribosome profiling are available at https://github.com/ingolia-lab/RiboSeq. The other key custom scripts used in this study are available at Zenodo (DOI: 10.5281/zenodo.15484603 [https://zenodo.org/records/15484603]).
- Any additional information required to reanalyze the data reported in this paper is available from the lead contact upon request.

## Acknowledgments

We are grateful to all the members of the Iwasaki laboratory, Dr. Kosuke Dodo, Dr. Yutaka Ogawa, Dr. Fubito Nakatsu, Dr. Franz Meitinger, Dr. Midori Ota, Dr. Akihiko Nakano, Dr. Tomoko Kawamata, Dr. Hotaka Kobayashi, Dr. Benoit Palancade, Dr. Martine Simonelig, and Dr. Kenji Ohtawa, for their constructive discussions and technical help. A portion of the deep sequencing was performed at the Advanced Genomics Center, National Institute of Genetics (supported by the Japan Society for the Promotion of Science [JSPS], JP22H04925). Computation was supported by the HOKUSAI SailingShip supercomputer facility at RIKEN. We are grateful to the Support Unit for Bio-Material Analysis, RIKEN CBS Research Resources Division, for Sanger sequencing and flow cytometry analysis. Fly stocks obtained from the Bloomington Drosophila Stock Center (NIH P40OD018537) were used in this study. This work was supported by the Ministry of Education, Culture, Sports, Science and Technology (MEXT) (JP24H02307 to S.I.; JP21H05734, JP23H04268, and JP25H01440 to Y.S.; JP24H02327 to T.F.), the Japan Agency for Medical Research and Development (AMED) (JP20gm1410001 to S.I.; JP23gm6910005 to Y.S.), JSPS (JP26H02372 and JP23H00095 to S.I.; JP21K15023 and JP23K05648 to Y.S.; JP23H00095 to K. Kawaguchi; JP26K09203 to S.M.; JP25H00967 to T.F.), the Japan Science and Technology Agency (JST) (JPMJCR25T2 to T.F., Y.S., and S.I.; JPMJBS2418 and JPMJAX253D to K.T.; JPMJFR214W to T.F.), the Japan Science Society (the Sasakawa Scientific Research Grant to K.T.), the Nakajima Foundation (to Y.S. and S.M.), the Sumitomo Foundation (to S.M.), the Inamori Foundation (to S.M.), the Exploratory Research Center on Life and Living Systems (ExCELLS) (23EXC601-2 and 25EXC602-4 to Y.S.; 23EXC601 and 25EXC602 to N.S.), and RIKEN (Pioneering Project to S.I. and Y.S.; RIKEN TRIP initiative “TRIP-AGIS” to S.I.). K.T. was supported by a Junior Research Associate Program (JRA) from RIKEN, the World-leading Innovative Graduate Study Program in Proactive Environmental Studies (WINGS-PES) from The University of Tokyo, and Broadening Opportunities for Outstanding young researchers and doctoral students in STrategic areas (BOOST) from JST.

## Author contributions

Conceptualization, K.T., Y.S., and S.I.;

Methodology, K.T., S.M., T.F., K. Kawaguchi, and Y.S.;

Formal analysis, K.T., K. Kawaguchi, and Y.S.;

Investigation, K.T., S.M., T.A., A.H., K. Kobayashi, T.F., K. Kawaguchi, and Y.S.;

Writing – Original Draft, K.T., Y.S., and S.I.;

Writing – Review & Editing, K.T., S.M., T.A., A.H., K. Kobayashi, N.S., T.F., K. Kawaguchi, Y.S., and S.I.;

Visualization, K.T.;

Supervision, N.S., T.F., K. Kawaguchi, Y.S., and S.I.;

Project administration, K. Kawaguchi, Y.S., and S.I.;

Funding Acquisition, K.T., S.M., T.F., K. Kawaguchi, N.S., Y.S., and S.I.

## Declaration of generative AI and AI-assisted technologies in the writing process

During the preparation of this work, the authors used ChatGPT and Claude in order to refine the description. After using this tool/service, the authors reviewed and edited the content as needed and take full responsibility for the content of the publication.

---

**Figure S1.**
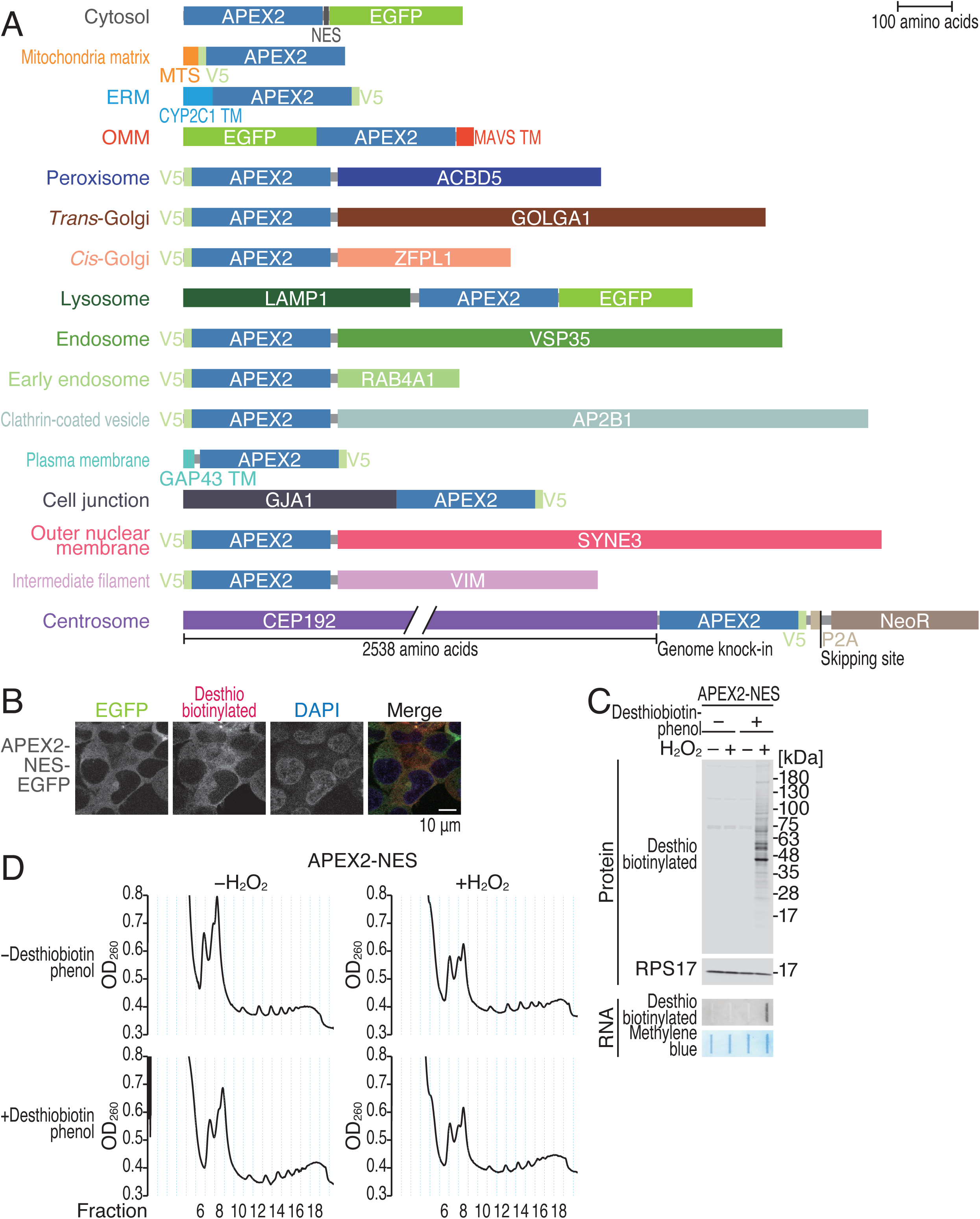
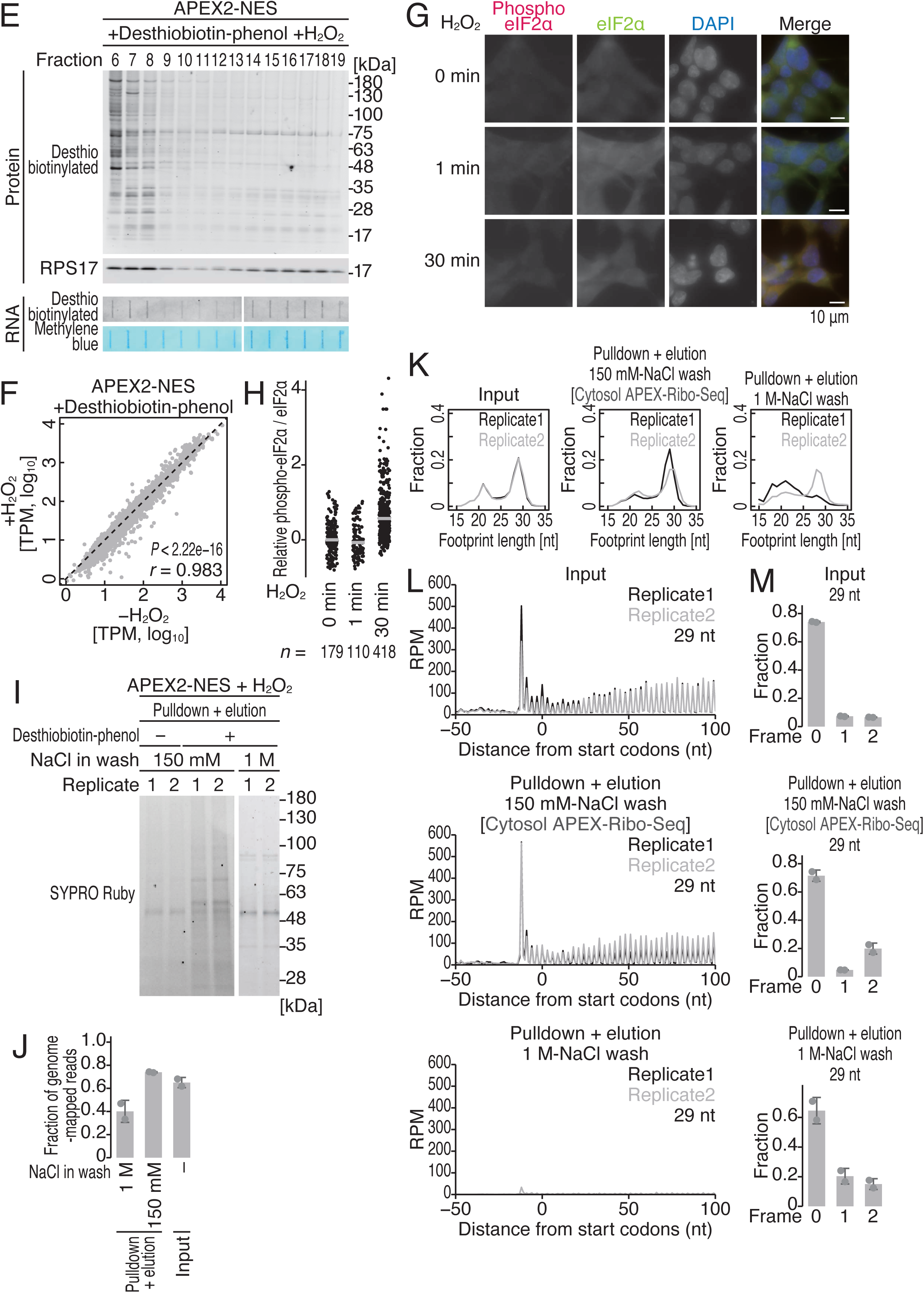
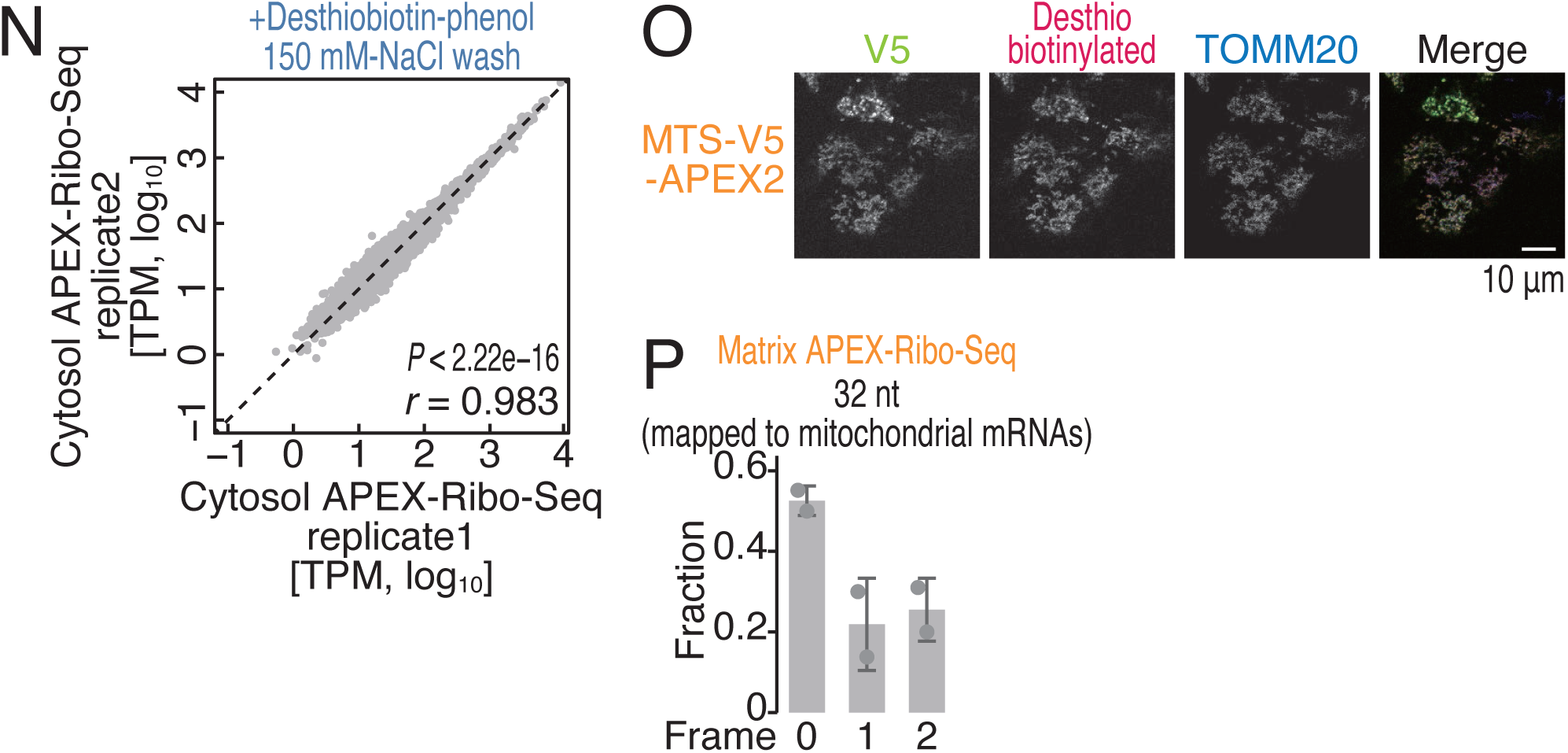
Optimization of the APEX-Ribo-Seq protocol, related to Figure 1. (A) Schematics of constructs for APEX2 fusion protein expression. (B) Immunofluorescence analysis of the indicated proteins (detected by EGFP) and desthiobiotinylated proteins/RNAs (detected with Alexa Fluor 555-conjugated streptavidin). DAPI (4’,6-diamidino-2-phenylindole) was used to stain the DNA. The scale bar represents 10 μm. (C) Western blotting and RNA dot blotting for proteins and RNAs under the indicated conditions. Desthiobiotinylated proteins and RNAs were detected using infrared dye-conjugated streptavidin. As a loading control, RPS17 was subjected to Western blotting, and total RNA was stained with methylene blue. (D) Polysome profiling under the indicated conditions. (E) Western blotting and RNA dot blotting for proteins and RNAs along the fractions of polysome profiling (see D for the corresponding fraction numbers). Desthiobiotinylated proteins and RNAs were detected by infrared dye-conjugated streptavidin. Total RNA was stained with methylene blue. (F) Scatter plot of the ribosome footprints from each mRNA in standard Ribo-Seq. The results in the presence and absence of H_2_O_2_ were compared. *P* values were calculated with Pearson’s product-moment correlation test (two-sided). (G) Immunofluorescence analysis of eIF2α and phosphorylated (phospho) eIF2α during H_2_O_2_ treatment. DAPI was used to stain the DNA. The scale bar represents 10 μm. (H) Quantification of the relative level of phosphorylated (phospho) eIF2α over total eIF2α is shown. The median (bars) and individual data (points; n for the number of cells analyzed) are shown. (I) SYPRO Ruby staining of proteins purified after streptavidin pulldown and elution under the indicated conditions. (J) Fractions of genome-mapped reads (reads after deduplication and removal of noncoding RNA-mapped reads) in standard Ribo-Seq and cytosol APEX-Ribo-Seq under the indicated conditions. The means (bars), s.d.s (errors), and individual replicates (n = 2, points) are shown. (K) Distribution of ribosome footprint length under the indicated conditions. (L) Metagene plots of ribosome footprints (the 5′-end positions) around the start codon under the indicated conditions. The data for the 29-nt footprints are shown. (M) Fraction of the frame position for the 5′ end of the footprint (29 nt) under the indicated conditions. The means (bars), s.d.s (errors), and individual replicates (n = 2, points) are shown. (N) Scatter plot of the ribosome footprints from each mRNA in cytosol APEX-Ribo-Seq replicates. *P* values were calculated with Pearson’s product-moment correlation test (two-sided). (O) Immunofluorescence analysis of the indicated proteins (detected by V5) and desthiobiotinylated proteins/RNAs (detected with Alexa Fluor 555-conjugated streptavidin). TOMM20 was detected as a mitochondrial marker. The scale bar represents 10 μm. (P) Fraction of the frame position for the 5′ end of the mitoribosome footprint (32 nt) in matrix APEX-Ribo-Seq. The means (bars), s.d.s (errors), and individual replicates (n = 2, points) are shown. TPM, transcripts per million; *r*, Pearson’s correlation coefficient; RPM, reads per million reads.

**Figure S2.**
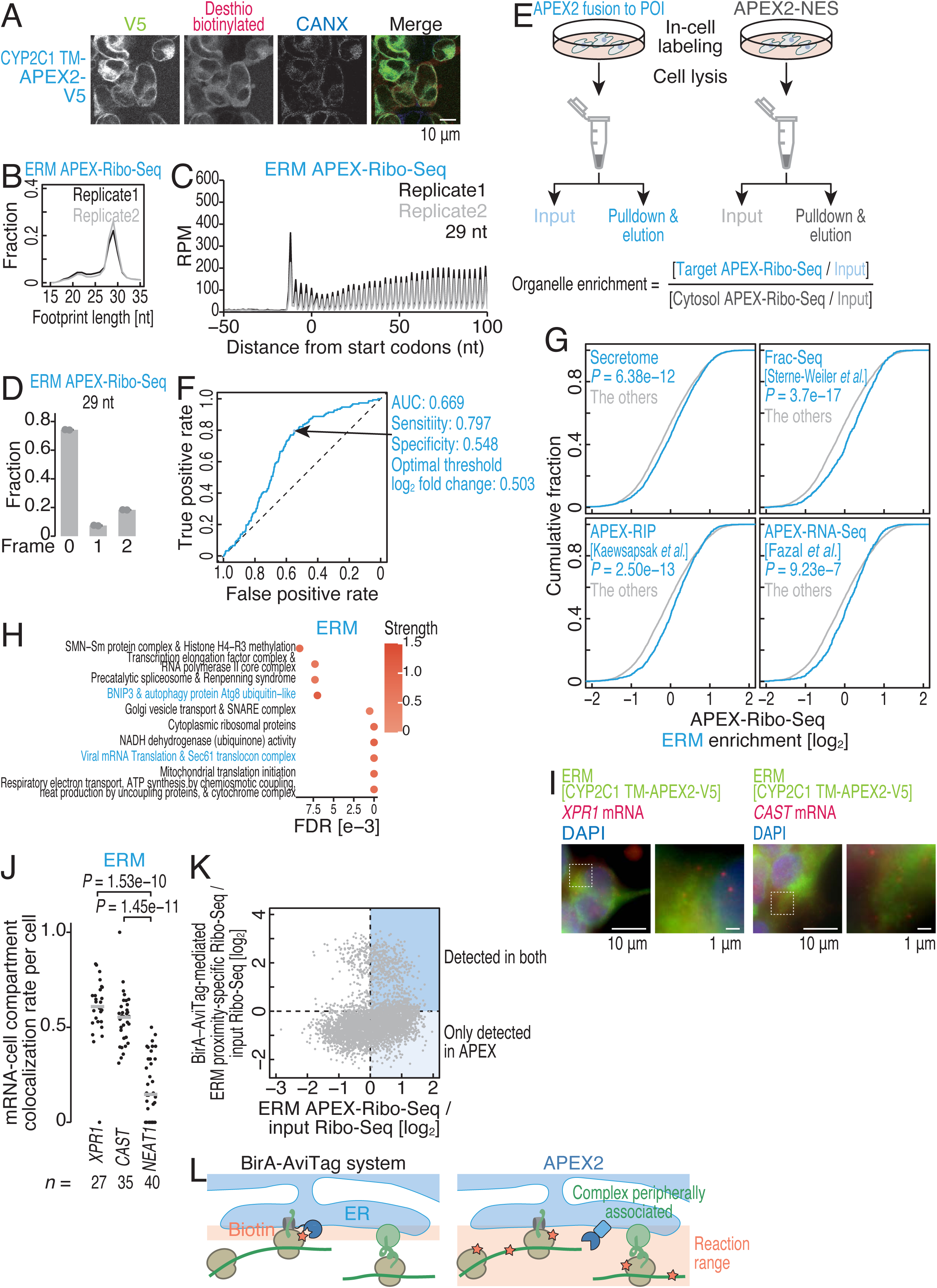
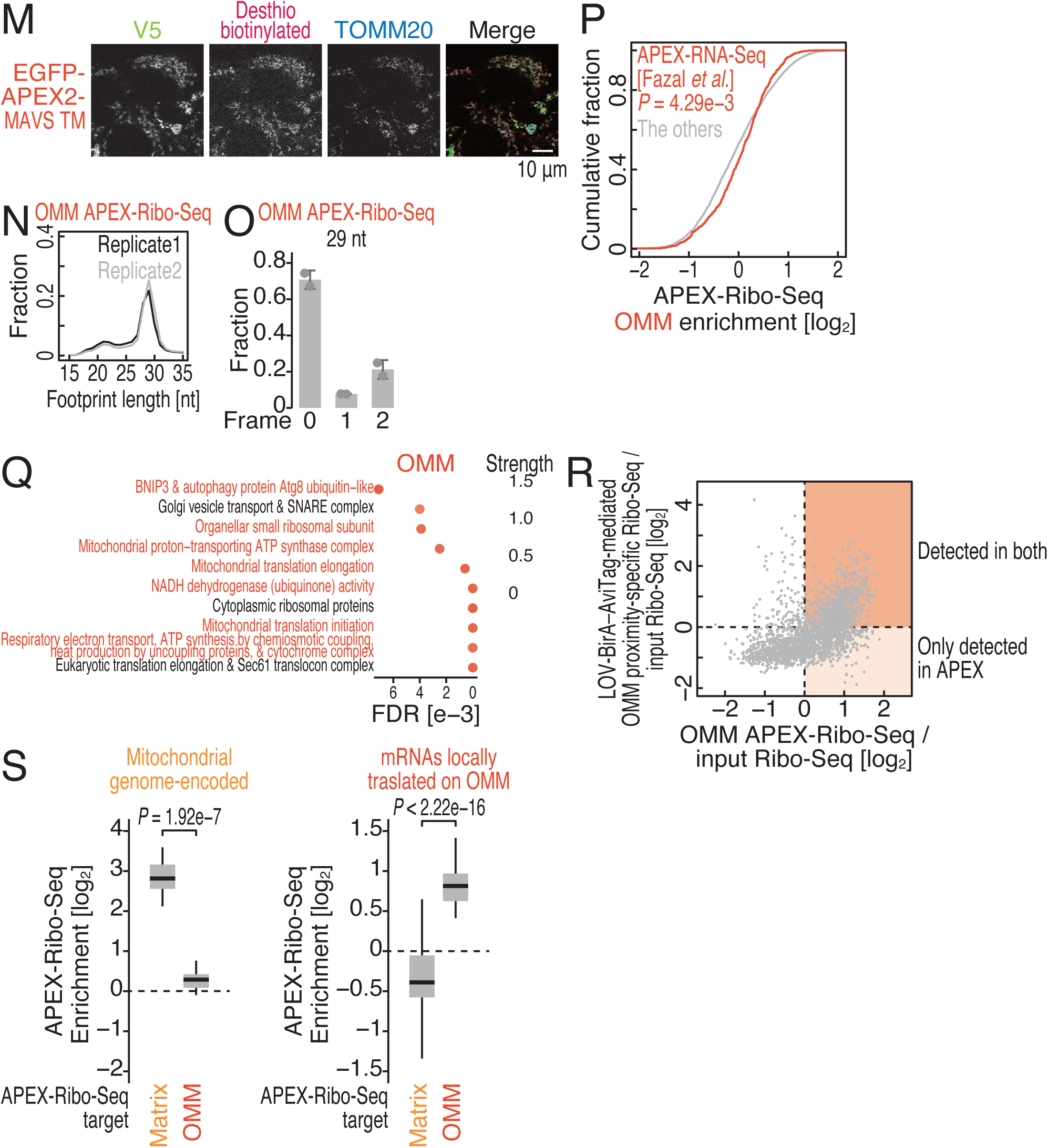
Characterization of ERM and OMM APEX-Ribo-Seq data, related to Figure 1. (A) Immunofluorescence analysis of the indicated proteins (detected by V5) with desthiobiotinylated proteins/RNAs (detected with Alexa Fluor 555-conjugated streptavidin). CANX was detected as a marker of the ER. The scale bar represents 10 μm. (B) Distribution of ribosome footprint length under the indicated conditions. (C) Metagene plots of ribosome footprints (the 5′-end positions) around the start codon under the indicated conditions. The data for the 29-nt footprints are shown. (D) Fraction of the frame position for the 5′ end of the footprint (29 nt) under the indicated conditions. The means (bars), s.d.s (errors), and individual replicates (n = 2, points) are shown. (E) Schematics of the organelle enrichment calculation. (F) ROC curve to determine the fold change threshold for organelle enrichment. (G) Cumulative distributions of ERM enrichment in APEX-Ribo-Seq with respect to the mRNAs found in the secretome (see Methods for details) and by Frac-Seq ^91^, APEX-RIP ^66^, and APEX-RNA-Seq ^64^ targeting the ERM. *P* values were calculated with the Mann Whitney *U* test (two-tailed). (H and Q) Complexes encoded by mRNAs locally translated at the indicated organelles were assessed by STRING analysis ^93^. The results of the top 10 terms, ranked by false discovery rate (FDR) value, are visualized. The terms shown in color are described in the main text. The color scale indicates the strength of the associations in the network (ranging from 0 to 1), which is a confidence indicator ^93^. (I) Immunofluorescence analysis of the indicated proteins (detected by V5) with smiFISH for the indicated mRNAs. DAPI was used to stain the DNA. An enlarged view of the region indicated by the dashed square is shown. The scale bar represents 10 μm. (J) Quantification of the colocalization rates of the marker proteins and the indicated RNAs is shown. Nuclear paraspeckle *NEAT1* RNA was detected and analyzed as a negative control. The median (bars) and individual data (points; n for the number of cells analyzed) are shown. *P* values were calculated with the Mann Whitney *U* test (two-tailed) and adjusted by the Benjamini–Hochberg method. (K) Scatter plot of ERM APEX-Ribo-Seq enrichment over input Ribo-Seq vs. the BirA–AviTag-mediated ERM proximity-specific Ribo-Seq enrichment over input Ribo-Seq ^58^. mRNAs detected in both approaches and mRNAs detected in only APEX-Ribo-Seq are highlighted. (L) Schematics of the reaction range difference in the BirA–AviTag-mediated system and the APEX2-mediated method. (M) Immunofluorescence analysis of the indicated proteins (detected by V5) with desthiobiotinylated proteins/RNAs (detected with Alexa Fluor 555-conjugated streptavidin). TOMM20 was detected as a mitochondrial marker. The scale bar represents 10 μm. (N) Distribution of ribosome footprint length under the indicated conditions. (O) Fraction of the frame position for the 5′ end of the footprint (29 nt) under the indicated conditions. The means (bars), s.d.s (errors), and individual replicates (n = 2, points) are shown. (P) Cumulative distributions of OMM enrichment in APEX-Ribo-Seq with respect to the mRNAs found in APEX-RNA-Seq ^64^ targeting the OMM. *P* values were calculated with the Mann Whitney *U* test (two-tailed). (R) Scatter plot of OMM APEX-Ribo-Seq enrichment over input Ribo-Seq vs. LOCL-TL-mediated OMM proximity-specific Ribo-Seq enrichment over input Ribo-Seq ^99^. mRNAs detected in both approaches and mRNAs detected in only APEX-Ribo-Seq are highlighted. (S) Box plots of matrix and OMM enrichment (Figure S2E) determined via APEX-Ribo-Seq for mitochondrial genome-encoded mRNAs (left) and mRNAs locally translated on the OMM (right, defined in Figure 1F). *P* values were calculated with the Mann Whitney *U* test (two-tailed). RPM, reads per million reads. Box plots show the median (centerline), upper/lower quartiles (box limits), and 1.5× interquartile range (whiskers).

**Figure S3.**
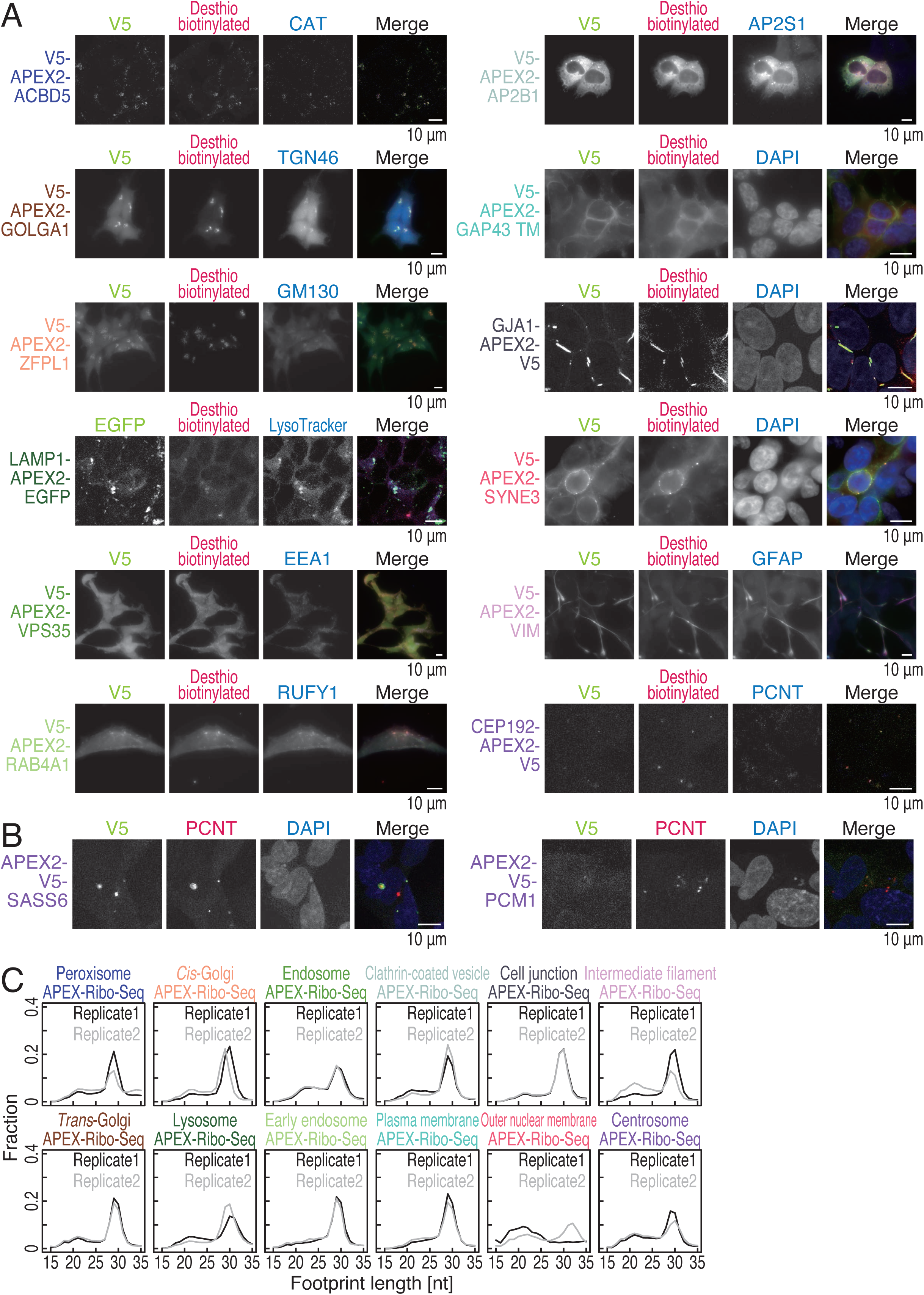
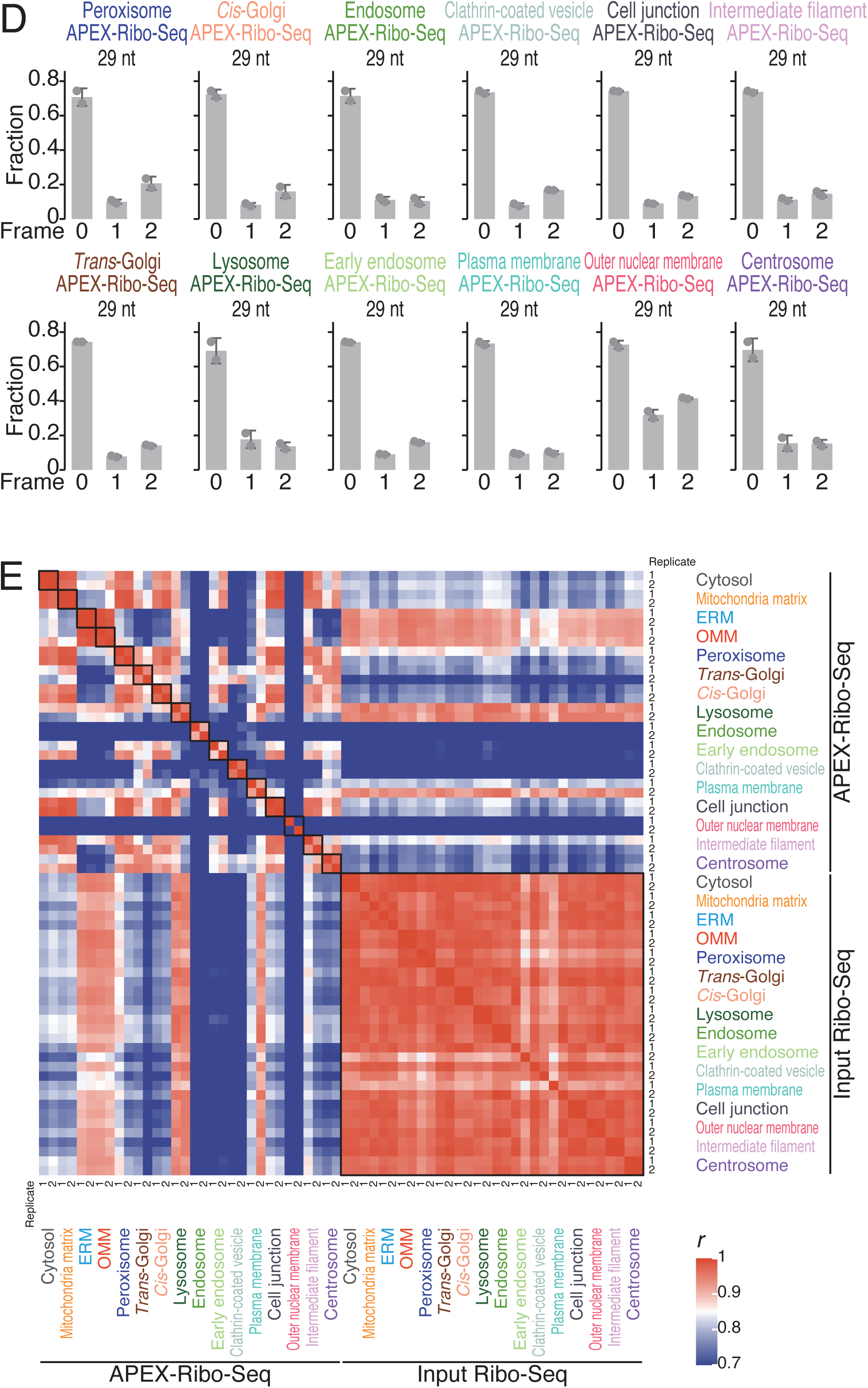
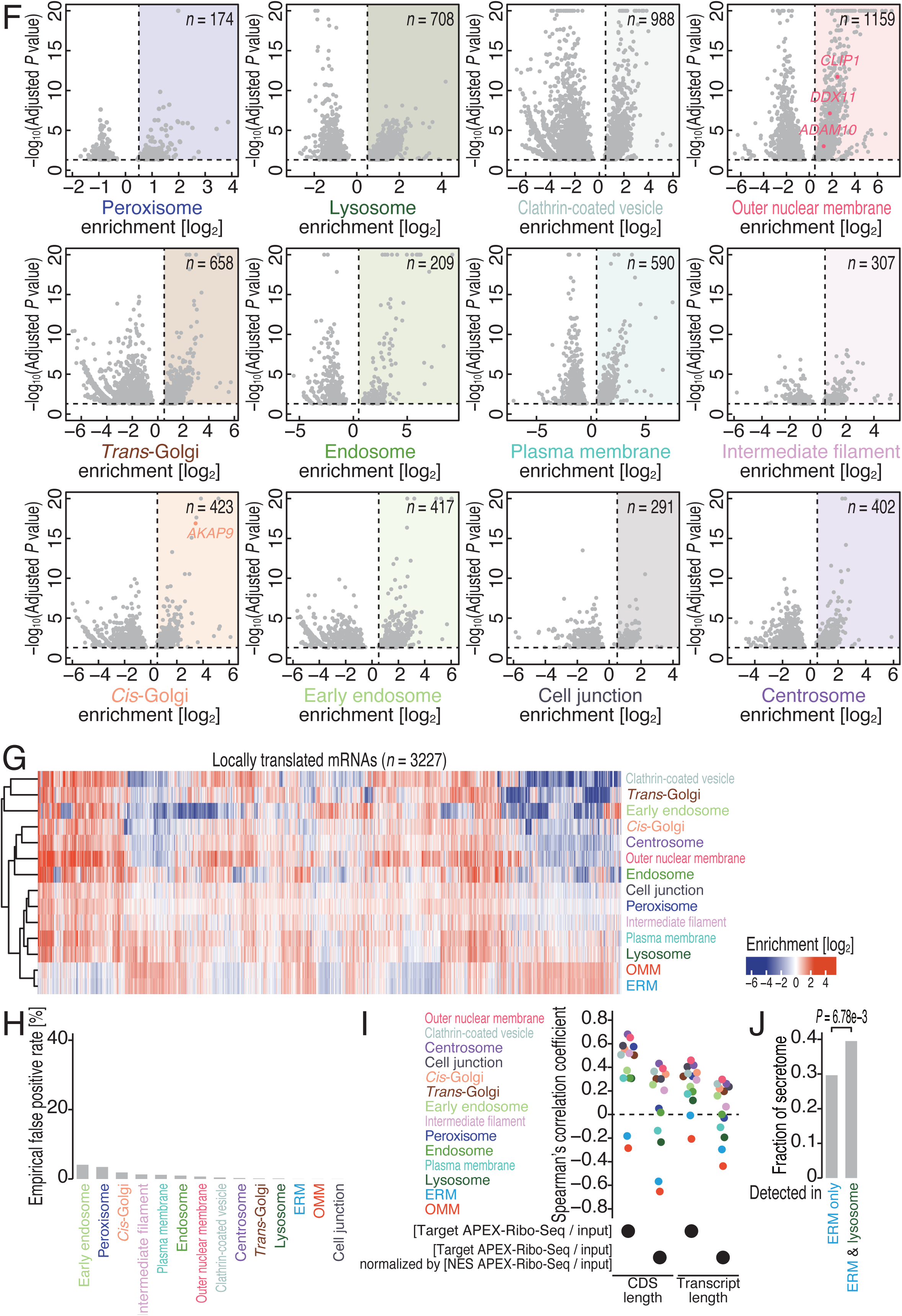
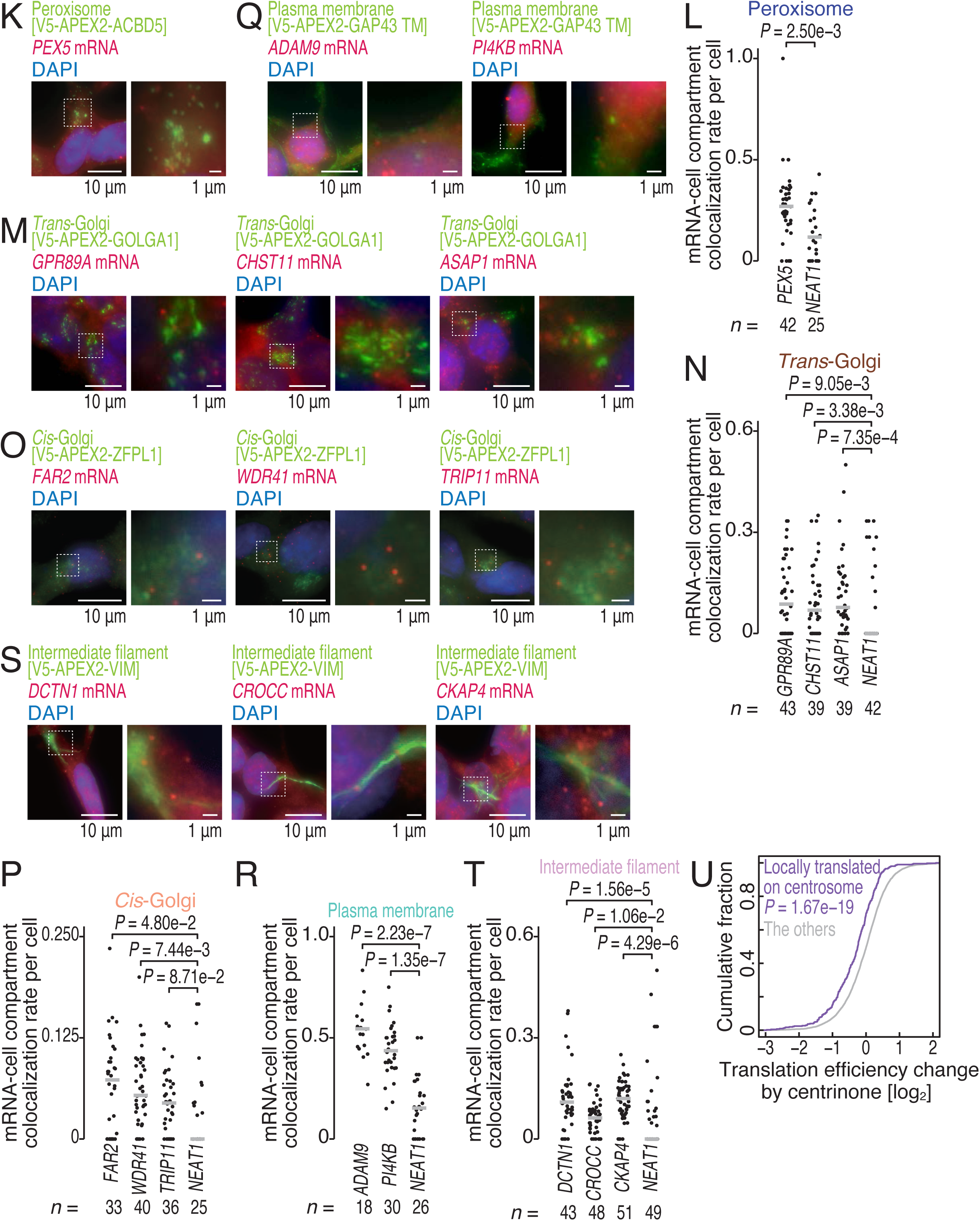
Characterization of APEX-Ribo-Seq data for 12 loci, related to Figure 2. (A) Immunofluorescence analysis of the indicated proteins (detected by EGFP or V5) with desthiobiotinylated proteins/RNAs (detected with Alexa Fluor 555-conjugated streptavidin). The following proteins were detected as organelle markers: catalase (CAT) (peroxisomes), TGN46 (*trans*-Golgi), GM130 (*cis-*Golgi), EEA1 (endosomes), RUFY1 (early endosomes), AP2S1 (clathrin-coated vesicles), GFAP (intermediate filaments), and PCNT (centrosomes). DAPI was used to stain the DNA. LysoTracker was used to stain lysosomes. The scale bar represents 10 μm. (B) Immunofluorescence analysis of the indicated proteins (detected by V5). PCNT was detected as a centrosomal marker. The white boxes highlight centrosomes with aberrant architecture. The scale bar represents 10 μm. (C) Distribution of ribosome footprint length under the indicated conditions. (D) Fraction of the frame position for the 5′ end of the footprint (29 nt) under the indicated conditions. The means (bars), s.d.s (errors), and individual replicates (n = 2, points) are shown. (E) Heatmap of the Pearson correlation coefficients (*r*) of APEX-Ribo-Seq and input standard Ribo-Seq under the indicated conditions. The value of *r* is indicated by the color scale. (F) Volcano plots showing organelle enrichment determined by APEX-Ribo-Seq (Figure S2E). mRNAs with an adjusted *P* value of 0.05 or lower are shown. The colored areas represent mRNAs that met the fold change threshold defined by ROC analysis (Figure S2F). The genes mentioned in the main text are highlighted. (G) Heatmap of organelle enrichment determined by APEX-Ribo-Seq (Figure S2E) for mRNAs locally translated at least one site in 14 organelles. The color scale indicates the degree of organelle enrichment. (H) How frequently proteins cytosolically translated are falsely called at each locus (or false-positive rate, FPR) was assessed. For this purpose, we first defined a cytosolic reference through the comparison of cytosolic APEX-Ribo-Seq (APEX-NES) to the input (log2[(cytosol pulldown/input)]), and defined 230 genes enriched. Then, we computed the fraction of those genes in the gene set of local translation at each locus. (I) Spearman’s correlation coefficient between CDS/transcript length vs. APEX-Ribo-Seq enrichment over input Ribo-Seq, or further normalized by NES APEX-Ribo-Seq over input Ribo-Seq (*i.e.*, organelle enrichment, Figure S2E). (J) Fraction of mRNAs defined as the secretome (see Methods for details) among mRNAs whose local translation is found both in ERM and lysosome or ERM only (defined in Figure S3G). (K, M, O, Q, and S) Immunofluorescence analysis of the indicated proteins (detected by V5) with smiFISH for the indicated mRNAs. DAPI was used to stain the DNA. An enlarged view of the region indicated by the dashed square is shown. The scale bar represents 1 or 10 μm. (L, N, P, R, and T) Quantification of the colocalization rates of the marker proteins and the indicated RNAs is shown. Nuclear paraspeckle *NEAT1* RNA was detected and analyzed as a negative control. The median (bars) and individual data (points, n for the number of cells analyzed) are shown. *P* values were calculated with the Mann Whitney *U* test (two-tailed) and adjusted by the Benjamini–Hochberg method. (U) Cumulative distributions of translation efficiency changes (measured by standard Ribo-Seq and RNA-Seq) with respect to the mRNAs locally translated on the centrosome (defined in Figure S3F). *P* values were calculated with the Mann Whitney *U* test (two-tailed).

**Figure S4.**
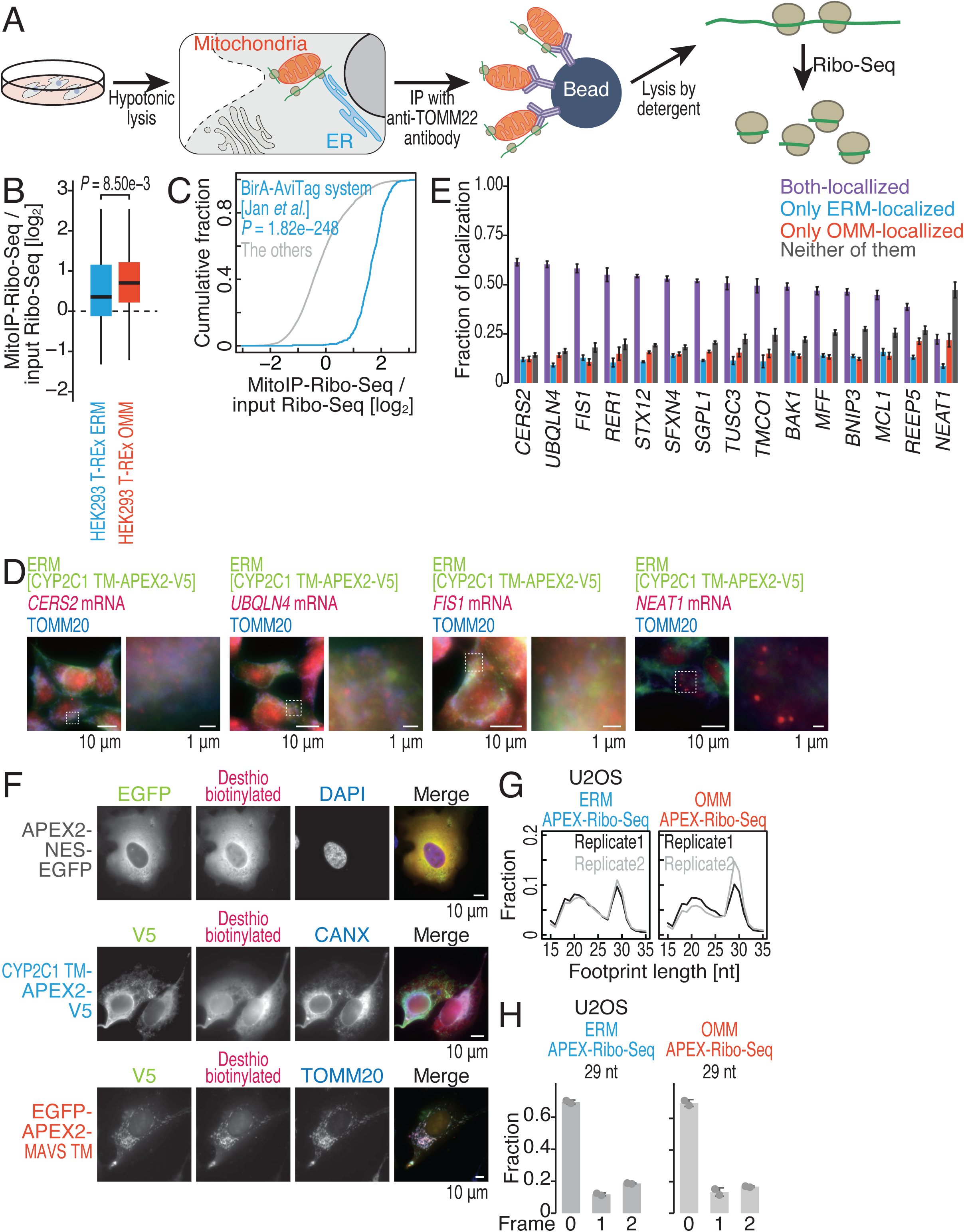
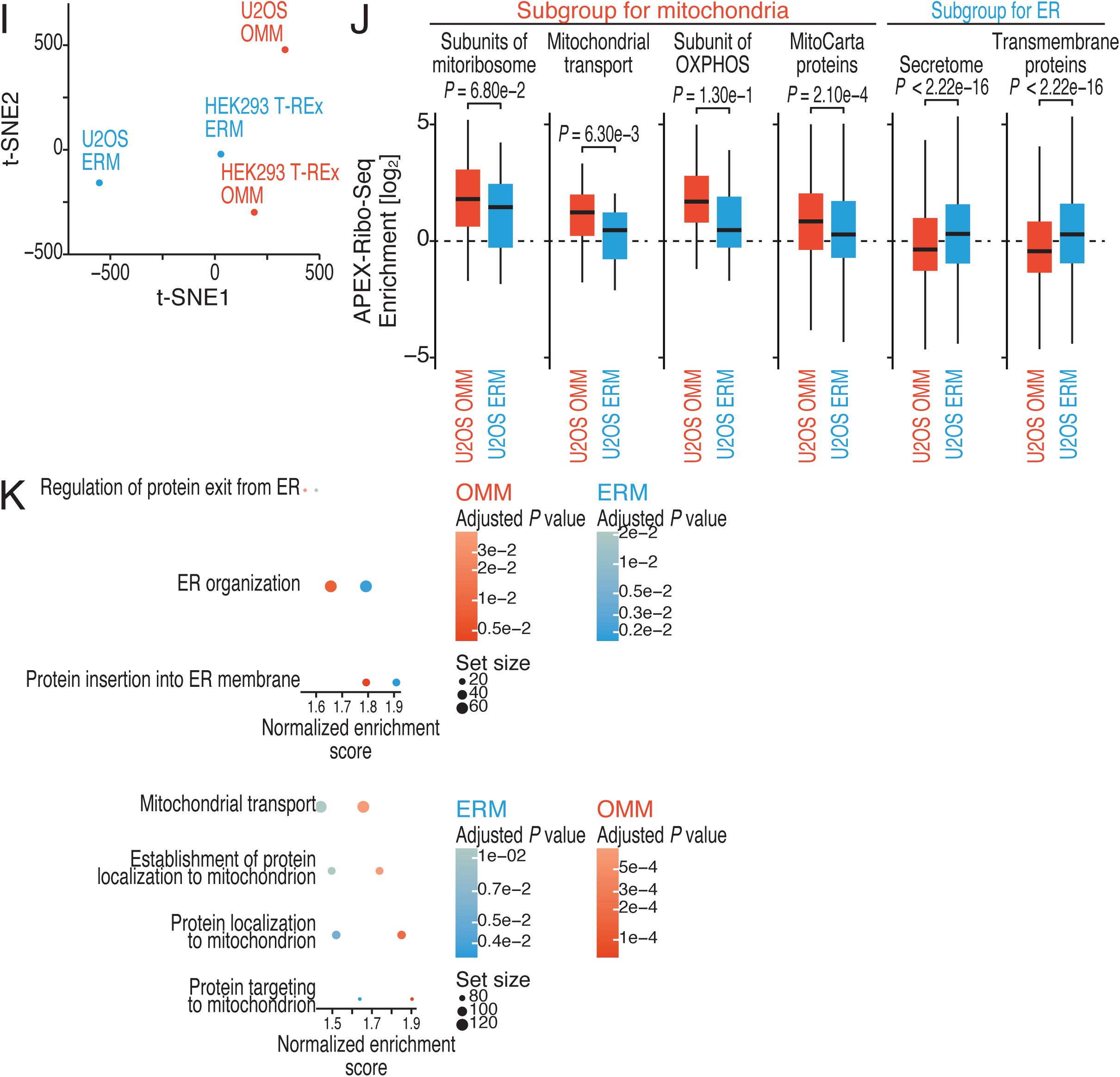
Characterization of ERM and OMM APEX-Ribo-Seq in U2OS, related to Figure 2. (A) Schematic of MitoIP-Ribo-Seq. (B) Box plots of MitoIP-Ribo-Seq enrichment over input Ribo-Seq for mRNAs locally translated on the ERM (defined in Figure 1E) and OMM (defined in Figure 1F). *P* values were calculated with the Mann Whitney *U* test (two-tailed). (C) Cumulative distributions of MitoIP-Ribo-Seq enrichment over input Ribo-Seq with respect to the mRNAs found in the BirA–AviTag-mediated system ^58^, targeting the ERM. *P* values were calculated with the Mann Whitney *U* test (two-tailed). (D) Immunofluorescence analysis of the indicated proteins (detected by V5) with smiFISH for the indicated mRNAs. DAPI was used to stain the DNA. An enlarged view of the region indicated by the dashed square is shown. The scale bar represents 1 or 10 μm. (E) Quantification of the colocalization rates of the marker proteins and the indicated RNAs is shown. Nuclear paraspeckle *NEAT1* RNA was detected and analyzed as a negative control. RNA spots were categorized into both ERM-OMM localized, only ERM-localized, only OMM-localized, or neither of them. The median (bars) and s.d.s (errors) are shown. (F) Immunofluorescence analysis of the indicated proteins (detected by V5 or EGFP) with desthiobiotinylated proteins/RNAs (detected with Alexa Fluor 555-conjugated streptavidin). CANX and TOMM20 were detected as markers of the ER and mitochondria, respectively. The scale bar represents 10 μm. (G) Distribution of ribosome footprint length under the indicated conditions. (H) Fraction of the frame position for the 5′ end of the footprint (29 nt) under the indicated conditions. The means (bars), s.d.s (errors), and individual replicates (n = 2, points) are shown. (I) t-SNE plot ^193^ for organelle enrichment (Figure S3G) determined by ERM and OMM APEX-Ribo-Seq in HEK293 T-REx cells and U2OS cells. (J) Box plots of ERM and OMM enrichment determined via APEX-Ribo-Seq in U2OS cells for the indicated mRNA categories. *P* values were calculated with the Mann Whitney *U* test (two-tailed). (K) GSEA analysis ^113^ for the indicated terms related to ER and mitochondrial functions. The color scale indicates the adjusted *P* value. Box plots show the median (centerline), upper/lower quartiles (box limits), and 1.5× interquartile range (whiskers).

**Figure S5.**
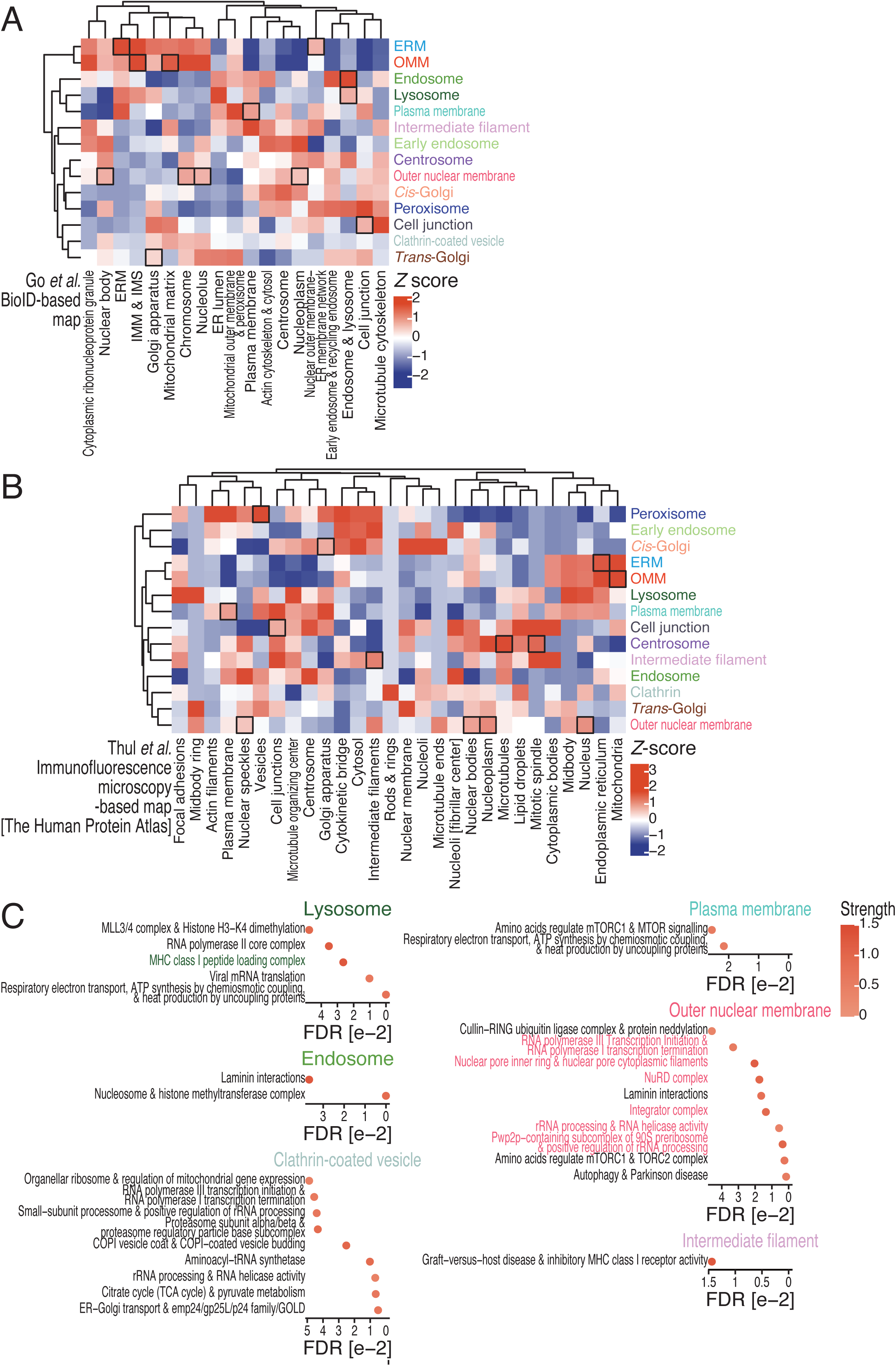
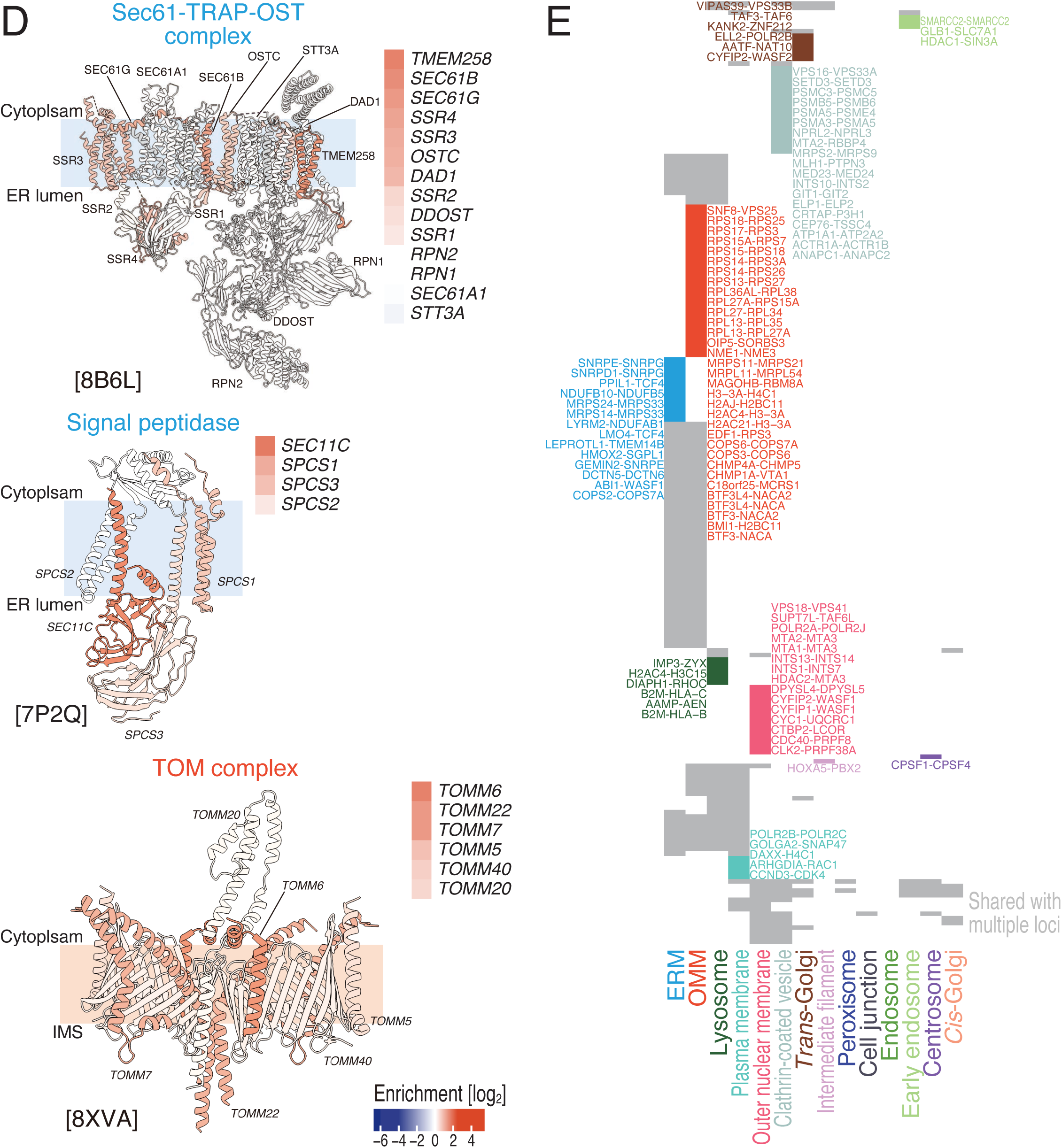
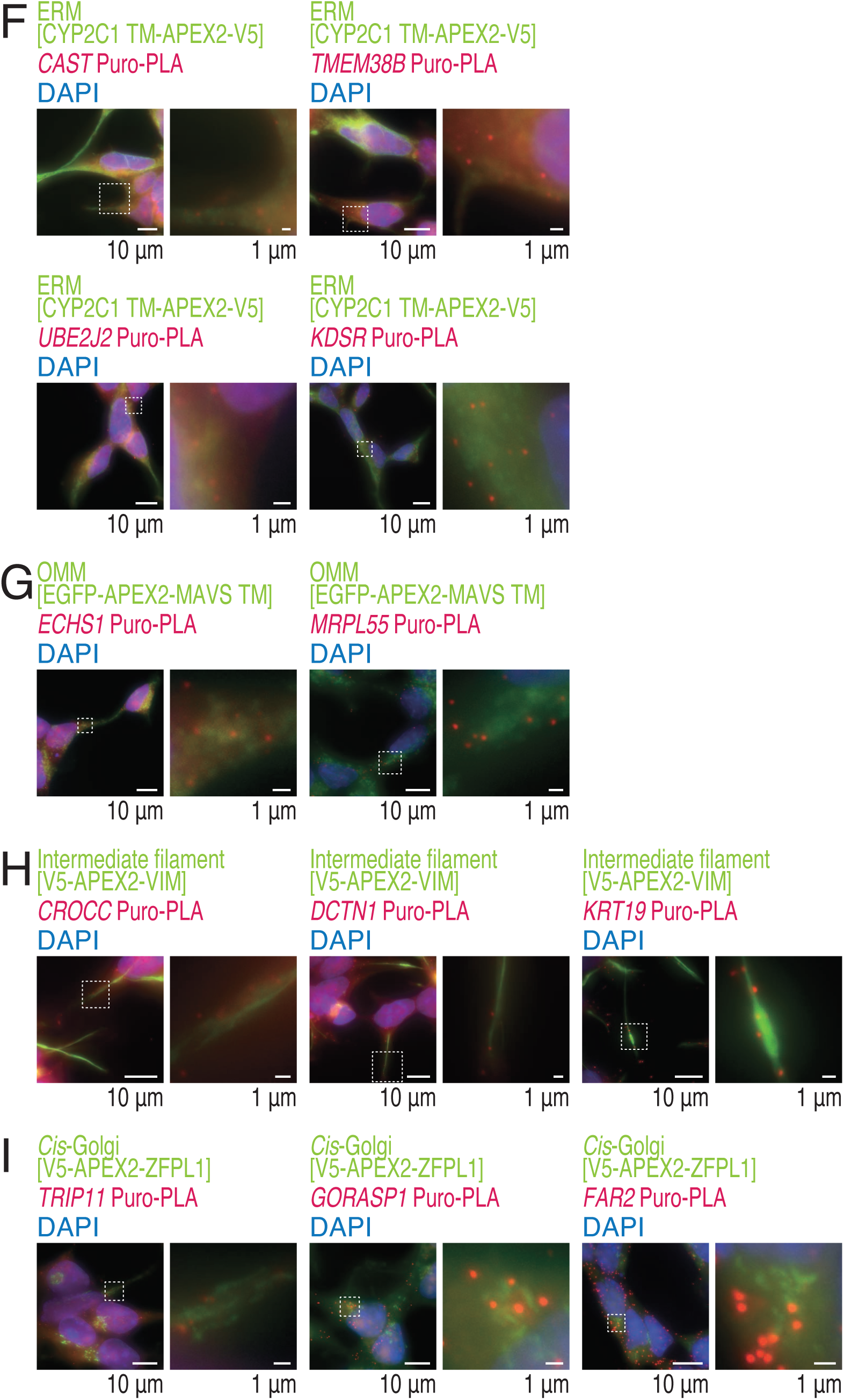
Comparative analysis of the local translation atlas with publicly available data, related to Figures 2-3. (A and B) Heatmap of the correspondence of locally translated mRNAs and the localization of proteins encoded in the mRNAs. The gene sets uniquely translated at each locus (Figure 2D) were used. Protein localization is defined by BioID-based proximity labeling ^100^ (Figure S5A) and immunofluorescence microscopy (The Human Protein Atlas) ^115^ (Figure S5B). The *Z* score represents the standardized enrichment or depletion of each localization term within the local translation category. The color scale indicates the *Z* score. The black box tiles are described in the main text. (C) Complexes encoded by mRNAs locally translated at the indicated organelles were assessed by STRING analysis ^93^. The gene sets uniquely translated at each locus (Figure 2D) were used. The results of the top 10 terms, ranked by false discovery rate (FDR) value, are visualized. The terms shown in color are described in the main text. The color scale indicates the strength of the associations in the network (ranging from 0 to 1), which is a confidence indicator ^93^. (D) Structural mapping of locally translated proteins at the indicated locus. The structures of the Sec61-TRAP-OST complex (Protein Data Bank [PDB]: 8B6L) ^116^, signal peptidase (PDB: 7P2Q) ^195^, and TOM complex (PDB: 8XVA) ^119^ are shown by UCSF Chimera X (ver. 1.1) ^194^. The color scale in the structure and heatmap of organelle enrichment determined by APEX-Ribo-Seq (Figure S3G) indicates the degree of organelle enrichment. (E) Tile plot showing the candidate co-co–assembled pairs ^129^ in the indicated mRNA groups. Pairs unique to each locus are highlighted. (F-I) Puro-PLA analysis with immunofluorescence analysis of the indicated proteins (detected by V5 or EGFP). DAPI was used to stain the DNA. An enlarged view of the region indicated by the dashed square is shown. The scale bar represents 1 or 10 μm. Quantification of the data is shown in Figure 3C-F.

**Figure S6.**
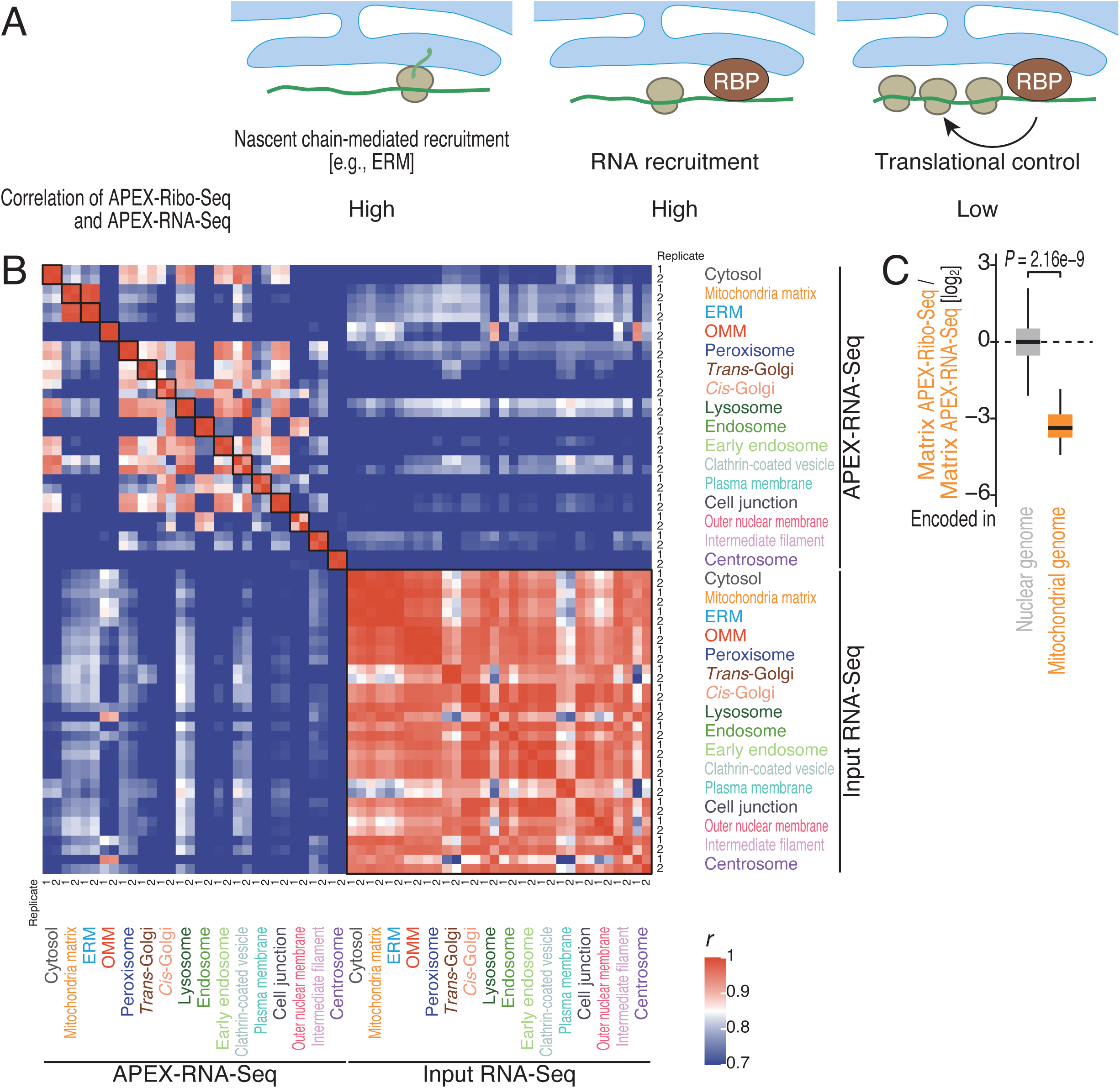
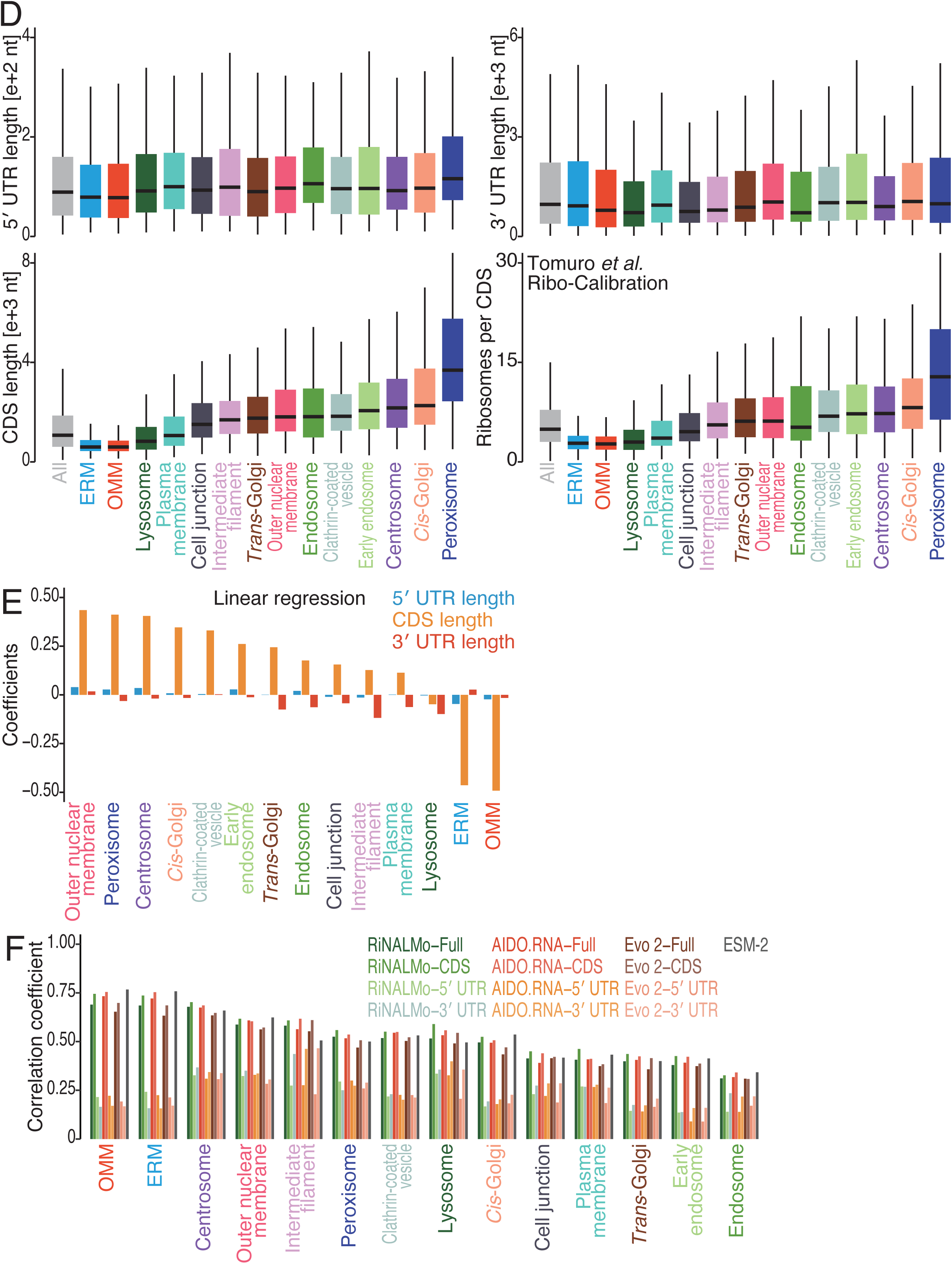
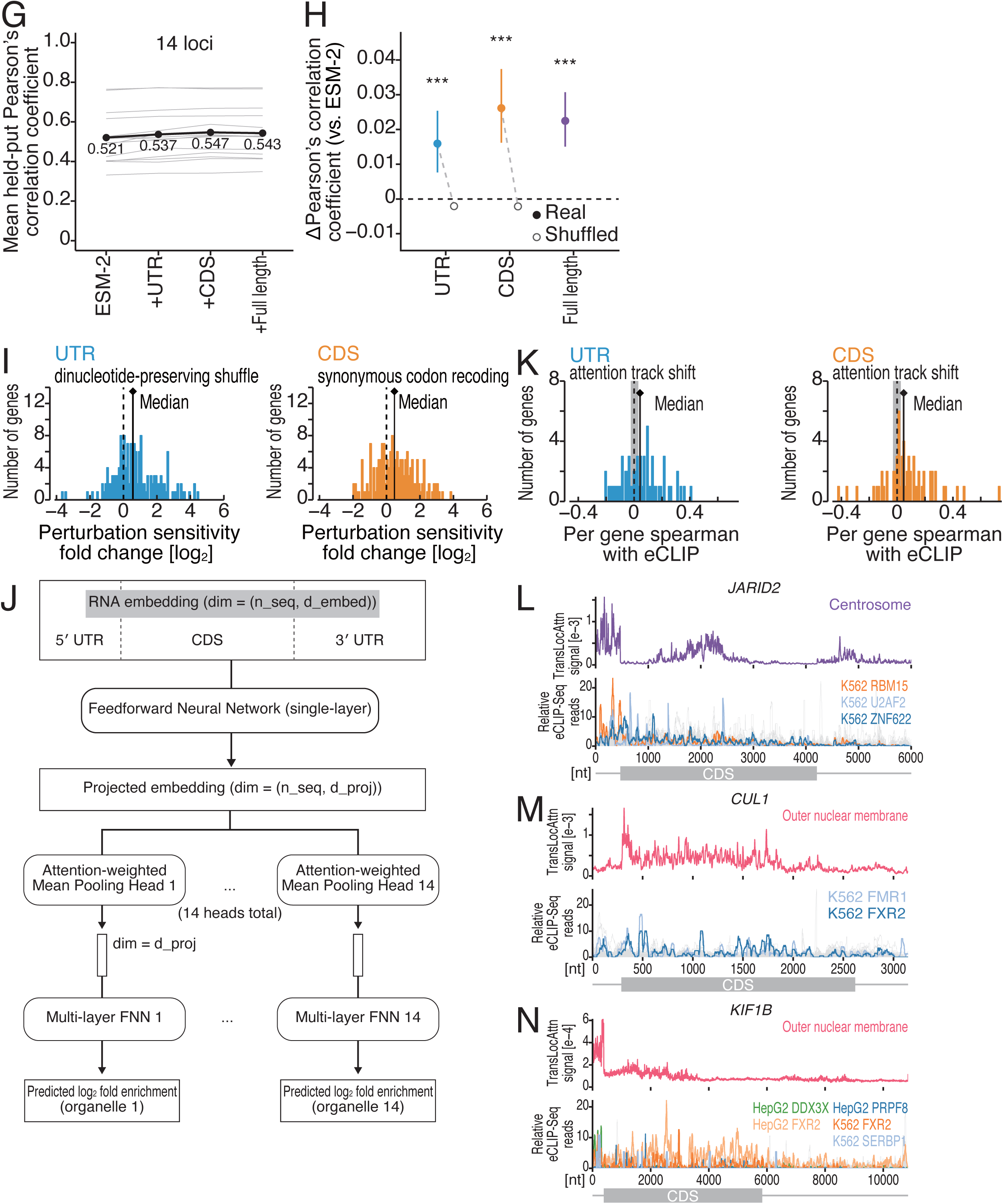
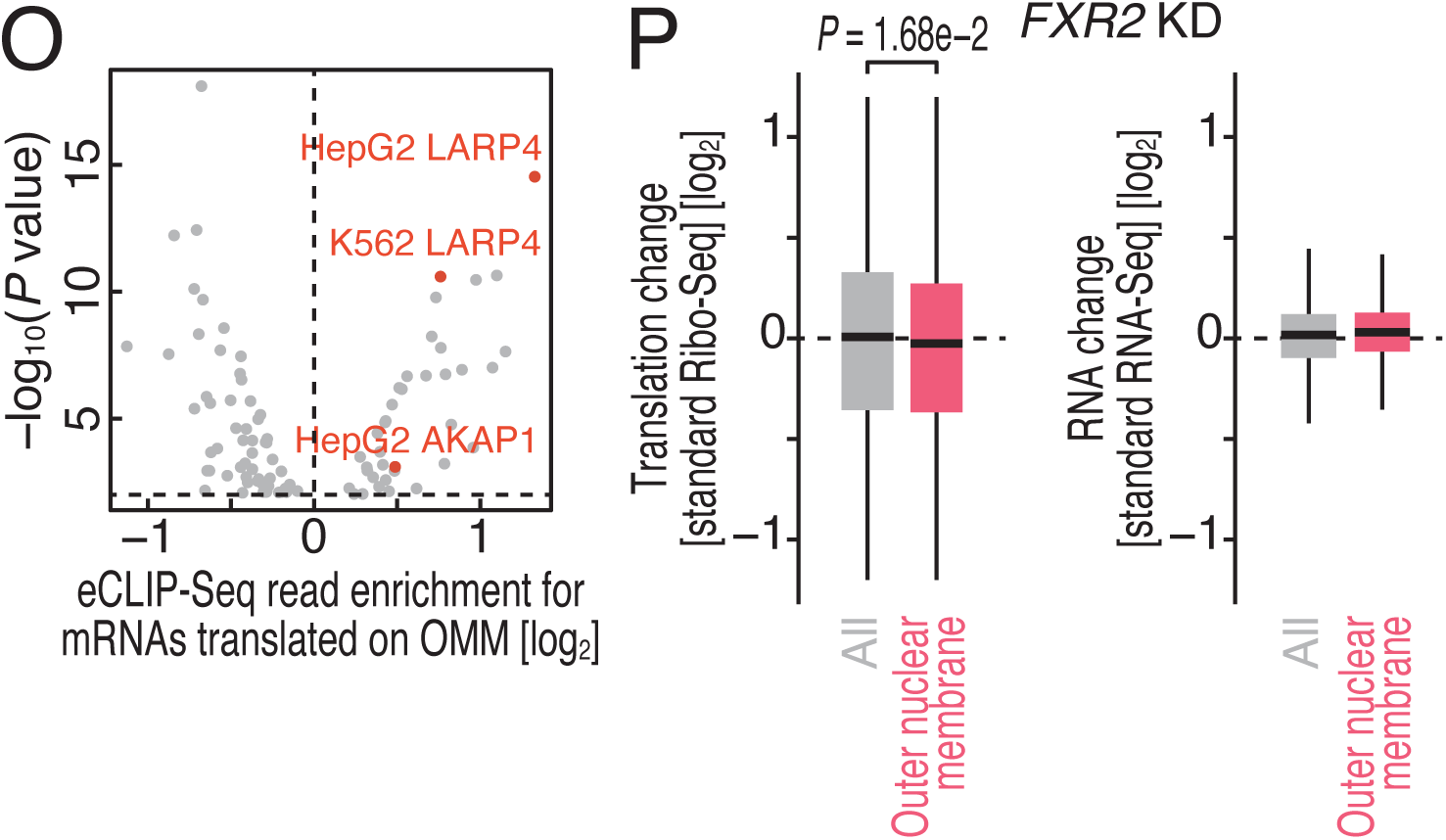
Application of the language model to the local translation atlas, related to Figures 4-5. (A) Schematic of the mechanisms for local translation and the correlation between APEX-Ribo-Seq and APEX-RNA-Seq. (B) Heatmap of the Pearson correlation coefficients (*r*) of APEX-RNA-Seq and input standard RNA-Seq under the indicated conditions. The value of *r* is indicated by the color scale. (C) Box plots of relative matrix translation efficiency (matrix APEX-Ribo-Seq normalized by matrix APEX-RNA-Seq) for the indicated mRNA groups. *P* values were calculated with the Mann Whitney *U* test (two-tailed). (D) Box plots of the 5′ UTR length (top left), the 3′ UTR length (top right), the CDS length (bottom left), and calibrated numbers of ribosomes on the CDSs, which were measured by Ribo-Calibration ^196^, for mRNAs translated on the indicated organelles. The gene sets uniquely translated at each locus (Figure 2D) were used. Ribosome numbers on CDS depend on the CDS length in general. (E) Coefficients from linear regression using all data, where the inputs are the lengths of each 5′ UTR, CDS, and 3′ UTR sequence, and the outputs are organelle enrichment determined by APEX-Ribo-Seq (log_2_ fold enrichment). (F) Pearson’s correlation between predicted and actual organelle enrichment determined by APEX-Ribo-Seq on the test set, where the models apply FNN to mean-pooled embeddings from the specified sequence region (full mRNA sequence, CDS, 5′ UTR, or 3′ UTR). RiNALMo, using embeddings generated by RiNALMo; AIDO.RNA, using embeddings generated by AIDO.RNA; Evo 2, using embeddings generated by Evo 2. For amino-acid sequence-based evaluation, embeddings generated by ESM-2 were used. (G) Held-out target-wise Pearson correlation between predicted and observed APEX-Ribo-Seq compartment enrichment, for the cross-fitted ESM-2 protein baseline and for that baseline combined with a UTR, CDS, or full-transcript RNA residual model, in the same 474-gene held-out set throughout. Thin lines, the 14 individual compartments; heavy line and circles, their mean. (H) Paired change in the same quantity relative to the ESM-2 baseline. Filled circles, real RNA representations; open circles, gene-shuffled controls that permute the RNA representation among genes within each split, run for the UTR and CDS features; bars, 95% target-block bootstrap intervals; ***, Benjamini-Hochberg-adjusted q < 0.001 across the three contrasts (one-sided target-block sign-flip test). (I) Per-gene ratio of the absolute change in residual model output for a rewritten high-attention window to that for a matched low-attention window in the same gene. Left, 50-nt UTR windows rewritten by dinucleotide-count-preserving shuffling; right, codon-aligned 51-nt CDS windows rewritten by synonymous recoding, which leaves the encoded amino-acid sequence and the ESM-2 input unchanged. The matched window was drawn from the bottom attention quintile of the same gene and region and matched on GC content and normalized position; each window was rewritten five times and scored in all five models. Vertical line and diamond, median; n = 161 (left) and 168 (right) held-out genes. (J) Architecture of the TransLocAttn model. See Methods for details. (K) Per-gene nucleotide-level Spearman correlation between the compartment-specific attention track and the eCLIP signal, for UTRs (left) and CDS (right). Genes are the 61 held-out genes whose top predicted compartment matched the observed one, selected using only observed and predicted compartment values. Every eligible gene-experiment pair was scored without selecting experiments by RNA-binding protein (median 8 experiments per gene, range 1 to 156; 1,171 pairs on the left and 1,177 on the right) and collapsed to one value per gene by the median. Tracks and eCLIP signal were Gaussian-smoothed (sigma = 5 nt) and rank-standardized within the 5’ UTR, 3’ UTR, or CDS before correlation. Vertical line and diamond, median; gray band, the 5th to 95th percentile of the same median under 199 circular shifts of the attention track within the same transcript regions. One-sided empirical P = 0.02 (left) and 0.01 (right); Benjamini-Hochberg-adjusted q = 0.02 in both, across the two region sets. (L-N) Examples of attention weights from TransLocAttn (top) and eCLIP-Seq read distributions (bottom) along the indicated mRNAs. eCLIP-Seq data with high correlation with the attention weights are shown with color-highlighted lines, and all other eCLIP-Seq datasets that do not pass the threshold are shown in gray. (O) Volcano plots showing eCLIP-Seq enrichment on CDSs for mRNAs locally translated on the OMM (defined in Figure 1F). eCLIP-Seq datasets with a *P* value of 0.01 or lower are shown. eCLIP-Seq data for LARP4 and AKAP1 are highlighted. (P) Box plots of changes in Ribo-Seq and RNA-Seq after siRNA-mediated knockdown of *FXR2* in the indicated mRNA groups. *P* values were calculated with the Mann Whitney *U* test (two-tailed). Effect sizes of RBP knockdown were estimated by the Hodges–Lehmann (HL) statistic (*FXR2*, HL = -0.059, 95% CI [-0.099, -0.018]). Box plots show the median (centerline), upper/lower quartiles (box limits), and 1.5× interquartile range (whiskers).

**Figure S7.**
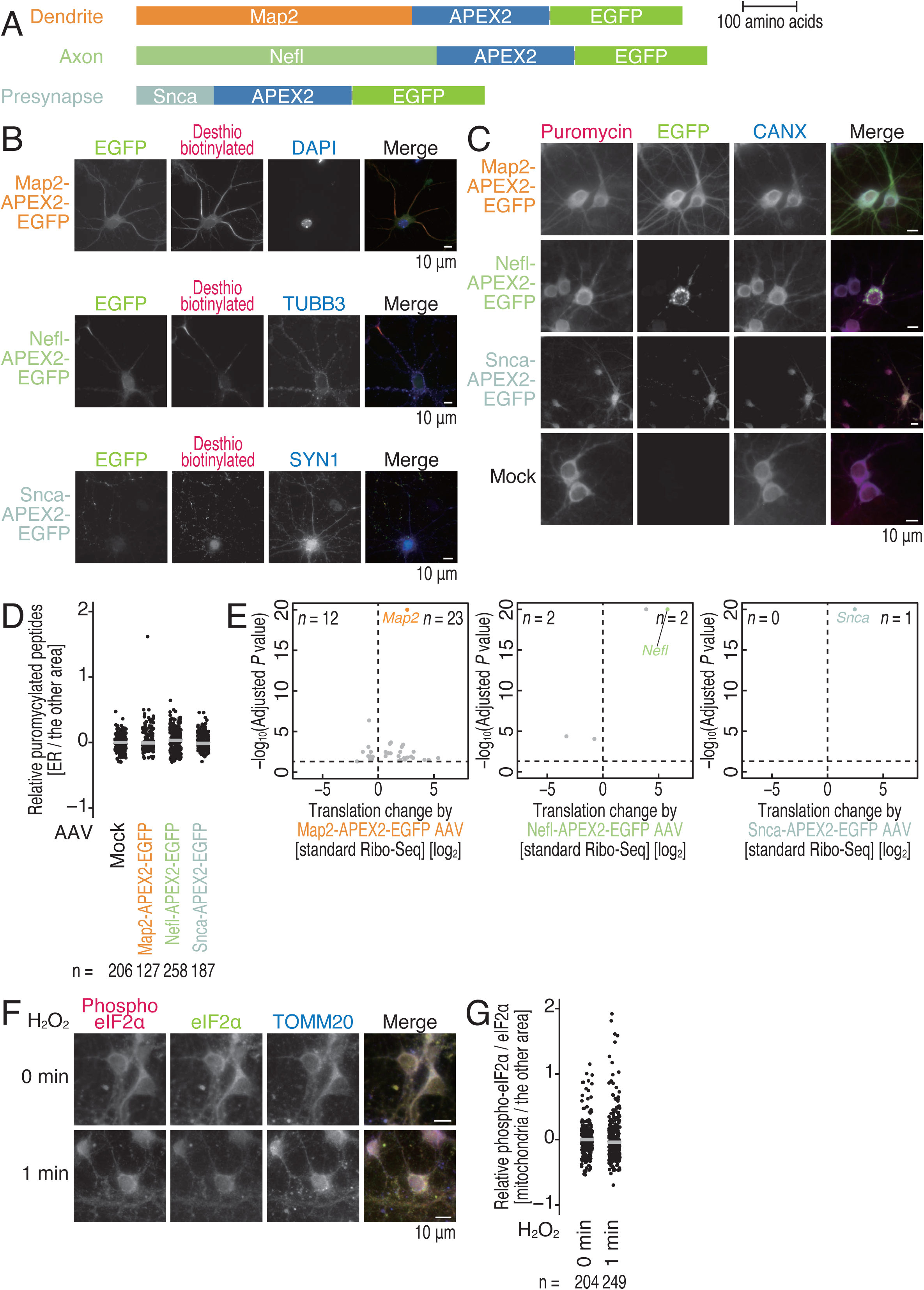
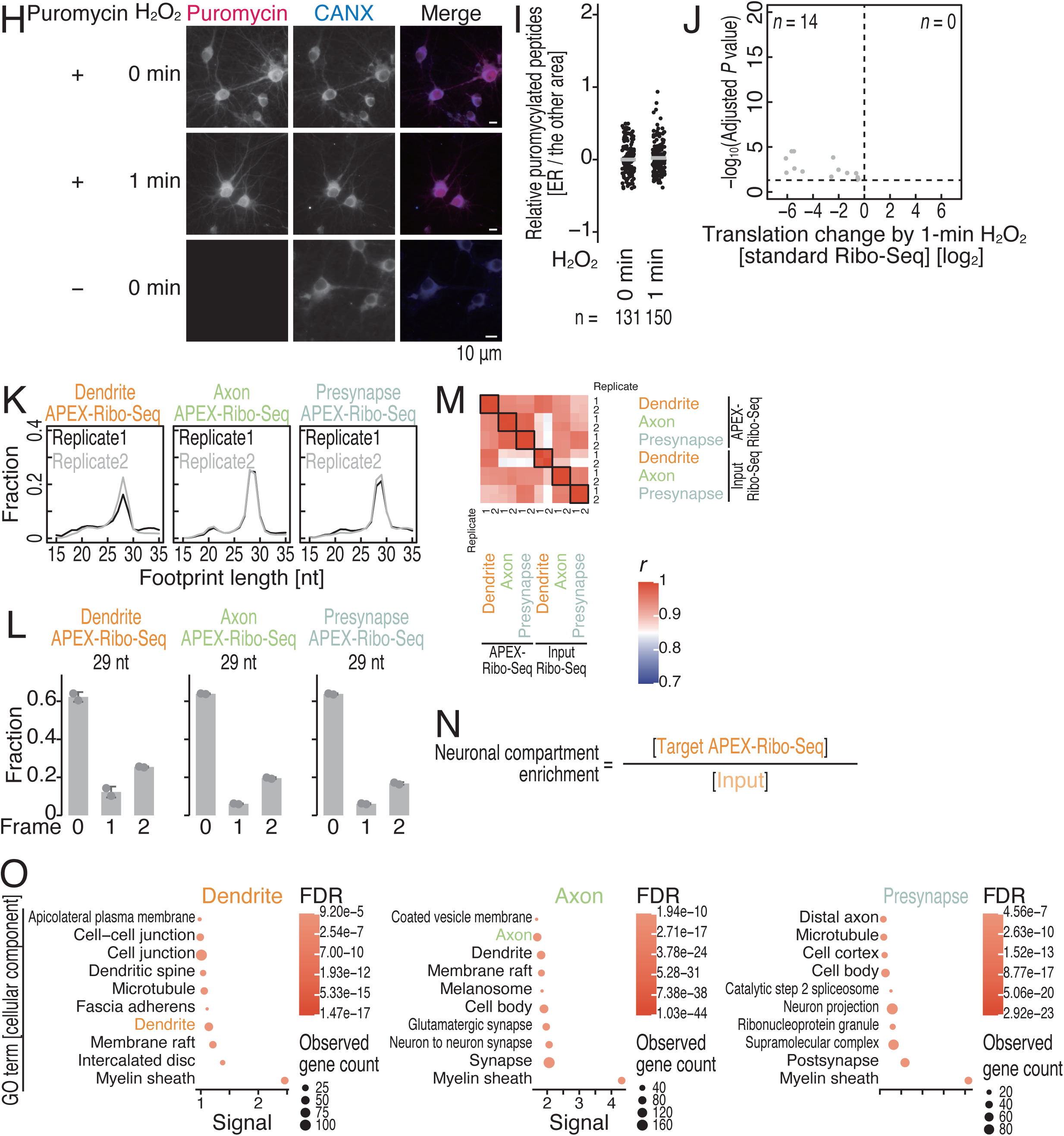
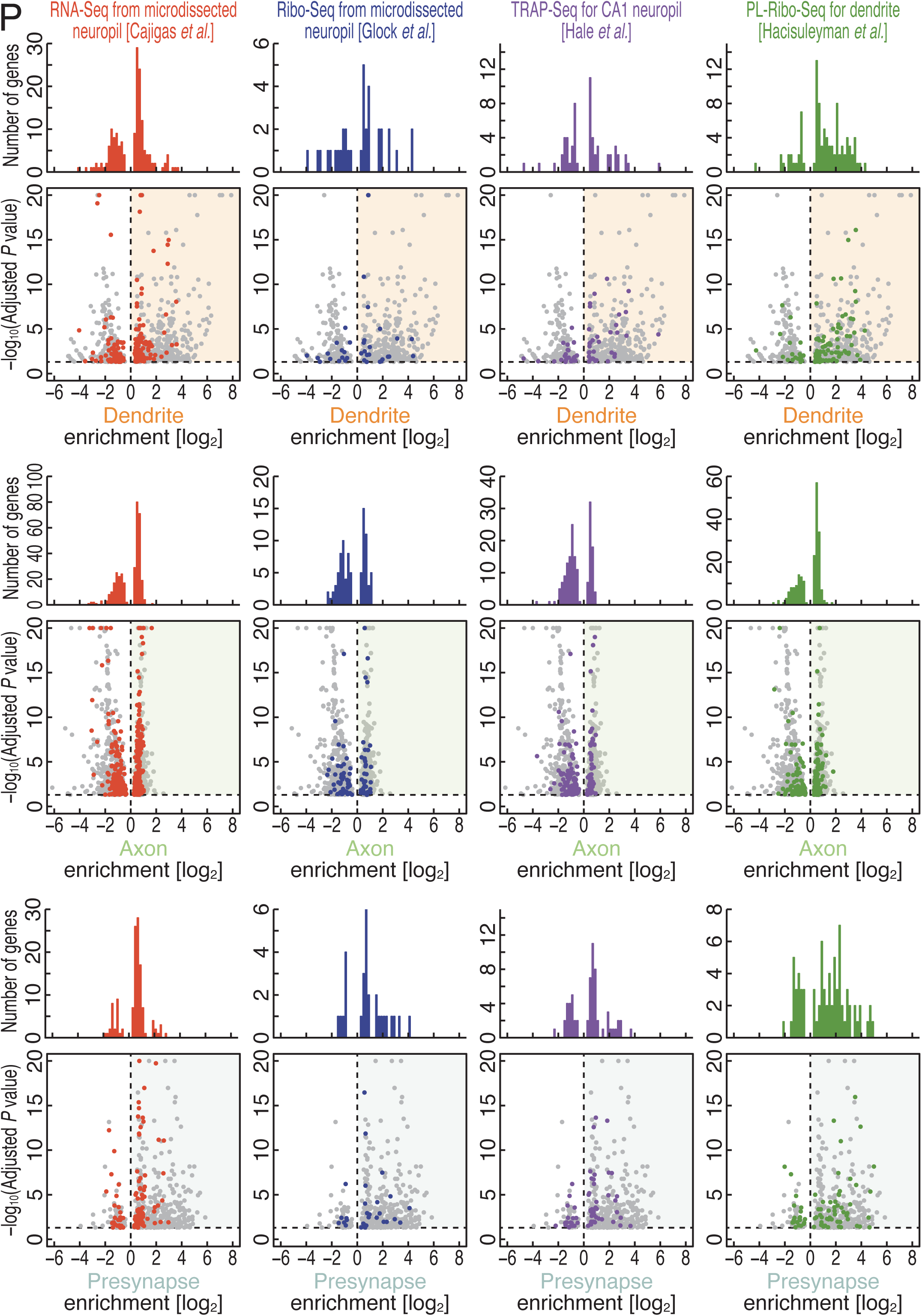
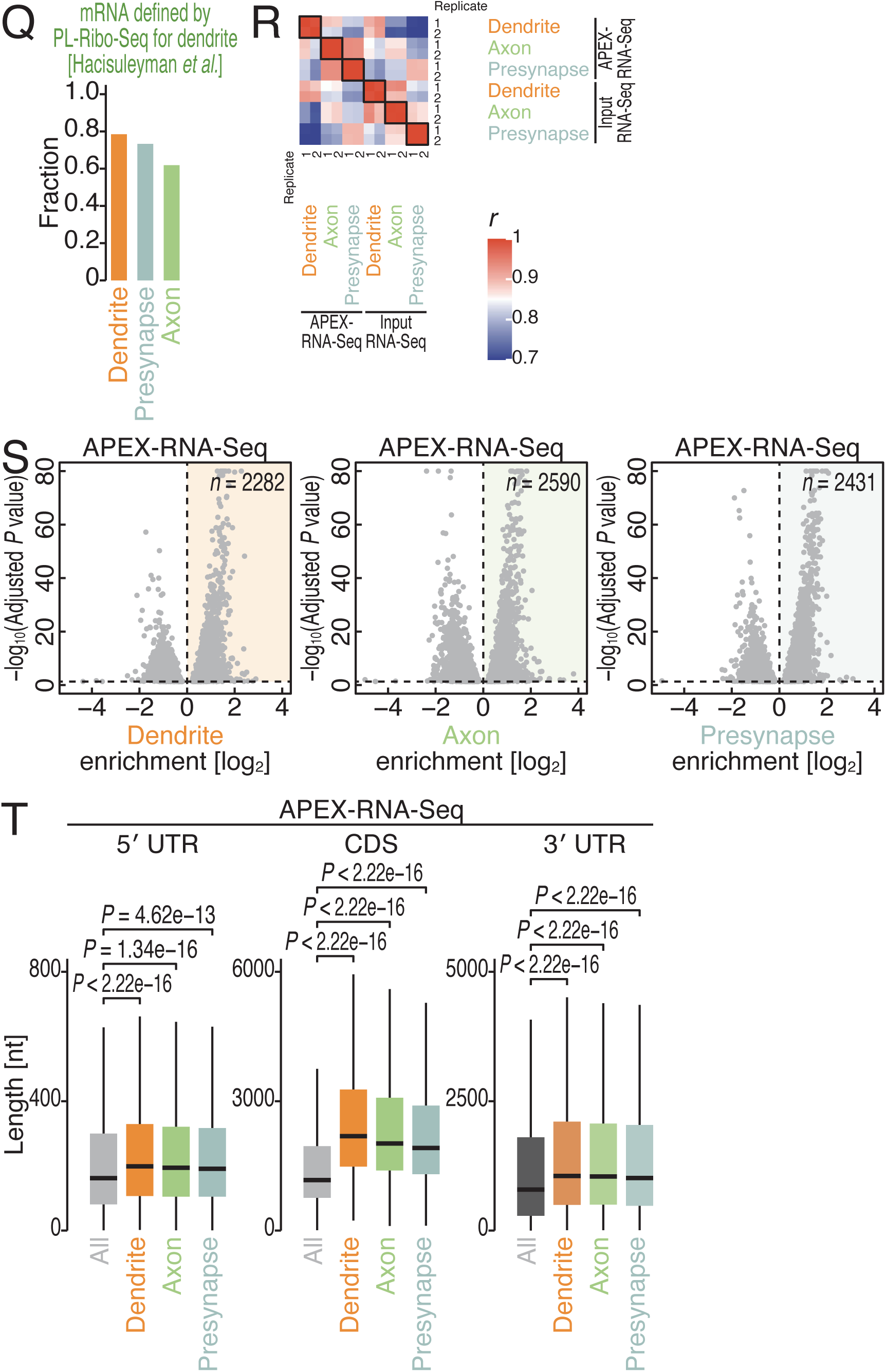
Characterization of local translation in neuronal compartments, related to Figure 6. (A) Schematics of constructs for APEX2 fusion protein expression. (B) Immunofluorescence analysis of the indicated proteins (detected by EGFP) with desthiobiotinylated proteins/RNAs (detected with Alexa Fluor 555-conjugated streptavidin) in mouse primary neurons. DAPI was used to stain the DNA. Tubb3 and Syn1 were detected as axonal and presynaptic markers, respectively. The scale bar represents 10 μm. (C and H) Immunofluorescence analysis of the indicated proteins (detected by EGFP) with the puromycylated peptides (detected with anti-puromycin antibody) in primary neurons. CANX was detected as a marker of the ER. The scale bar represents 10 μm. (D and I) Quantification of the relative level of puromycylated peptides. The normalized signal in the ER mask over the other cellular regions was shown. The median (bars) and individual data (points, n for the number of cells analyzed) are shown. (E) Volcano plots showing translation changes determined by standard Ribo-Seq in primary neurons. Relative changes induced by marker protein expression compared to the control AAV (APEX2-NES-GFP) are calculated. mRNAs with an adjusted *P* value of 0.05 or lower are shown. The marker genes expressed in each AAV are highlighted. (F) Immunofluorescence analysis of eIF2α and phosphorylated (phospho) eIF2α during H_2_O_2_ treatment in primary neurons. TOMM20 was detected as a marker of the mitochondria. The scale bar represents 10 μm. (G) Quantification of the relative level of phosphorylated (phospho) eIF2α over total eIF2α. The normalized signal in the mitochondria mask over the other cellular regions was shown. The median (bars) and individual data (points, n for the number of cells analyzed) are shown. (J) Volcano plots showing relative translation changes determined by standard Ribo-Seq in primary neurons after H_2_O_2_ treatment (at 1 mM for 1 min). mRNAs with an adjusted *P* value of 0.05 or lower are shown. (K) Distribution of ribosome footprint length under the indicated conditions. (L) Fraction of the frame position for the 5′ end of the footprint (29 nt) under the indicated conditions. The means (bars), s.d.s (errors), and individual replicates (n = 2, points) are shown. (M) Heatmap of the Pearson correlation coefficients (*r*) of APEX-Ribo-Seq and input standard Ribo-Seq under the indicated conditions. The value of *r* is indicated by the color scale. (N) Schematics of the neuronal compartment enrichment calculation. (O) GO analysis of cellular components ^93^ for the indicated mRNA groups. The results of the top 10 terms, ranked by signal score, are visualized. The color-highlighted terms are described in the main text. (P) Volcano plots same as in Figure 6A. On top, histograms for mRNAs defined by RNA-Seq (microdissected neuropil) ^154^, Ribo-Seq (microdissected neuropil) ^53^, TRAP-Seq (CA1 neuropil) ^155^, and PL-Ribo-Seq (dendrite) ^63^ within the volcano plots are shown. (Q) Fraction of mRNAs defined by dendrite PL-Ribo-Seq ^63^ among mRNAs translated in the indicated neuronal compartments (defined in Figure 6A). (R) Heatmap of the Pearson correlation coefficients (*r*) of APEX-RNA-Seq and input standard RNA-Seq under the indicated conditions. The value of *r* is indicated by the color scale. (S) Volcano plots showing neuronal compartment enrichment (Figure S7N) for dendrites, axons, and presynapses determined by APEX-RNA-Seq. mRNAs with an adjusted *P* value of 0.05 or lower are shown. The colored areas represent enriched mRNAs. (T) Box plots of the lengths of the 5′ UTR, CDS, and 3′ UTR of the indicated mRNA groups defined by APEX-RNA-Seq. *P* values were calculated with the Mann Whitney *U* test (two-tailed) and adjusted by the Benjamini–Hochberg method. Box plots show the median (centerline), upper/lower quartiles (box limits), and 1.5× interquartile range (whiskers).

**Figure S8.**
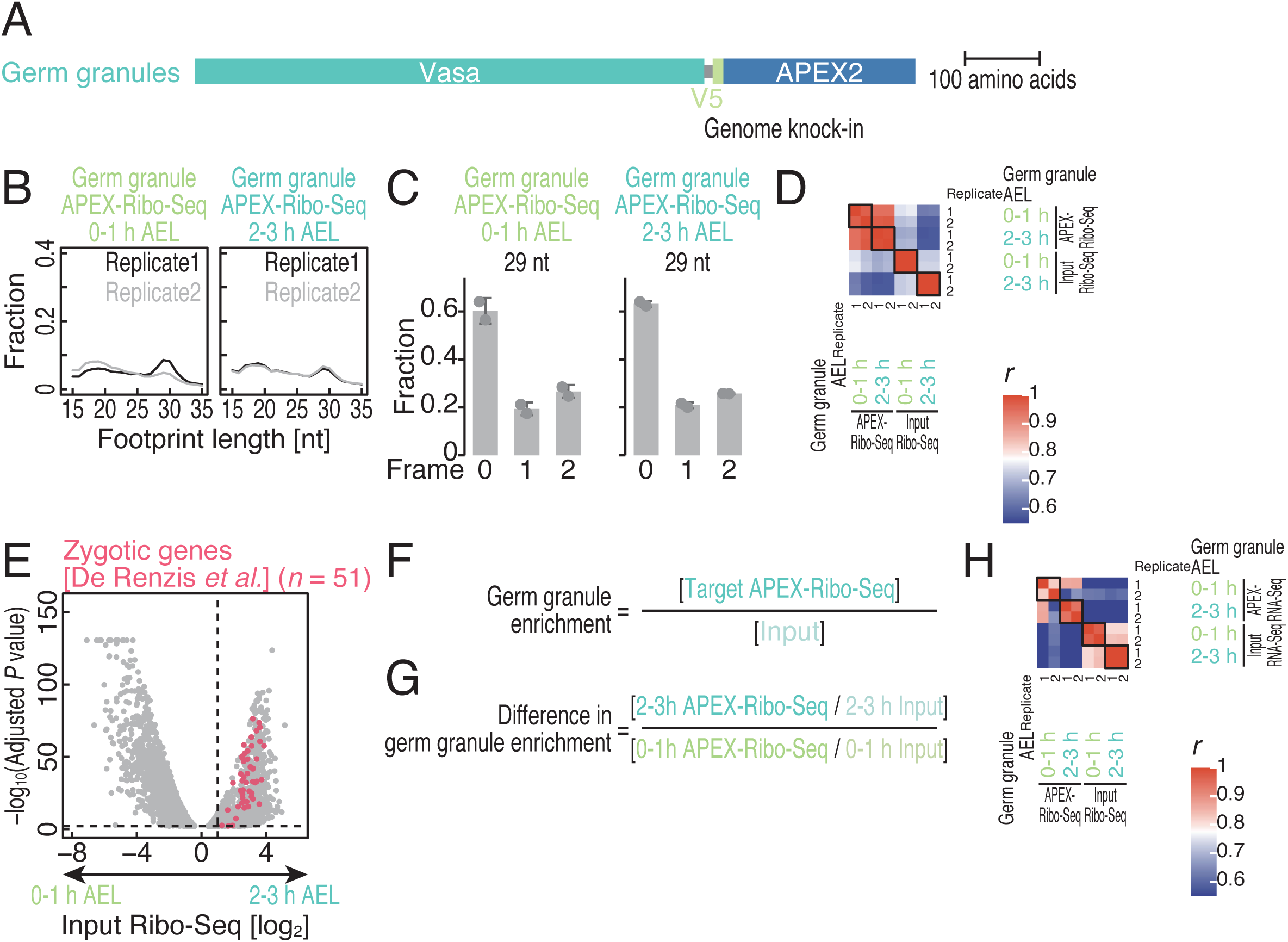
Characterization of local translation on fly germ granules, related to Figure 7. (A) Schematics of constructs for APEX2 fusion protein expression. (B) Distribution of ribosome footprint length under the indicated conditions. (C) Fraction of the frame position for the 5′ end of the footprint (29 nt) under the indicated conditions. The means (bars), s.d.s (errors), and individual replicates (n = 2, points) are shown. (D) Heatmap of the Pearson correlation coefficients (*r*) of APEX-Ribo-Seq and input standard Ribo-Seq under the indicated conditions. The value of *r* is indicated by the color scale. (E) Volcano plots showing translational differences for developmental stages determined by input standard Ribo-Seq. mRNAs with an adjusted *P* value of 0.05 or lower are shown. The zygotic genes defined in a previous study ^166^ are highlighted. (F) Schematics of the germ granule enrichment calculation. (G) Schematics of the calculation for the difference in germ granule enrichment in the developmental stages. (H) Heatmap of the Pearson correlation coefficients (*r*) of APEX-RNA-Seq and input standard RNA-Seq under the indicated conditions. The value of *r* is indicated by the color scale.

**Table S1. Log_2_ fold enrichment and statistical metrics of APEX-Ribo-Seq and APEX-RNA-Seq in HEK293 T-REx and U2OS cells, as well as MitoIP-Ribo-Seq, related to Figures 1-3**.

**Table S2. Summary of HOMER and TransLocAttn analysis, related to Figure 5**.

**Table S3. Log_2_ fold enrichment and statistical metrics of APEX-Ribo-Seq and APEX-RNA-Seq at neuronal compartments and germ granules, related to Figures 6-7**.

**Table S4. Library information of APEX-Ribo-Seq and APEX-RNA-Seq, related to Figures 1, 2, 4, 6, and 7.**

**Table S5. Sequence of smiFISH probes and organelle mask parameters, related to Figure 2**.

**Table S6. Statistics of the correlations shown in this work, related to Figures 4-5**.

**Table S7. List of antibodies used in this study, related to Figures 3 and 7**.

## Methods

### Experimental model and study participant details

#### Ethics statement

All animal procedures were approved by the Animal Experiment Committee of the National Institutes of Natural Sciences (Approval No. 24A024) and conducted in accordance with institutional guidelines and the recommendations of the Science Council of Japan.

#### Cell culture

HEK293 Flp-In T-REx cells (referred as HEK293 T-REx in the text; Thermo Fisher Scientific, R78007, female) with and without stable integration of APEX2-bait (Figures S1A) and U2OS cells (American Type Culture Collection [ATCC], HTB-96, female) were maintained in DMEM, high glucose, GlutaMAX Supplement (Thermo Fisher Scientific, 10566024) supplemented with 10% fetal bovine serum (FBS, Sigma Aldrich) at 37°C and 5% CO_2_.

#### Fly

The *Drosophila melanogaster vasa-V5-APEX2* fly line generated in this study was used for all experiments.

### Method details

#### DNA construction

##### pcDNA5/FRT/TO-APEX2 plasmids

The APEX2 gene with organelle-targeting signals or organelle markers was cloned and inserted into the pcDNA5/FRT/TO vector (Thermo Fisher Scientific, V652020) (Figure S1A). PCR amplification and assembly of the DNA fragments were performed with PrimeSTAR MAX (TaKaRa, R045A) and In-Fusion HD (TaKaRa, 639649), respectively. For the constructs designed to target the cytosol (NES), mitochondrial matrix, endoplasmic reticulum membrane (ERM), and outer mitochondrial membrane (OMM), we followed the procedures used in previous APEX-Seq studies ^64,65,67^. For peroxisomes, *cis*-Golgi, *trans*-Golgi, lysosomes, endosomes, early endosomes, cell junctions, intermediate filaments, and clathrin-coated vesicles, we selected marker proteins on the basis of proximity proteomics data ^100^. We used the transmembrane domain of GAP43 for the plasma membrane ^197,198^ and SYNE3 for the outer nuclear membrane ^199^.

##### pSEPT-CEP192-APEX plasmid

The pSEPT-CEP192-APEX2 construct was generated from the pSEPT-CEP192-NeonGreen plasmid ^200^, replacing NeonGreen with APEX2 and a V5 tag.

##### pAAV[Exp]-CBh>bait-APEX-GFP:WPRE3 plasmids

Neuronal compartment markers tagged with APEX2-GFP at the C-terminus were cloned and inserted into a single-stranded adeno-associated virus (ssAAV) mammalian gene expression vector containing a CBh promoter and a WPRE3 regulatory element (Figure S7A). We used mouse Map2 for dendrites ^149^, mouse Nefl for axons ^150^, and mouse Snca for presynapses ^151^. APEX2-NES-GFP was used only to estimate the effect of overexpression of each marker gene as a control. All pAAV[Exp] vectors were constructed by Vector Builder.

##### pBS-vasa 5′ region-V5-APEX2-loxP-dsRed-loxP-vasa 3′ region-plasmid

The *vasa* 5′ and 3′ homology arms were amplified from *yw* genomic DNA and cloned into pBlueScript ^201^ together with the V5-APEX2 coding sequence, preceded by a GGSGSGSS linker and followed by a stop codon, and a loxP-flanked 3×P3-dsRed marker, yielding pBS-5′ region-V5-APEX2-loxP-dsRed-loxP-3′ region.

#### Generation of cell lines

The pcDNA5/FRT/TO vectors were stably integrated into the HEK293 T-REx genome via cotransfection with pOG44 (Thermo Fisher Scientific, V600520) using X-tremeGENE9 (Roche, 6365809001) and selected with blasticidin S (InvivoGen, ant-bl-1) and hygromycin B (InvivoGen, ant-hg-1). In the subsequent experiments, cells were incubated with tetracycline for 3 d to induce APEX2 protein expression.

For the CEP192-APEX2-V5 cell line, we performed a C terminal knock in to the endogenous locus using the CRISPR-Cas9 system as previously described ^200^, with modifications. The vector expressing Cas9 and guide RNA targeting the C-terminal region of the *CEP192* gene was cotransfected with the repair construct pSEPT-CEP192-APEX using Lipofectamine 3000 (Thermo Fisher Scientific, L3000015). After incubation for 14 d with G418 (InvivoGen, ant-gn-1), integrants were seeded in 96-well plates at 0.1 cells per well to isolate single-cell clones. Integration was confirmed by immunostaining.

For U2OS cell lines, the pcDNA5/FRT/TO vectors were transiently expressed via transfection with Lipofectamine LTX (Thermo Fisher Scientific, 15338100). Cells were subjected to APEX-Ribo-Seq 48 h after transfection.

#### CRISPR/Cas9-mediated genome editing of the endogenous vasa locus

The genome-edited fly line was obtained using CRISPR/Cas9-mediated homology-directed repair. Guide RNA (gRNA) was designed to target the PAM sequence just upstream of the stop codon of *vasa*. pCFD3 gRNA expression plasmid, pBS-vasa 5′ region-V5-APEX2-loxP-dsRed-loxP-vasa 3′ region donor plasmid, and pBS-Hsp70-Cas9 plasmid (a gift from Melissa Harrison, Kate O’Connor-Giles, and Jill Wildonger, Addgene plasmid # 46294; http://n2t.net/addgene:46294; RRID: Addgene_46294) ^202^ were coinjected using the standard *yw* laboratory strain. Microinjection was performed as previously described ^203^. *3×P3-dsRed* was used for screening. Resulting genome-edited flies were then crossed with y1 w67c23 P{Crey}1b; snaSco/CyO (Bloomington Drosophila Stock Center, #766) to remove the *3×P3-dsRed* marker from the endogenous locus. Insertion was confirmed by PCR analysis of purified genomic DNA.

#### Drug treatment

CEP192-APEX2-V5 cells were treated with 500 nM centrinone-B (Tocris Bioscience, 5690) for 4 d to deplete centrosomes.

#### Primary neuron culture and AAV production

The AAV vectors were packaged as described previously ^204^ using the AAV-DJ Helper Free Expression System (Cell Biolabs, VPK-410-DJ). Briefly, the pAAV transfer plasmid was transfected into HEK293T cells with packaging plasmids. The AAV vector particles in the crude lysates were purified by serial ultracentrifugation with cesium chloride and dialyzed against PBS containing 0.001% Pluronic F-68 (Sigma Aldrich, P5556-100ML), followed by concentration with an Amicon 10 K MWCO filter (Merck Millipore, UFC801096). The viral genome (vg) copy number was measured by real-time quantitative PCR with primers targeting the WPRE3 region (forward 5′-GGTATTCTTAACTATGTTGCTCCTT-3′ and reverse 5′-GAATTGTCAGTGCCCAACAG-3′).

Primary cortical neurons were prepared as previously described ^205^. Briefly, cortices were isolated from embryonic day 15.5 (E15.5) ICR mouse embryos, enzymatically and mechanically dissociated, and plated onto poly-D-lysine-coated 10-cm culture dishes at 2.0 × 10^6^ cells/ml. The neurons were maintained in a humidified 5% CO_2_ incubator at 37°C in Neurobasal-A medium (Thermo Fisher Scientific, 10888-022) supplemented with 2% B-27 (Thermo Fisher Scientific), 1% GlutaMAX (Thermo Fisher Scientific, 35050-061), and 25% Neuron Culture Medium (FUJIFILM Wako, 148-09671). At 6 days *in vitro* (DIV6), AAV vectors were added directly to the culture medium at a final concentration of 1.4 × 10^11^ vg/ml. Neurons at DIV9 were used for downstream experiments.

#### In situ APEX labeling in human cell culture and mouse primary neurons

APEX labeling was initiated by replacing the culture medium with fresh medium containing 0.5 mM desthiobiotin tyramide (Iris Biotech, LS-1660). For cultured primary neurons, 0.5 mM desthiobiotin tyramide was added directly to the medium. After a 30-min incubation, the reaction was started by treating the cells with a final concentration of 1 mM H_2_O_2_ (nacalai tesque, 18411-25) for 1 min at room temperature. The reaction was quenched by replacing the medium with quench PBS buffer consisting of 10 mM sodium azide (nacalai tesque, 31233-42), 10 mM sodium ascorbate (Sigma Aldrich, A7631), and 5 mM Trolox (Sigma Aldrich, 238813) in PBS; the cells were subsequently washed twice with quench PBS buffer. After the second wash step, the cells were immediately placed on ice.

Quenched cells were lysed directly on dishes with 500 μl of APEX lysis buffer (20 mM Tris-HCl pH 7.5, 150 mM NaCl, 5 mM MgCl_2_, 1% Triton X-100, 1 mM dithiothreitol, 100 µg/ml cycloheximide, 100 µg/ml chloramphenicol, 10 mM sodium azide, 10 mM sodium ascorbate, and 5 mM Trolox) and incubated with 25 U/ml TURBO DNase (Thermo Fisher Scientific, AM2238) for 10 min. After DNase treatment, the lysates were clarified by centrifugation at 20,000 × *g* for 10 min at 4°C.

#### In situ APEX labeling in fly embryos

APEX labeling in fly embryos was performed with slight modifications to a protocol previously described for fly ovaries ^164^. Flies were provided with yeast paste and allowed to lay eggs on apple-juice agar plates. After a 2 h pre-lay, embryos were collected for 1 h to obtain 0-1 h after egg laying (AEL) samples. For 2-3 h AEL samples, embryos were collected for 1 h and aged for 2 h at 25 °C. Embryos were dechorionated in bleach for 2 min and washed extensively with Milli-Q water. Dechorionated embryos were immersed in 0.16% digitonin and 1 mM desthiobiotin tyramide in 5 ml of PBS, and the embryos were incubated at room temperature for 10 min. Freshly diluted H_2_O_2_ was added to the solution containing embryos to a final concentration of 1 mM and incubated for 1 min with shaking. Labeled embryos were immediately flash-frozen in liquid nitrogen and stored at −80 °C. Frozen embryos were mixed with 300 µl of frozen APEX lysis buffer droplets, and then ground at 3000 rpm for 15 s in a Multi-beads Shocker (YASUI KIKAI). The lysate was slowly thawed at 4°C and treated with 10 U Turbo DNase (Thermo Fisher Scientific) on ice for 10 min to remove genomic DNA. The lysate was then centrifuged at 20,000 × *g* for 30 min at 4°C, and the supernatant was collected.

#### Purification of desthiobiotinylated ribosomes and RNAs

To remove unreacted desthiobiotin tyramide, the supernatant was applied to a NAP5 column (Cytiva, 17085302) according to the manufacturer’s instructions. Elution was performed with 1000 μl of APEX lysis buffer, and 150 μl of the lysate was retained to serve as the input samples for APEX-Ribo-Seq and APEX-RNA-Seq. The remaining lysate was subjected to further purification.

To enrich the sample for biotinylated proteins and RNA, 100 μl of Dynabeads MyOne Streptavidin C1 (Thermo Fisher Scientific, 65002) pre-equilibrated with APEX lysis buffer was added to the NAP5 eluate and incubated at 4°C for 1 h. The beads were washed five times with 500 µl of APEX lysis buffer, with transfer to a fresh tube during each wash step. For elution, the beads were resuspended in 150 μl of APEX lysis buffer supplemented with 5 mM D-biotin (Thermo Fisher Scientific, B20656) and incubated overnight at 4°C. The supernatant was collected, and the beads were preincubated with an additional 150 μl of APEX lysis buffer containing 5 mM D-biotin for 1 h at 4°C. This elution step was repeated twice, yielding 450 μl of eluate in total.

#### Library preparation for Ribo-Seq

For the streptavidin pulldown and corresponding input samples, Thor Ribo Seq library preparation was performed with 300 μl of eluate and 100 µl of lysate, respectively ^72^. Briefly, lysates were treated with 0.067 U/µl RNase I (LGC Biosearch Technologies, N6901K) at 25°C for 45 min. After ultracentrifugation, RNA was extracted from the ribosome pellet using TRIzol reagent (Thermo Fisher Scientific, 15596018) and a Direct-zol RNA MicroPrep Kit (Zymo Research, R2062). The RNA fragments corresponding to 17–34 nt in length were excised from the gel, dephosphorylated by T4 polynucleotide kinase (New England Biolabs, M0201S), and ligated to the 3′ DNA linkers containing barcodes and the T7 promoter sequence by T4 RNA Ligase 2, truncated KQ (New England Biolabs, M0373L). After hybridization of complementary DNA to the T7 promoter sequence, RNAs were amplified by *in vitro* transcription using the T7-Scribe Standard RNA IVT Kit (CELLSCRIPT, C-AS3107). rRNA depletion was performed using Human Ribo-Seq riboPOOL (siTOOLs biotech, dp-K024-000042) and Mouse-Rat Ribo-Seq riboPOOL (siTOOLs biotech, dp-K024-000052). The RNAs were then ligated with a 5′ linker, reverse transcribed with ProtoScript II Reverse Transcriptase (New England Biolabs, M0368L), and amplified by PCR.

For the centrinone B-treated samples, cell lysates containing 10 μg of total RNA were used for the Thor Ribo Seq library preparation. The RNA concentration of the lysates was measured with a Qubit RNA BR kit (Thermo Fisher Scientific, Q10210). The mitochondria immunoprecipitation (MitoIP) Ribo-Seq was performed in

HEK293 T-REx cells as previously described ^89^.

#### Library preparation for RNA-Seq

For the streptavidin pulldown and corresponding input samples, total RNA was extracted from 150 μl of eluate and 50 µl of lysate, respectively, with TRIzol LS (Thermo Fisher Scientific, 10296-010) and a Direct-zol RNA MicroPrep Kit (Zymo Research, R2062). The libraries were prepared using the SEQuoia Express Stranded RNA Library Prep Kit (Bio-Rad, 12017265). For the centrinone B-treated samples, 500 ng of total RNA extracted from the cell lysate was used.

All libraries were sequenced on a NovaSeq 6000, NovaSeq X Plus, or HiSeq 4000 system (Illumina) to produce 150-nt paired end reads.

#### SDS-PAGE, SYPRO Ruby protein staining, and Western blotting

Equal amounts of purified lysates from APEX2 NES EGFP cells under the indicated conditions were separated by SDS PAGE. The gels were stained with SYPRO Ruby Protein Gel Stain (Thermo Fisher Scientific, S12001) and imaged using a Pharos FX imaging system (Bio-Rad).

For Western blotting, the proteins were transferred from the gels to nitrocellulose membranes, which were subsequently blocked with Intercept (TBS) Blocking Buffer (LI-COR, 927-60001). Immunoblotting was performed using an anti-RPS17 (Abcam, AB138991, 1:1,000) primary antibody and an IRDye 680RD anti-rabbit IgG (LI-COR, 926-68071, 1:10,000) secondary antibody. IRDye 800CW streptavidin (LI-COR, 925-32230, 1:2,000) was used for the detection of desthiobiotinylated proteins. Images were acquired with an Odyssey CLx imager (LI-COR).

#### Polysome profiling

Polysome profiling was performed as previously described ^206,207^, with some modifications. Lysates with in situ APEX labeling (as described in the “*In situ APEX labeling in human cell culture and mouse primary neurons*” section) were loaded onto the top of a 10–50% (w/v) sucrose gradient in APEX polysome buffer (20 mM Tris-HCl pH 7.5, 150 mM NaCl, 5 mM MgCl_2_, 1 mM dithiothreitol, 100 µg/ml cycloheximide; and 10 mM sodium azide) and centrifuged at 221,230 × *g* for 2.5 h at 4°C in a P40ST rotor (Hitachi Koki). Fractionation and absorbance monitoring were performed using the Gradient Station (BioComp) and the TRiAX Flowcell (BioComp). For Western blotting, the fractions were incubated with 20% trichloroacetic acid (Fujifilm Wako) for 20 min on ice and then centrifuged at 20,000 × *g* for 15 min at 4°C. The precipitates were washed twice with 150 µl of ice-cold acetone, dried, and resuspended in 15 µl of 1 M Tris-HCl, pH 7.5. For RNA dot blotting, RNA was extracted from the fractions using a Maxwell RSC simplyRNA Cells Kit (Promega, AS1390) and a Maxwell RSC Instrument (Promega).

#### RNA dot blotting

The RNA samples were diluted in RNA denaturing buffer (50% dimethyl sulfoxide and 1 M glyoxal in 0.01 M Na_2_PO_4_-NaHPO_4_ buffer, pH 7.5) and incubated at 50°C for 1 h. The mixtures were spotted on a Zeta-Probe blotting membrane (Bio-Rad, 1620153) with Bio-Dot SF (Bio-Rad, 1706542), then washed once with 100 µl of TE buffer (10 mM Tris-HCl and 1 mM ethylenediaminetetraacetic acid, pH 8.0) and once with 100 µl of 2×SSC (0.3 M NaCl and 0.03 M sodium citrate, pH 7.0). The membranes were then dried, incubated at 80°C for 1 h for crosslinking, blocked for 2 h using Intercept (TBS)

Blocking Buffer with 1% SDS, and blotted with IRDye 800CW Streptavidin (LI-COR, 925-32230, 1:2,000). Total RNA was visualized using methylene blue staining buffer (0.2% methylene blue in 0.4 M sodium acetate and 0.4 M acetic acid). Images were acquired using an Odyssey CLx (LI-COR).

#### Immunofluorescence staining and fluorescence microscopy

Cells seeded on 12-mm coverslips (Matsunami glass, C012001) were fixed in 4% paraformaldehyde in PBS at room temperature for 15 min. After two washes with PBS, the cells were permeabilized in 0.1% Triton X 100 in PBS at room temperature for 10 min, again washed twice with PBS, and blocked for 1 h at room temperature in Intercept (TBS) Blocking Buffer (LI-COR, 927-60001) containing 0.2% Triton X 100. The cells were then incubated overnight at 4°C with the primary and secondary antibodies listed in Table S7. The cells were washed twice with PBS containing 0.2% Triton X 100 and mounted in VECTASHIELD Vibrance (Vector Labs, H-1800). Images were acquired using a Zeiss LSM 710 confocal microscope with a 63× objective lens (Zeiss, Plan-Apochromat 63x/1.4 Oil DIC M27) or an Olympus IX83 inverted microscope with a 100× objective lens (Olympus, UPLXAPO100XO) and a CMOS camera ORCA-Fusion (Hamamatsu Photonics, C15939-20U).

Labeled embryos were fixed at room temperature for 45 min in a biphasic mixture of 5 ml of fixation buffer (1× PBS and 7.4% formaldehyde in nuclease-free H_2_O) and 5 ml of heptane. The aqueous layer was removed, 10 ml of methanol was added, and the embryos were shaken for 1 min for devitellinization. The devitellinized embryos were washed three times with methanol and rehydrated in PBST (1× PBS without CaCl_2_ and MgCl_2_, and 0.05% Tween 20). The embryos were blocked for 30 min at room temperature in blocking solution (1.5× Western Blocking Reagent [Roche, 11921673001] in PBST), incubated overnight at 4°C with the primary antibody listed in Table S7, washed with PBST, blocked again for 30 min in blocking solution, and then incubated for 2 h at room temperature with the fluorescent secondary antibody listed in Table S7. The embryos were washed with PBST, stained with DAPI for 10 min, washed again with PBST, and mounted in ProLong Gold Antifade Mountant (Thermo Fisher Scientific, P36930).

#### smiFISH

Single-molecule inexpensive FISH (smiFISH) was performed as previously described ^94,208^ with modifications. Cells seeded on 12-mm coverslips were fixed in 4% paraformaldehyde in PBS at room temperature for 15 min, washed three times with PBS, and then permeabilized in 70% ethanol in PBS overnight at 4°C. After two washes with PBS, the coverslips were incubated with 120 µl of prehybridization buffer (2×SSC containing 10% formamide, 5 mg/ml UltraPure BSA [Thermo Fisher Scientific, AM2618], 2 mM vanadyl ribonucleoside complexes solution [Sigma Aldrich, 94742], and 0.1% Triton X 100) for 30 min at room temperature and subsequently hybridized with FLAP probes.

To prepare the FLAP probes, target specific oligonucleotide probe sets (0.4 µM each, listed in Table S5) were mixed with 20 µM Cy3- or Cy5-labeled FLAP oligo (synthesized by Integrated DNA Technologies) in FLAP buffer (5 mM Tris-HCl pH 7.5, 10 mM NaCl, and 1 mM MgCl_2_). The mixture was incubated at 85°C for 3 min, 65°C for 3 min, and 25°C for 5 min using a thermal cycler for annealing, and then held at 12°C until use.

For hybridization, cells on coverslips were placed on a drop of 20 µl hybridization mix comprising FLAP probe duplexes (4 nM each), 10% formamide, 0.5 mg/ml BSA, 2 mM vanadyl ribonucleoside complex, 0.1% Triton X 100, 1 mg/ml yeast tRNA (Thermo Fisher Scientific, AM7119), and 12% dextran sulfate and incubated at 37°C for 3 h. Coverslips were then washed three times with RNA wash buffer (2×SSC containing 10% formamide and 0.1% Triton X 100) and incubated in RNA wash buffer for 30 min at 37°C, followed by immunostaining and mounting as described in the “*Immunofluorescence staining and fluorescence microscopy*” section. Images were obtained with an Olympus IX83 inverted microscope with a 100× objective lens (Olympus, UPLXAPO100XO) and a CMOS camera ORCA-Fusion (Hamamatsu Photonics, C15939-20U).

#### Puro-PLA

Puromycylation with the proximity ligation assay (Puro-PLA) was performed as previously described ^131^. Briefly, cells were incubated with 2 μM puromycin for 10 min, then fixed in 4% paraformaldehyde in PBS at room temperature for 15 min. After three washes with PBS, cells were permeabilized in 70% ethanol in PBS overnight at 4°C. Proximity ligation reaction was performed with Duolink reagents (Sigma Aldrich, DUO92002 and DUO92004) and Duolink Detection reagents (Sigma Aldrich, DUO92008) according to the manufacturer’s instructions. Antibodies used in this study are listed in Table S7.

#### Puromycylation assay

The puromycylation assay was performed as previously described ^209^. Briefly, primary neurons were incubated with 10 μM puromycin for 5 min, then fixed in 4% paraformaldehyde in PBS at room temperature for 15 min. After three washes with PBS, cells were permeabilized in 0.1% Triton X 100 in PBS at room temperature for 10 min, again washed twice with PBS, and blocked for 1 h at room temperature in Intercept (TBS) Blocking Buffer (LI-COR, 927-60001) containing 0.2% Triton X 100. The cells were then incubated overnight at 4°C with the primary and secondary antibodies listed in Table S7.

### Quantification and statistical analysis

#### smiFISH and Puro-PLA spot detection and quantification

smiFISH and Puro-PLA images were processed as maximum-intensity z-projections and then analyzed using a custom pipeline in Fiji (ImageJ, ver. 2.16.0) ^210^ and then analyzed using a custom pipeline. Nuclei were segmented via CellPose (ver. 2.2, nuclei model) ^211^, and cell boundaries were estimated by watershed expansion of nuclear masks within cytoplasmic regions defined by the smiFISH and Puro-PLA fluorescence signal. Organelle masks were generated for each image by applying an organelle-specific Laplacian of Gaussian filter, followed by adaptive thresholding. To refine each mask, small objects and holes were removed by size-based morphological filtering using the “skimage.morphology” module. The detailed filtering and thresholding parameters for each organelle are provided in Table S5.

smiFISH and Puro-PLA puncta were detected with Big FISH (ver. 0.6.2) ^212^ using parameters optimized for a 100 nm pixel size and ∼150 nm spot radius. The detected spots were assigned to individual cells on the basis of their spatial overlap with the segmentation masks. Colocalization with organelles was defined by whether the center of a spot fell within the refined organelle masks in the corresponding cell. The number and fraction of colocalized spots were quantified on a per-cell basis.

#### Quantification of immunofluorescence staining

Per-cell puromycin signal or phosphorylated eIF2α / eIF2α levels were quantified from immunofluorescence images using single-cell segmentation masks generated as above. For each labeled cell, mean pixel intensities were extracted from the puromycin, the phosphorylated eIF2α, and total eIF2α channels to calculate their ratio. For compartment-associated quantification, organelle masks were generated as described above and intersected with each single-cell mask. For each cell, mean intensities of each channel were measured separately within the organelle mask and within the complementary untargeted region. All image acquisitions and quantifications were performed using identical microscope settings across conditions.

#### Mapping of deep sequencing data

The libraries generated in this study are summarized in Table S4. The sequencing reads were processed as previously described ^72,213^. Briefly, we used fastp (ver. 0.21.0) ^214^ for read quality filtering and adapter trimming, cutadapt (ver. 3.7) ^215^ for demultiplexing by linker barcodes, and UMI-tools (ver. 1.1.2) ^216^ for deduplication on the basis of unique molecule identifiers. Reads that mapped to noncoding RNAs were removed, and the remaining reads were aligned to the human genome hg38, mouse genome mm39, or *Drosophila* genome dm6 and assigned to the GENCODE Human release 44 reference or Mouse VM36 reference or the UCSC dm6 Ensembl Genes annotation using STAR

(ver. 2.7.0a) ^217^. Libraries of poor quality (see Table S4 for details) were used only to show the correlation between replicates (Figure S3E and S6B) and were excluded from other downstream analyses. All downstream analyses focused exclusively on canonical transcripts defined by MANE Select (ver. 1.0) for humans ^218^. Ensembl canonical genes that were missed by MANE Select were also included in the list. For the mouse datasets, the Ensembl canonical gene set was used. For the *Drosophila* datasets, the FlyBase transcript annotation was used. Because neither MANE Select nor an Ensembl canonical set is defined for *Drosophila*, transcript-level quantification was performed over all annotated isoforms, and a single representative isoform per gene was selected *post hoc* as the isoform with the smallest adjusted *P* value in the DESeq2 analysis.

#### Enrichment calculation for 14 organelle compartments

Transcripts with ≥10 read counts in both NES Pulldown replicates (7408 genes in total) were included for downstream APEX Ribo Seq and APEX RNA Seq analyses of the 14 organelles. Mitochondrial genome encoded genes were excluded, except for the matrix APEX Ribo/RNA Seq datasets and the associated analysis. Genes used as APEX baits were also excluded.

Differential expression analysis of the read counts was performed using DESeq2 (ver. 1.32.0) ^219^. The log_2_-transformed fold enrichment for each region was calculated by comparing the count for each pulldown to that of the cytosolic control, with normalization to the input samples, *i.e.*, log_2_[(target pulldown/input)/(cytosol pulldown/input)] (Figure S2E). The input libraries for all 14 regions were handled as replicates, as previously described ^64^.

The specificity and sensitivity of ERM APEX-Ribo-Seq were evaluated via receiver operating characteristic (ROC) curve analysis ^92^. Using BirA proximity specific ribosome profiling as a positive reference ^58^, positive mRNAs for local translation were defined by log_2_ fold enrichment (ERM/cytosol) > 0.503 and adjusted *P* value < 0.05 (Figure S2F). These criteria were applied to all 14 organelle datasets of both APEX-Ribo-Seq datasets. The log_2_ fold enrichment for all analyzed mRNAs across each subcellular location is summarized in Table S1.

#### Protein annotations and localization database

Annotation of secretory and membrane-inserted/translocated genes was performed using Phobius ^220^, TMHMM ^221^, and SignalP ^222^, as described in earlier studies ^64,66^. MitoCarta 3.0 ^97^ was used for the annotation of mitochondrial proteins. Protein localization data were obtained from three sources: native organelle IP ^114^, immunofluorescence-based annotations in the Human Protein Atlas (HPA) ^100,115^, and a proximity-dependent biotinylation dataset generated via BioID ^100,115^. For the OpenCell dataset, we used 15 localization categories defined by the image-based classifier annotation, excluding the “unclassified” class. For the HPA dataset, we used 28 major protein localization classes defined by immunofluorescence imaging. For the BioID dataset, we used 19 nonnegative matrix factorization (NMF)-derived localization categories, excluding the “miscellaneous” class. For each dataset, genes identified as positive for a given compartment by APEX-Ribo-Seq were extracted, and the fraction of these genes falling into each localization category was calculated. The resulting fractions were *Z*-scored across the 14 APEX-Ribo-Seq compartments for each category and visualized as a hierarchically clustered heatmap.

#### Functional term clustering by STRING

Functional term clustering was performed using the STRING database (ver. 12.0) ^93^. Terms with a high-confidence interaction score cutoff (combined score ≥ 0.7) and a false discovery rate (FDR) < 0.05 were retained for subsequent analyses. Clusters with term similarity > 0.5 were merged. To highlight compartment-specific biological processes, clusters that appeared in more than four compartments were excluded.

#### Detection of co-co–assembled protein complexes

Locally assembled heteromeric protein complexes were identified using a predicted co-co–translation dataset ^129^. Complexes were classified as uniquely co-co–translated if at least one constituent gene was detected exclusively in a single compartment within the local translation atlas.

#### Enrichment analysis for short sequence motifs

To identify sequence motifs associated with differential translation localization across the 14 compartments, we employed HOMER (ver. 5.1) ^223^ for *de novo* motif discovery from the full-length transcript sequences in the mRNAs defined in Figure S3G. For each compartment, transcripts were categorized as target (high localization: log_2_ fold enrichment > 70th percentile and > 0) or background (low localization: log_2_ fold enrichment < 30th percentile and < 0). HOMER was used to compare the target and background sets, searching for enriched RNA motifs of 5–7 nucleotides. We then parsed the results, calculating enrichment *P* values, fold enrichment (%target/%background), and target prevalence (%target) for each identified motif.

High-confidence motifs were selected according to the following criteria: *P* value < 1e−5, log_2_ fold enrichment > 1.5, and %target > 5.0%.

#### Linear regression and random forest using sequence lengths

To evaluate the predictive power of basic RNA length features, we employed linear regression and random forest regression. The input features for these models were the lengths of the full transcript, 5′ UTR, CDS, and 3′ UTR. The target variables were the log_2_ fold enrichment for the 14 organelles, which were standardized by subtracting the mean and scaling to unit variance. Models were trained using the predefined training set; in all machine learning tasks, we consistently used the data (a total of 3209 valid sequences) partitioned into training (70%, 2249 sequences), validation (15%, 480 sequences), and testing (15%, 480 sequences) sets using stratified random sampling. For the random forest model, 100 estimators (n_estimators=100) were used with no maximum depth restriction. Model performance was evaluated according to Pearson’s correlation coefficient between the predicted and true log_2_ fold enrichment in the held-out test set for each organelle.

#### Residual convolutional neural network (CNN) approach

As a baseline for nucleotide sequence-based machine learning, we implemented a residual convolutional neural network (ResNet-CNN). Full transcript sequences were first converted into one-hot encoded representations (A, C, G, U channels) and padded or truncated to a fixed maximum length (set as 15000 nt). The CNN architecture comprised an initial convolutional layer, followed by batch normalization and ReLU activation, succeeded by a series of residual blocks, each containing two convolutional layers, batch normalization, ReLU, dropout, and a shortcut connection to facilitate gradient flow. Key architectural hyperparameters, including the number of filters, kernel sizes, strides (for downsampling), and dilation rates within these blocks, were varied during tuning. Following the residual blocks, a global average pooling layer was applied to condense the sequence dimension into a fixed-size feature vector irrespective of the effective sequence length after convolution. This pooled representation was then processed by a final feed-forward network consisting of a hidden layer with ReLU and dropout, connected to a linear output layer predicting the 14 continuous log_2_ fold enrichments.

The Pearson’s correlation coefficient shown in Figure 5B was for the model that achieved the highest average Pearson’s correlation coefficient across the organelles. This specific architecture used an initial convolutional layer with 32 filters and a kernel size of 7 (stride 1), followed by two residual blocks. The first residual block maintained 32 input channels and output 64 channels using a kernel size of 7 (stride 1, dilation 1). The second block received 64 input channels and output 128 channels using a kernel size of 5 (stride 2, dilation 1). A dropout rate of 0.3 was applied within the residual blocks and before the final output layer. The final feed-forward network utilized a hidden layer with 128 units.

#### Per-nucleotide embedding generation for attention model analysis

We utilized the pretrained RiNALMo language model ^145^ (giga-v1, 650 M parameters) accessed via the RiNALMo pretrained library (https://github.com/lbcb-sci/RiNALMo) to generate feature representations for RNA sequences. Input sequences corresponding to different segments (5′ UTR, CDS, 3′ UTR, or full sequence [5′ UTR+CDS+3′ UTR]) were used. The tokenized sequences were processed by the frozen RiNALMo model to obtain per-nucleotide embeddings from the final transformer layer. This resulted in a tensor of shape (sequence length × embedding dimension 1280) for each input RNA sequence/segment. A similar procedure was conducted for AIDO.RNA 1.6B (embedding dimension 2048) ^144^. For the Evo 2 DNA model, we used intermediate embeddings as suggested (layer_name = ’blocks.28.mlp.l3’, embedding dimension 4096) ^146^. For AIDO.RNA and Evo 2, we removed sequences longer than 7200 nucleotides (28 sequences in total, <1%) as these sequences produced memory errors during the embedding calculation.

#### Mean-pooling feed-forward network (FNN) approach

We also evaluated a baseline model using mean-pooled embeddings. For this approach, we utilized the per-nucleotide embeddings corresponding to a specific transcript region (5′ UTR, CDS, 3′ UTR, or full sequence). For each transcript within the selected region, a single fixed-size feature vector was generated by calculating the element-wise mean across all nucleotide embeddings in that sequence. This mean-pooled vector (with dimension 1280) served as the direct input to the regressor.

The regressor itself was a standard feed-forward network (FNN). Its architecture consisted of an input layer matching the pooled embedding dimension, followed by a configurable number of hidden layers employing ReLU activations and dropout for regularization. The final linear layer mapped the features from the last hidden layer to the 14 target compartment outputs.

To evaluate the performance of different RNA language models (RiNALMo, AIDO.RNA, and Evo 2), their respective embeddings were used as inputs for this mean-pooling FNN. The performance, as assessed by Pearson’s correlation between the predicted and experimental log_2_ fold enrichment from a held-out test set, is presented in Figure S6F. For these comparisons, an FNN architecture with three hidden layers (containing 512, 256, and 128 units, respectively) was selected, as this configuration yielded the highest average correlation across various input settings. All three embedding types demonstrated strong predictive performance, though RiNALMo and AIDO.RNA embeddings achieved slightly higher correlations than did Evo 2 embeddings.

#### Interpretable attention regressor model (TransLocAttn)

We developed a feed-forward neural network (FNN) model to predict continuous log_2_ fold enrichment across 14 subcellular compartments using precomputed, per-nucleotide RiNALMo embeddings (embedding dimension d_embed=1280). Input representations for each transcript were created by concatenating embeddings derived from its 5′ UTR, CDS, and 3′ UTR regions. To manage the dimensionality of these sequence embeddings, an optional linear projection layer with ReLU activation and layer normalization was implemented at the input stage, reducing the embedding dimension to a specified size (*e.g.*, d_proj=128 or d_proj=256) before attention processing.

The core of the model utilized an attention pooling module inspired by DeepLoc

2.0 ^224^. Instead of a single attention vector, this module learns K (*e.g.*, K=4) distinct attention (query) vectors for each of the 14 target compartments. For a given compartment, each of its K attention vectors is used to independently calculate a set of raw attention scores via dot product with the (potentially projected) per-nucleotide embeddings. These K sets of scores are then individually smoothed along the sequence using a 1D Gaussian filter (sigma=2). Subsequently, K sets of normalized attention weights for that compartment are generated by applying a softmax function across the sequence length to each smoothed score set. Each of these K attention weight sets is then used to compute a compartment-specific, vector-specific pooled representation via a weighted average of the (projected) input nucleotide embeddings. Finally, these K pooled representations for the compartment are averaged to yield a single, robust pooled vector of dimension d_proj for that compartment. The interpretable attention weights for each compartment, used for analysis, are the average of its K corresponding normalized attention weight sets.

This process results in 14 final compartment-specific pooled vectors. Each of these 14 vectors was then fed into its own dedicated FNN prediction head. Each head consisted of typically two or three hidden layers featuring ReLU activation and dropout, culminating in a single linear output neuron predicting the log_2_ fold enrichment for that specific compartment. The entire model, including the optional input projection, the multiple attention vectors per task, and the separate FNN heads, was trained end-to-end. For the final reported results (*e.g.*, Figure 5B), we utilize the model (we call it TransLocAttn) with RiNALMo embeddings, K=4 attention vectors per compartment, and an input projection layer reducing the dimension to d_proj=128. The independent FNN head for each compartment in this example configuration could comprise two hidden layers with 128 and 64 units, respectively. Training hyperparameters were: learning rate of 5e−5, dropout of 0.5, and weight decay of 1e−3.

#### Attention-based region identification and comparison with eCLIP-Seq

Following model training, learned attention weights across test-set transcripts were extracted from the best-performing configuration. To facilitate interpretation, these values, which indicate the positional contribution of each nucleotide to the pooled compartment representation rather than direct nucleotide-level effect sizes, were converted into sequence-resolved profiles for each compartment (Figures 5D-F and S6L-M).

To assess whether these inferred attention profiles aligned with known protein– RNA interaction sites, we compared them qualitatively against compiled eCLIP-Seq datasets ^137^ for representative genes. For these visualizations, attention profiles were smoothed using a Gaussian filter (σ = 5 nt) and filtered for eCLIP peaks with a signal threshold of at least 10 (see “eCLIP-Seq data processing” for details).

#### Model validation

The post-hoc analyses presented in Figure S6G-I and S6K, along with the accompanying partition controls, were performed across five random seeds (17, 42, 83, 101, and 137) on the fixed, gene-disjoint publication split. To construct the matched protein baseline, annotated CDS amino acid sequences were embedded using ESM-2 (esm2_t33_650M_UR50D) ^147^. Representations from layer 33 were mean-pooled; sequences longer than 1,022 residues were segmented into consecutive, non-overlapping windows and aggregated via length-weighted averaging. Out-of-fold ESM-2 predictions were obtained through five-fold cross-fitting on the training set, defining the target residuals as the observed compartment enrichment minus the corresponding ESM-2 prediction. Feed-forward architectures identical to those in Figure S6F were then trained on mean-pooled RiNALMo embeddings (UTR, CDS, or full transcript) to predict these residual values. Feature scaling was determined exclusively from training folds, and UTR/CDS models included gene-shuffled controls generated by permuting RNA representations within each split. The evaluation endpoint was the mean difference in test-set Pearson correlation across the 14 targets between the combined model (ESM-2 plus RNA residual model) and the baseline ESM-2 model. Statistical significance was assessed using target-block bootstrap confidence intervals and one-sided target-block sign-flip tests, adjusted across the three empirical contrasts via the Benjamini–Hochberg method. Figure S6G displays the raw target-wise Pearson correlations, and Figure S6H shows the corresponding differences.

Split controls are reported directly within the text. Sequence clusters generated with MMseqs2 for transcripts and proteins were grouped into connected components and partitioned into 70/15/15 train/validation/test sets without dividing any component. A companion model was trained on a random split matched for cluster density and partition size. Both models were evaluated on the identical 483-gene test set. Across the five seeds, the mean test-set Pearson correlation over the 14 compartments was 0.580 for the homology-aware partition compared to 0.582 for the density-matched random split.

Position-resolved analyses utilized frozen, base-level RiNALMo embeddings within the TransLocAttn framework (AdamW optimizer, learning rate 0.00005, weight decay 0.001, dropout 0.5, batch size 16, up to 100 epochs, early stopping patience 10) to predict the residual targets. A capacity-matched mean-pooling model and a gene-shuffled UTR baseline served as prespecified controls behind a frozen predictive gate. Transcripts were segmented into non-overlapping windows, 50 nt for UTRs and frame-aligned 51 nt for CDS regions (up to 60 windows per gene), and ranked by their compartment-specific attention weights independently of eCLIP data.

In silico mutagenesis was conducted on held-out genes where the top predicted compartment matched the observed label. For each gene, the top-ranked window supporting five valid modifications was compared against a bottom-quintile control window within the same gene and region, selected to minimize differences in GC content and relative transcript position. UTR windows received five deterministic permutations preserving dinucleotide frequencies; CDS windows received five deterministic synonymous recodings that preserved the translated amino acid sequence and ESM-2 inputs. Each perturbed sequence was re-embedded with the full RiNALMo model and evaluated across all five frozen models. The gene-level effect was calculated as the mean absolute shift in residual output across edits and seeds, with the overall contrast reported as the median ratio of high- to low-attention effects alongside bootstrap intervals. Significance testing for high-versus-low differences used a one-sided gene-level sign-flip test (99,999 iterations with a plus-one correction), with joint Benjamini–Hochberg adjustment across UTR and CDS sets (adjusted q = 4×10^−5^ for UTRs; q = 5×10^−5^ for CDS). For scale normalization, gene-level effects were also standardized by the sample standard deviation of observed APEX-Ribo targets across all 474 held-out genes.

#### eCLIP-Seq data processing

For analysis of the eCLIP Seq datasets ^137^, the BAM files aligned to hg38 were obtained from the ENCODE portal (https://www.encodeproject.org). The 5′ ends of the reads corresponding to each region (5′ UTR, CDS, and 3′ UTR, based on the

GENCODE Human release 44) were counted. The eCLIP-Seq enrichment was then calculated using DESeq2 (ver. 1.32.0) as the fold change in IP samples relative to size-matched input samples in each region. Only mRNAs with a read count of at least one were included in the analysis. The mean eCLIP enrichment was calculated for each RBP–cell line combination from mRNAs positively enriched in the local translation atlas.

#### Spatial comparison of attention with eCLIP

For comparison with eCLIP (Figure S6K), genes were selected based solely on observed and predicted compartment values. eCLIP tracks with maximum signal below

10 were excluded. Tracks and eCLIP signal were Gaussian-smoothed (σ = 5 nt), converted to ranks, and z-scored within the 5′ UTR, 3′ UTR, or CDS, and five-seed tracks were rank-transformed within each region prior to seed averaging; spatial correspondence for each gene–target–experiment association was the nucleotide-level Spearman correlation of the concatenated region-standardized tracks.

The null distribution circularly shifted each track 199 times independently within the same regions. For group-level inference, correlations were collapsed to a median across experiments within gene and target, and then to a median across genes, and compared with the matched null medians as a one-sided empirical *P* value with a plus-one correction. UTR and CDS *P* values were jointly adjusted using the Benjamini– Hochberg procedure, yielding an adjusted q of 0.02 in both region sets; Figure S6K shows the corresponding per-gene correlations. All eligible eCLIP experiments for a gene were included in the analysis; experiments were not selected based on the RNA-binding protein. Association-level tests, which evaluated the two-sided absolute correlation against the same null, left no individual gene–target–experiment association below the adjusted threshold at 199 shifts.

#### Enrichment calculation for neuronal compartments

mRNAs with read counts ≥ 1 in both APEX-Ribo-Seq replicates of Nefl-pulldown, Map2-pulldown, or Snca-pulldown (14766 genes in total) were selected for downstream analyses. The enrichment was calculated as pulldown samples normalized by each input, *i.e.*, target pulldown/input. mRNAs with log_2_ fold enrichment > 0 and adjusted *P* value < 0.05 were defined as locally translated genes.

The following neuron-specific transcriptome and translatome datasets were compared with our data: RNA-Seq data from microdissected neuropils ^154^, Ribo-Seq data from microdissected neuropils ^53^, TRAP-Seq data from CA1 neuropils ^155^, and dendritic PL-Ribo-Seq data ^63^.

#### Enrichment calculation for germ granules in fly embryos

Transcripts with read counts ≥ 10 in at least three of the four samples (two APEX-pulldown and two input replicates) at both 0–1 h and 2–3 h AEL were retained, yielding a list of 13,734 transcripts used for all downstream analyses. The enrichment was calculated as pulldown samples normalized by each input by DESeq2 (ver. 1.32.0). Transcript-level results were collapsed to gene level by stripping isoform suffixes and retaining, for each gene symbol, the isoform with the smallest adjusted *P* value, yielding 4,814 genes. mRNAs with log_2_ fold enrichment > 1 and adjusted *P* value < 0.01 were defined as germ granule-localized genes.

