## Supplementary material for "Sequence features and dynamics of subcellular translation revealed by APEX-Ribo-Seq": Table S7

| Antibodies | | |
| --- | --- | --- |
| Rabbit monoclonal anti-V5 Epitope Tag Alexa Fluor 488 | Bio-Techne | Cat#FAB8926G; RRID:AB_2938590 |
| Mouse monoclonal anti-TOMM20 (F-10) Alexa Fluor 647 | Santa Cruz Biotechnology | Cat#sc-17764 AF647; RRID:AB_628381 |
| Rabbit monoclonal anti-CANX [EPR3632] Alexa Fluor 647 | Abcam | Cat#ab225062; RRID:N/A |
| Rabbit polyclonal anti-Catalase | Abcam | Cat#ab16731; RRID:AB_302482 |
| Rabbit monoclonal anti-AP2S1 | Proteintech | Cat#84174-3-RR; RRID:AB_3671731 |
| Mouse monoclonal anti-TGN46 | Proteintech | Cat#66477-1-Ig; RRID:AB_2881843 |
| Rabbit monoclonal anti-GM130 (D6B1) | Cell Signaling Technology | Cat#12480; RRID:AB_2797933 |
| Rabbit polyclonal anti-EEA1 | MBL | Cat#PM062; RRID:AB_10598350 |
| Rabbit polyclonal anti-GFAP | Proteintech | Cat#16825-1-AP; RRID:AB_2109646 |
| Rabbit polyclonal anti-RUFY1 | Proteintech | Cat#13498-1-AP; RRID:AB_2183747 |
| Rabbit monoclonal anti-PCNT | Sigma-Aldrich | Cat# ZRB1318;  RRID:N/A |
| Mouse monoclonal anti-TUBB3 | Proteintech | Cat#66375-1-Ig; RRID:AB_2814998 |
| Rabbit polyclonal anti-SYN1 | Proteintech | Cat#20258-1-AP; RRID:AB_2800493 |
| Alpaca monoclonal anti-mouse IgG1 Alexa Fluor 568 | Proteintech | Cat#sms1AF568-1-10; RRID:AB_2827579 |
| Rabbit polyclonal anti-CAST | Proteintech | Cat#12250-1-AP; RRID:AB_2071702 |
| Rabbit polyclonal anti-TMEM38B | Proteintech | Cat#19919-1-AP; RRID:AB_2878623 |
| Rabbit polyclonal anti-UBE2J2 | Proteintech | Cat#17713-1-AP; RRID:AB_2210754 |
| Rabbit polyclonal anti-KDSR | Proteintech | Cat#16228-1-AP; RRID:AB_2249648 |
| Rabbit polyclonal anti-ECHS1 | Proteintech | Cat#11305-1-AP; RRID:AB_2262166 |
| Rabbit polyclonal anti-MRPL55 | Proteintech | Cat#17679-1-AP; RRID:AB_2878427 |
| Rabbit polyclonal anti-CROCC | Novus Biologicals | Cat#NBP1-80820; RRID:AB_11019491 |
| Rabbit polyclonal anti-DCTN1 | Proteintech | Cat#55182-1-AP; RRID:AB_10837074 |
| Rabbit polyclonal anti-KRT19 | Proteintech | Cat#14965-1-AP; RRID:AB_2133324 |
| Rabbit polyclonal anti-TRIP11 | Proteintech | Cat#26456-1-AP; RRID:AB_2880519 |
| Rabbit polyclonal anti-GORASP1 | Proteintech | Cat#10747-2-AP; RRID:AB_2113204 |
| Rabbit polyclonal anti-FAR2 | Proteintech | Cat#27717-1-AP; RRID:AB_3669618 |
| Rabbit monoclonal anti-phospho-eIF2α (Ser51) (D9G8) | Cell Signaling Technology | Cat#3398; RRID:AB_2096481 |
| Rabbit monoclonal anti-eIF2α (D7D3) | Cell Signaling Technology | Cat#5324; RRID:AB_10692650 |
| Rabbit polyclonal anti-RPS17 | Abcam | Cat#ab138991; RRID:AB_2687572 |
| Rabbit monoclonal anti-puromycin | Abcam | Cat#ab315887; RRID:AB_3716527 |
| Mouse monoclonal anti-puromycin | Sigma-Aldrich | Cat#ZMS1016; RRID:AB_3099686 |
| Goat polyclonal anti-rabbit IgG IRDye 680 | LI-COR | Cat#926-68071; RRID:AB_10956166 |
| Rabbit Monoclonal V5-Tag (D3H8Q) Antibody | Cell Signaling Technology | Cat# 13202; RRID: AB_2687461 |
| Donkey anti-Rabbit IgG (H+L) Highly Cross-Adsorbed Secondary Antibody, Alexa Fluor 488 | Thermo Fisher Scientific | Cat# A-21206; RRID:AB_ 2535792 |
